# Why most Principal Component Analyses (PCA) in population genetic studies are wrong

**DOI:** 10.1101/2021.04.11.439381

**Authors:** Eran Elhaik

## Abstract

Principal Component Analysis (PCA) is a multivariate analysis that allows reduction of the complexity of datasets while preserving data covariance and visualizing the information on colorful scatterplots, ideally with only a minimal loss of information. PCA applications are extensively used as the foremost analyses in population genetics and related fields (e.g., animal and plant or medical genetics), implemented in well-cited packages like EIGENSOFT and PLINK. PCA outcomes are used to shape study design, identify, and characterize individuals and populations, and draw historical and ethnobiological conclusions on origins, evolution, dispersion, and relatedness. The replicability crisis in science has prompted us to evaluate whether PCA results are reliable, robust, and replicable. We employed an intuitive color-based model alongside human population data for eleven common test cases. We demonstrate that PCA results are artifacts of the data and that they can be easily manipulated to generate desired outcomes. PCA results may not be reliable, robust, or replicable as the field assumes. Our findings raise concerns about the validity of results reported in the literature of population genetics and related fields that place a disproportionate reliance upon PCA outcomes and the insights derived from them. We conclude that PCA may have a biasing role in genetic investigations. An alternative mixed-admixture population genetic model is discussed.

---

> “Would you tell me, please, which way I ought to go from here?”
>
> “That depends a good deal on where you want to get to,” said the Cat.
>
> Lewis Carroll, Alice in Wonderland 1865, Macmillan Publishers

## Introduction

The ongoing reproducibility crisis, undermining the foundation of science (Baker 2016), raises various concerns ranging from study design to statistical rigor (Ioannidis 2005; Krafczyk et al. 2021). Population genetics is confounded by its utilization of small sample sizes, ignoring effect sizes, and adopting questionable study designs. The field is relatively small and may involve financial interests (Lee et al. 2009; Kaiser 2015; Stokstad 2019) and ethical dilemmas (Pennisi 2005; Holmes 2018). Since biases in the field rapidly propagate to related disciplines like medical genetics, biogeography, association studies (Hannon et al. 2016), forensics, and paleogenomics in humans and non-humans alike, it is imperative to ask whether and to what extent our most elementary tools satisfy risk criteria.

Principal Component Analysis (PCA) is a multivariate analysis that reduces the data’s dimensionality while preserving their covariance. When applied to genotype data, typically encoded as AA, AB, and BB (where A and B are two alleles), PCA finds the eigenvalues and eigenvectors of the covariance matrix of allele frequencies. The data are reduced to a small number of dimensions or principal components (PCs); each describes a decreased proportion of the genomic variation. Individual genotypes are then projected onto space spanned by the PC axes, which allows visualizing the samples and their distances from one another in a colorful scatter plot. In this visualization, sample overlap is considered a reflection of common ancestry (Patterson, Price, and Reich 2006; Price et al. 2006). PCA’s most attractive property for population geneticists is that the distances between clusters allegedly reflect the genetic and geographic distances between them. PCA also supports the projection of points onto the components calculated by a different dataset, potentially accounting for insufficient data in the projected dataset. Initially adapted for human genomic data in 1963 (Edwards and Cavalli-Sforza 1963), the popularity of PCA has increased over time. Still, it was not until the release of the SmartPCA tool (EIGENSOFT package) (Price et al. 2006) that PCA was propelled to the front stage of population genetics.

PCA is used as the first analysis of data investigation and data description in most population genetic analyses (e.g., Atzmon et al. 2010; Behar et al. 2010; Campbell et al. 2012; Lazaridis et al. 2016). It has a wide range of applications. It is used to examine the population structure of a cohort to determine ancestry, analyze the demographic history and admixture, decide on the similarity of samples and exclude outliers, decide how to model the populations in downstream analyses, describe the relationships between the samples, infer kinship, identify clines of ancestries in the data (e.g., Patterson et al. 2010; Yang et al. 2010; Moorjani et al. 2011; Ramstetter et al. 2017), detect genomic signatures of natural selection (e.g., Duforet-Frebourg et al. 2015), and identify convergent evolution (e.g., Galinsky et al. 2016). It is used by direct-to-consumer companies to estimate ethnicity (Ball et al. 2020) and in forensics to infer ancestry and biogeography (e.g., McNevin 2020). PCA is embedded in databases like gnomAD (Karczewski et al. 2020), MR-Base (Hemani et al. 2018), and the UK Biobank (Bycroft et al. 2018), where PCA loadings are offered. PCA or PCA-like tools are considered the ‘gold standard’ in genome-wide studies (GWAS) and GWAS meta-analyses, where they are routinely used to cluster individuals with shared genetic ancestry and to detect, quantify, and adjust for population structure (e.g., Chen et al. 2017; Connolly et al. 2019; Müller et al. 2019). PCA is also used to identify cases and controls (e.g., Luca et al. 2008; Genovese et al. 2010; Willis et al. 2014; Mobuchon et al. 2017) as well as outliers (samples or data) (Moorjani et al. 2011; van’t Hof et al. 2016), assess SNP effects (Price et al. 2006), and calculate population structure covariates (Peterson et al. 2019). The demand for large sample sizes has prompted researchers to “outsource” analyses to direct-to-consumer companies that employ full discretion in their choice of tools, methods, and data – none of which are shared – and return the PCA loadings and other “summary statistics” (e.g., Ganna et al. 2019). PCA also serves as the primary tool to identify the origins of ancient samples in paleogenomics (Lazaridis et al. 2016), to identify biomarkers for forensic reconstruction in evolutionary biology (Li et al. 2020), and to localize samples (Novembre et al. 2008). As of April 2021, over 15,000 papers employed PC scatterplots to interpret genetic data, draw historical and ethnobiological conclusions, and describe the evolution of various taxa from prehistorical times to the present – no doubt Herculean tasks for any scatterplot.

PCA’s widespread use could not have been achieved without several key traits that distinguish it from other tools – all tied to the replicability crisis. PCA can be applied to any genomic dataset, whether small or large, and it always yields results. It is parameter-free and nearly assumption-free (Patterson, Price, and Reich 2006). It does not involve measures of significance, effect size evaluations, or error estimates. It is, by large, a “black box” harboring complex calculations. The proportion of explained variance of the data is the single reflection of the quality of PCA; however, there is no consensus on the number of PCs to analyze. Price et al. (2006) recommended using 10 PCs, and Patterson et al. (2006) proposed the Tracy–Widom statistic to determine the number of components. However, this statistic is highly sensitive and inflates the number of PCs. In practicality, most authors use the first two PCs, which are expected to reflect genetic similarities and are difficult to observe in higher PCs. The remaining authors use an arbitrary number of PCs or adopt *ad hoc* strategies to aid their decision (e.g., Solovieff et al. 2010). Pardiñas et al. (2018), for example, selected the first five PC “as recommended for most GWAS approaches” and principal components 6, 9, 11, 12, 13, and 19, whereas Wainschtein et al. (2019) preferred the top 280 PCs. There are no proper usage guidelines for PCA, and “innovations” towards less restrictive usage are adopted quickly. Recently, even the practice of displaying the proportion of variation explained by each PC faded as those proportions dwarfed (e.g., Lazaridis et al. 2016). Since PCA is affected by the markers, samples, the precise implementation, and various flags implemented in the PCA packages – each has an unpredictable effect on the results – replication cannot be expected.

In population genetics, PCA and admixture-like analyses are the *de-facto* standards used as non-parametric genetic data descriptors. They are considered the hammer and chisel of genetic analyses (Elhaik 2016). Lawson et al. (2018) commented on the misuse of admixture-like tools and argued that they should not be used to draw historical conclusions. Thus far, no investigation has thoroughly explored PCA usage and accuracy across most common study designs.

Because PCA fulfills many of the risk-criteria for reproducibility (Ioannidis 2005) and due to its centrality as a first hypothesis generator in population genetic studies, this study is set to assess its reliability, robustness, and reproducibility. As PCA is a mathematical model employed to describe the unknown truth, testing its accuracy requires a convincing model where the truth is unambiguous. For that, we developed an intuitive and simple color-based model (Figure 1A).

**Figure 1.**
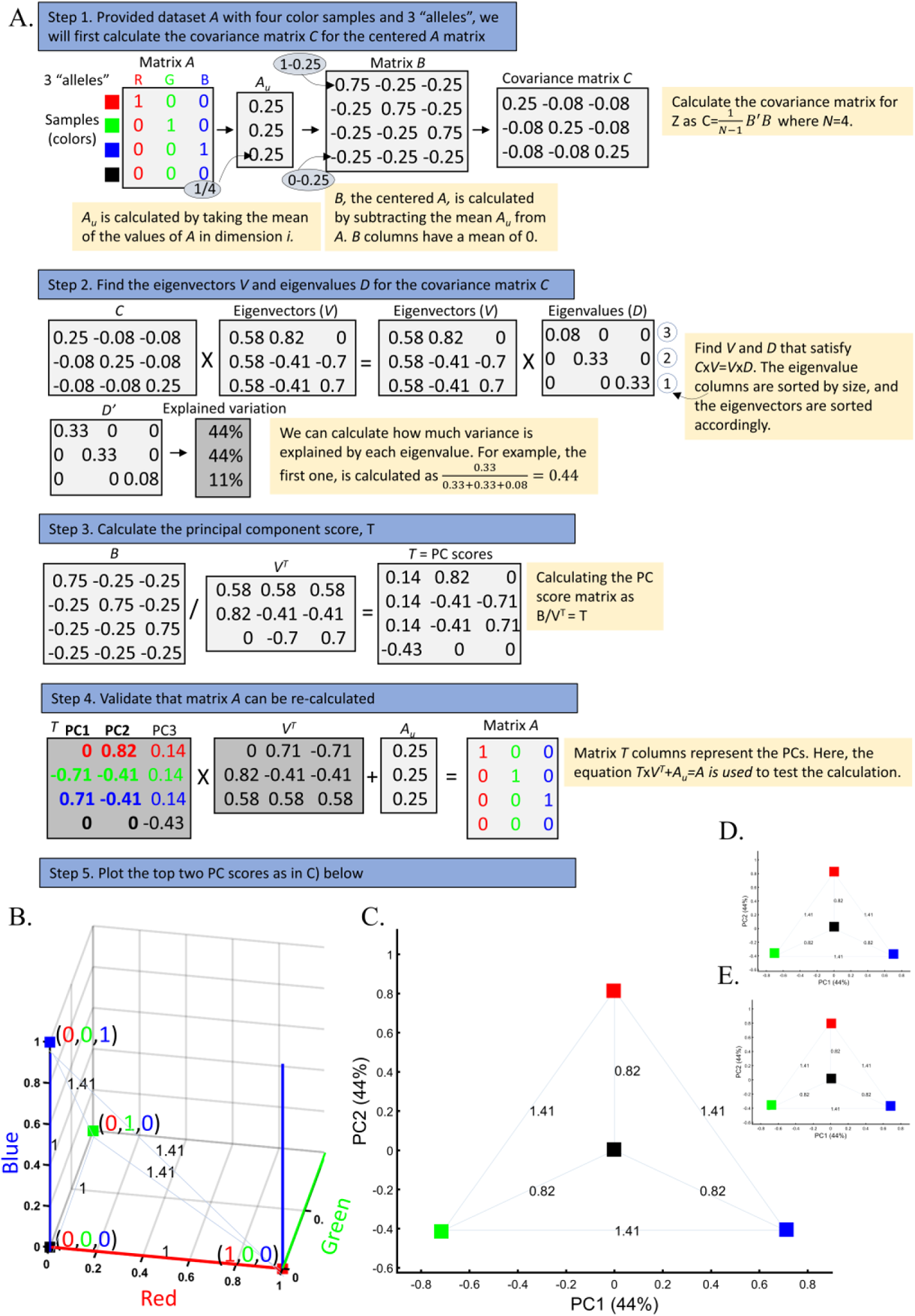
Applying PCA to four color populations. A) An illustration of the PCA procedure (using the singular value decomposition (SVD) approach) applied to a color dataset consisting of four colors (*N_All_*=1). B) A 3D plot of the original color dataset with the axes representing the primary colors. The color populations (in their true color) are then plotted along with their two top PCs with C) *N_All_*=1, D) *N_All_*=100, and E) *N_All_*=10,000. The latter two results are identical to those of C). Grey lines and labels mark the Euclidean distances between the color populations calculated across all three PCs.

Because all colors consist of three components—red, green, and blue (analogous to SNPs)—they can be plotted in a 3D plot representing this truth (Figure 1B). Applied to these data, PCA reduces the dataset to two dimensions that explain most of the variation. This allows us to visualize the results in a 2D plot and measure the difference from the original 3D plot. Let us agree that if PCA cannot perform well in this simplistic setting, it should not be used in more complex analyses and certainly cannot be used to derive far-reaching conclusions about history.

We carried out an extensive empirical evaluation of PCA through eleven test cases; each assesses a common usage of PCA using color and human genomic data. The PCA version used here yields near-identical results to the PCA implemented in EIGENSOFT (Figure S1-S2). Some of the results for the color populations are summarized in boxes and illustrate realistic investigation scenarios using common terminology to the field. If PCA results are irreproducible, if they can be manipulated, directed, or controlled by the experimenter, if multiple and conflicting results can be generated from the same dataset, or if PCA produces absurd results, then PCA must not be used for genetic investigations, and an incalculable number of findings based on its results should be reevaluated. We found that this is indeed the case.

## Results

### 1. The near-perfect case of dimensionality reduction

Applying principal component analysis (PCA) to a dataset of four even-sized populations: the three primary colors (Red, Green, and Blue) and Black illustrates a near-ideal dimension reduction example. PCA condensed the dataset of these four samples from a 3D Euclidean space (Figure 1B) into three principal components (PCs), the first two of which explained 88% of the variation (Figure 1C). Here, and in all other color-based analyses, the colors represent the true 3D structure, whereas the 2D plots are the outcome of PCA. Although PCA correctly positioned the primary colors in even distances from each other and Black, it distorted the distances between the primary colors and Black (from 1 in 3D space to 0.82 in 2D space). Thereby, even in this limited and near-perfect demonstration of data reduction, the observed distances do not reflect the actual distances between the samples (which are impossible to recreate in a 2D dataset). Evenly increasing all the sample sizes yields identical results irrespective of the sample size (Figure 1D-E). In other words, the number of samples does not affect the plot’s topography if the increase is proportional to all the populations.

When analyzing human populations, which harbor most of the genomic variation between continental populations (12%) with only 1% of the genetic variation distributed within continental populations (Elhaik 2012), PCA tends to position Africans, Europeans, and East Asians along the three edges of an imaginary triangle, which closely resembles our color-population model and illustration. Analyzing continental populations, we obtained similar results for two even-sized population datasets (Figures 2A, 2C) and their quadrupled counterparts (Figures 2B, 2D). As before, the distances between the populations remain similar (Figures 2A-B and 2C-D), demonstrating that for populations of the same size, sample size does not contribute to the distortion of the results if the increase in size is proportional.

**Figure 2.**
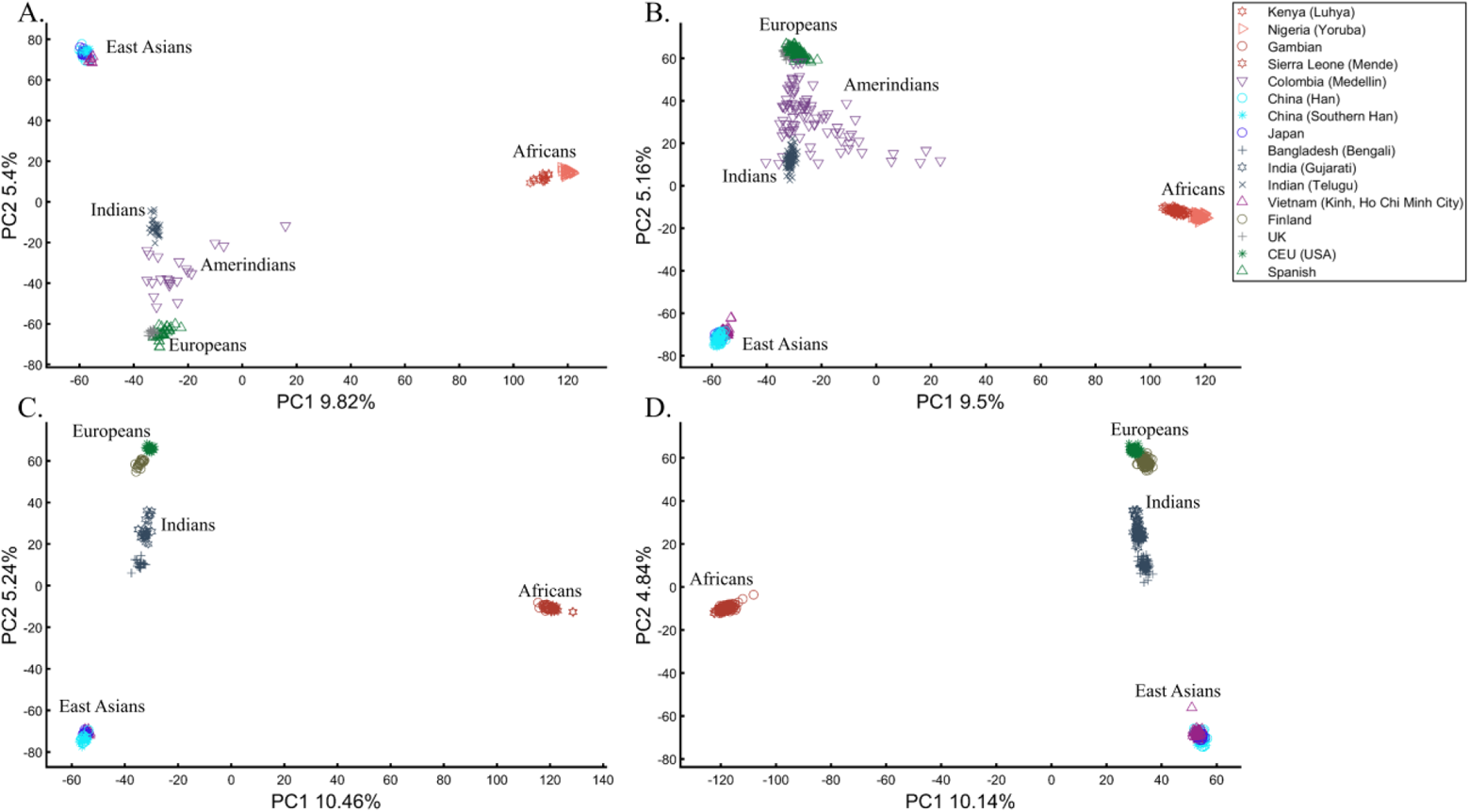
Testing the effect of sample sizes on even-sized populations. The top plots show nine populations with *n*=50 (A) and *n*=188 (B). The bottom plots show a different set of nine populations with *n*=50 (C) and *n*=192 (D). In both cases, increasing the sample size did not alter the PCs (the y-axis flip between C and D is a known phenomenon).

### 2. The case of populations of different size

The extent to which different-sized populations produce vastly different results with conflicting interpretations is illustrated through a typical study case in Box 2.

#### Box 2

**Studying the origin of Black using the primary colors**

Three research groups sought to study the origin of Black. A previous study that employed even sample-sized color populations alluded that Black is a mixture of all colors (Figure 1B-D). A follow-up study with a larger sample size (*N_Red_*=*N_Green_*=*N_Blue_*=10) and enriched in Black samples (*N_Black_*=200) (Figure 3A) reached the same conclusion. However, the Black-is-Blue group suspected that the Blue population was mixed. After QC procedures, it reduced its sample size, which decreased the distance between Black and Blue and supported their speculation that Black has a Blue origin (Figure 3B). The Black-is-Red group hypothesized that the underrepresentation of Green, compared to its actual population size, masks the Red origin of Black. They comprehensively sampled the Green population and showed that Black is very close to Red (Figure 3C). Another Black-is-Red group contributed to the debate by genotyping more Red samples. To reduce the bias from other color populations, they kept the Blue and Green sample sizes even. Their results replicated the previous finding that Black is closer to Red and thereby shares a common origin with it (Figure 3D). A new Black-is-Green group challenged those results, arguing that the small sample size and omission of Green samples biased the results. They increased the sample sizes of all the other populations of the previous study and demonstrated that Black is closer to Green (Figure 3E). The Black-is-Blue group challenged these findings on the grounds of the relatively small sample sizes that may have skewed the results and dramatically increased all the sample sizes. However, believing that they are of Purple descent, Blue refused to participate in further studies. Their relatively small cohort was explained by their isolation and small effective population size. The results of the new sampling scheme confirmed that Black is closer to Blue (Figure 3F), and the group was praised for the large sample sizes that, no doubt, captured the actual variation in nature better than the former studies.

Note that unlike in Figures 1A and 3A, where Black is in the middle, in other figures, the overrepresentation of certain “alleles” (e.g., Figure 3B) shifts Black away from (0,0). Intuitively, this can be thought of as the most common “allele” (Green in Figure 3B) repelling Black, which has three null or alternative “alleles.”

**Figure 3.**
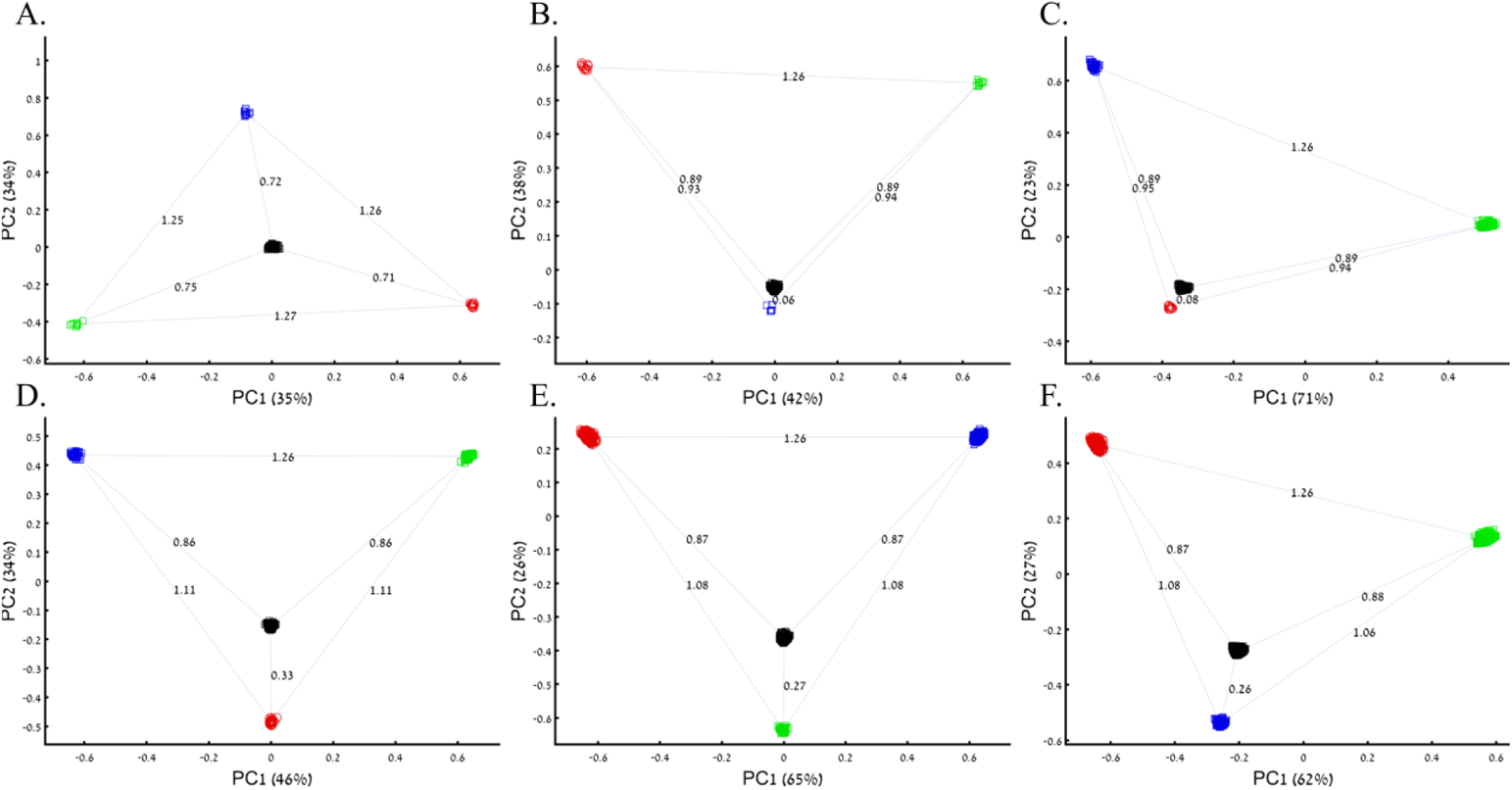
PCA of uneven-sized samples of four color populations. A) *N_Red_*=*N_Green_*=*N_Blue_*=10; *N_Black_*=200, B) *N_Red_*=*N_Green_*=10; *N_Blue_*=5; *N_Black_*=200, C) *N_Red_*=10; *N_Green_*=200; *N_Blue_*=50; *N_Black_*=200 D) *N_Red_*=25; *N_Green_*=*N_Blue_*=50; *N_Black_*=200, E) *N_Red_*=300; *N_Green_*=200; *N_Blue_*=*N_Black_*=300, and F) *N_Red_*=1000; *N_Green_*=2000; *N_Blue_*=300; *N_Black_*=2000. Scatter plots show the top two PCs. The numbers on the grey bars reflect the Euclidean distances between the color populations over all PCs.

PCA is commonly reported as yielding a stable differentiation of Africans vs. non-Africans and Europeans vs. Asians on the primary PCs (Li et al. 2008; Qin et al. 2015), which prompted prehistorical inferences of PCA results as representing the post Out Of Africa event followed by multiple migrations, differentiation, and admixture events to explain these patterns (Reich, Price, and Patterson 2008). For instance, Reich et al. (2008) claimed that the “rough cluster[ing]” of non-Africans is “about what would be expected if all non-African populations were founded by a single dispersal ‘out of Africa.’” Silva-Zolezzi et al. (2009) argued that the Zapotecos did not experience a recent admixture due to their location on the Amerindian cluster at the Asian end of the European-Asian cline.

However, variable population sizes can easily create alternative results as well as alternative “clines.” The same Mexican-American cohort can be made to appear closer to Europeans (Figure 4A) or as a European-Asian admixed group (Figure 4B). East Asians can appear on an Oceanian-African cline (Figure 4C), whereas Europeans can appear on an African-East Asian cline (Figure 4D) as an admixed group. Europeans can also appear in the middle of the plot as an admixed group of Africans-Asians-Oceanians origins (Figure 4E), and Oceanians can be shown to cluster with (Figure 4F) or without East Asians (Figure 4E). The latter depiction maximizes the proportion of explained variance, which common wisdom would consider to be the correct explanation. According to some of these results, only Europeans and Oceanians (Figure 4C) or East Asians and Oceanians (Figure 4D) experienced the Out of Africa event. By contrast, East Asians (Figure 4C) and Europeans (Figure 4D) may have remained in Africa.

**Figure 4.**
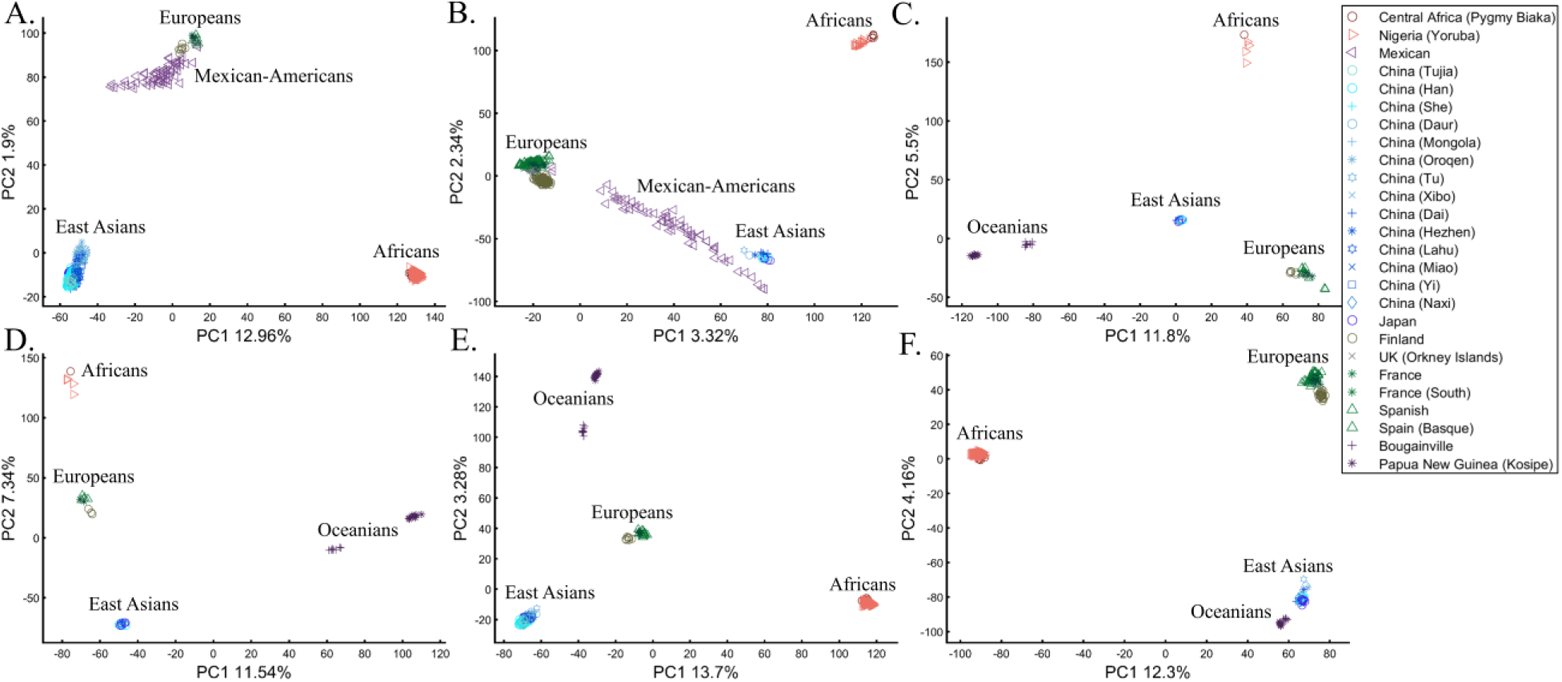
PCA of uneven-sized African (Af), European (Eu), Asian (As), Amerindian (Am), and Oceanian (Oc) populations. Fixing the sample size of Mexican-Americans and altering the sample sizes of other populations: (A) *N_Af_*=198; *N_Eu_*=20; *N_As_*=483; *N_Am_*=64 and (B) *N_Af_*=20; *N_Eu_*=343; *N_As_*=20; *N_Am_*=64) changes the results. An even more dramatic change can be seen when repeating this analysis on Oceanians: (C) *N_Af_*=5; *N_Eu_*=25; *N_As_*=10; *N_Oce_*=20 and (D) *N_Afr_*=5; *N_Eu_*=10; *N_As_*=15; *N_Oc_*=20 and when altering their sample sizes: (E) *N_Af_*=98; *N_Eu_*=25; *N_As_*=150; *N_Oc_*=24 and (F) *N_Af_*=98; *N_Eu_*=83; *N_As_*=30; *N_Oc_*=15.

Contrary to Reich et al. (2008), we observed no “rough cluster” of non-Africans, but rather multiple clusters that we created using PCA, intentionally, knowingly, and purposely by carefully selecting the populations and manipulating the sample sizes to reflect different imaginary versions of human history. Unlike Reich et al. (2008), we do not believe that their example “highlights how PCA methods can provide evidence of important migration events.” Instead, our example (Figure 4) shows how PCA can be used to generate multiple and alternative scenarios, all mathematically correct but, obviously, biologically incorrect. It is, thereby, misleading to show only one solution without acknowledging the existence of other solutions, let alone while not disclosing the proportion of explained variance.

### 3. The case of one admixed population

#### Box 3

**Studying the origin of Black using the primary and one secondary (admixed) color populations**

Following criticism on the sampling scheme used to study the origin of Black (Box 2), the redoubtable Black-is-Red group genotyped Cyan. Using even sample-sized populations, they demonstrated that Black is closer to Red (*D_Black-Red_*=0.46) (Figure 5A), where *D* is the Euclidean distance between the samples over all three PCs (short distances indicate high similarity). Their findings were criticized by the Black-is-Green school on the grounds that their Cyan samples were biased and their results do not apply to the broad Black cohort. They also reckoned that the even sampling scheme favored Red because Blue is related to Cyan through shared language and customs. The Black-is-Red group responded by enriching their cohort in Cyan and Black (*N_Cyan_*, *N_Black_*=1,000) and provided even more robust evidence that Black is Red (*D_Black-Red_*=0.12) (Figure 5B). However, the Black-is-Green camp dismissed these findings. Conscious of the effects of admixture, they retained only the most homogeneous Green and Cyan samples (*N_Green_, N_Cyan_*=33), genotyped new Blue and Black (*N_Blue_, N_Black_*=400), and analyzed them with the published Red cohort (*N_Red_*=100). The Black-is-Green results supported their hypothesis that Black is Green (*D_Black-Green_*=0.27) and that Cyan shared a common origin with Blue (*D_Blue-Green_*=0.27) (Figure 5C) and should thereby be considered an admixed Blue population. Unsurprisingly, the Black-is-Red group claimed that these results were due to the under-representation of Black since when they oversampled Black, PCA supported their findings (Figure 5A). In response, the Black-is-Green school maintained even sample sizes for Cyan, Blue, and Green (*N_Blue_, N_Green_*, *N_Cyan_*=33) and enriched Black and Red (*N_Red_*, *N_Black_*=100). Not only did their results (*D_Black-Green_*=0.63 < *D_Black-Red_*=0.89) support their previous findings, but they also demonstrated that Green and Blue completely overlapped, presumably due to their shared co-ancestry, and that together with Cyan (*D_Cyan-Green_*=0.63 < *D_Cyan-Red_* =1.09) (Figure 5B, Figure 5D) they represent an antique clade. They explained that these color populations only appeared separated due to the effects of genetic drift. However, they still retained sufficient genetic information that PCA can uncover if the correct sampling scheme is used. Further analyses by the other groups contested these findings. Black was determined to be closer to Blue with Green and Cyan sharing common origins (Figure S3A) and closer to Green with Blue and Cyan sharing common origins (Figure S3B). It was also shown that Black, not Cyan, is a Blue-Red admixed group (Figure S3C) and that Black has Red-Green ancestors (Figure S3D).

In the first large-scale study of Indian population history, Reich et al. (2009) applied PCA to a cohort of Indians, Europeans, Asians, and Africans using various sample sizes that ranged from 2 (Srivastava) to 203 (YRI) samples. After applying PCA to Indians and the three continental populations to exclude “outliers” that supposedly had more African or Asian ancestries than other samples, PCA was applied again in various settings. The authors have generated a plethora of conflicting figures, none of which disclosed the proportion of explained variance along with the first four PCs that were examined. Their concluding analysis that consisted of Indians, Asians, and Europeans (their Figure 3) showed Indians at the apex with Europeans and Asians at the opposite ends. This plot was interpreted as evidence of an “ancestry that is unique to India” and an “Indian cline.” Indian groups were explained to have inherited different proportions of ancestry from “Ancestral North Indians” (ANI), related to western Eurasians, and “Ancestral South Indians” (ASI), who split from Onge. Indians have since been described using the terms ANI and ASI.

**Figure 5.**
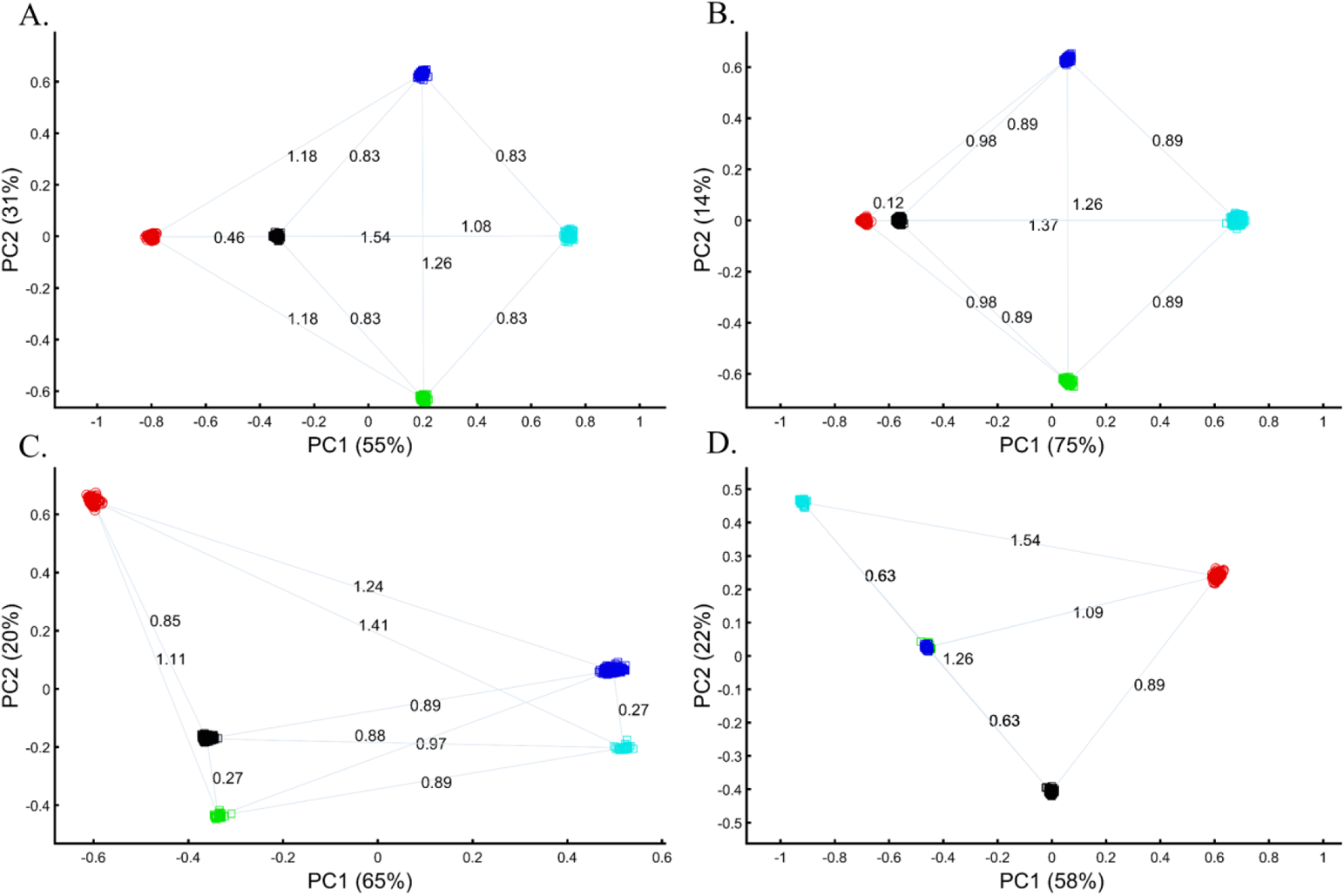
PCA with the primary and mixed color populations. A) *N_all_*=100; *N_Black_*=200, B) *N_Red_*=*N_Green_*= *N_Blue_*=100; *N_Black_*=*N_Cyan_*=500, C) *N_Red_*=100; *N_Green_*=*N_Cyan_*=33; *N_Blue_*=*N_Black_*=400; and D) *N_Red_*=*N_Black_*=100; *N_Green_*=*N_Blue_*=*N_Cyan_*=33; Scatter plots show the top two PCs. The numbers on the grey bars reflect the Euclidean distances between the color populations over all PCs.

In evaluating the claims of Reich et al. (2009) that rest on PCA, we first replicated the finding of the original “Indian cline” (Figure 6A). We then garnered support for an alternative cline using Indians, Africans, and Europeans (Figure 6B). We then demonstrated that PCA results support Indians to be European (Figure 6C), East Asians (Figure 6D), and Africans (Figure 6E), as well as a two-ways admixed European-Asian population (Figure 6F). Whereas the first two PCs of Reich et al.’s (2009) primary figure explain less than 8% of the variation (according to our Figure 6A, Reich et al.’s Figure 3 does not report this information), four out of five of our alternative depictions explain 8-14% of the variation. Our results question the authors’ choice in using an analysis that explained such a small proportion of the variation (let alone not reporting it), yielded no support for a unique ancestry to India, and cast doubt on the reliability and usefulness of the ANI-ASI model to describe Indians. It is difficult to answer whether Africa is in India or the other way around (Figure 6E). Clearly, geographical inferences based on PCA can lead to errors of Columbian magnitude.

**Figure 6.**
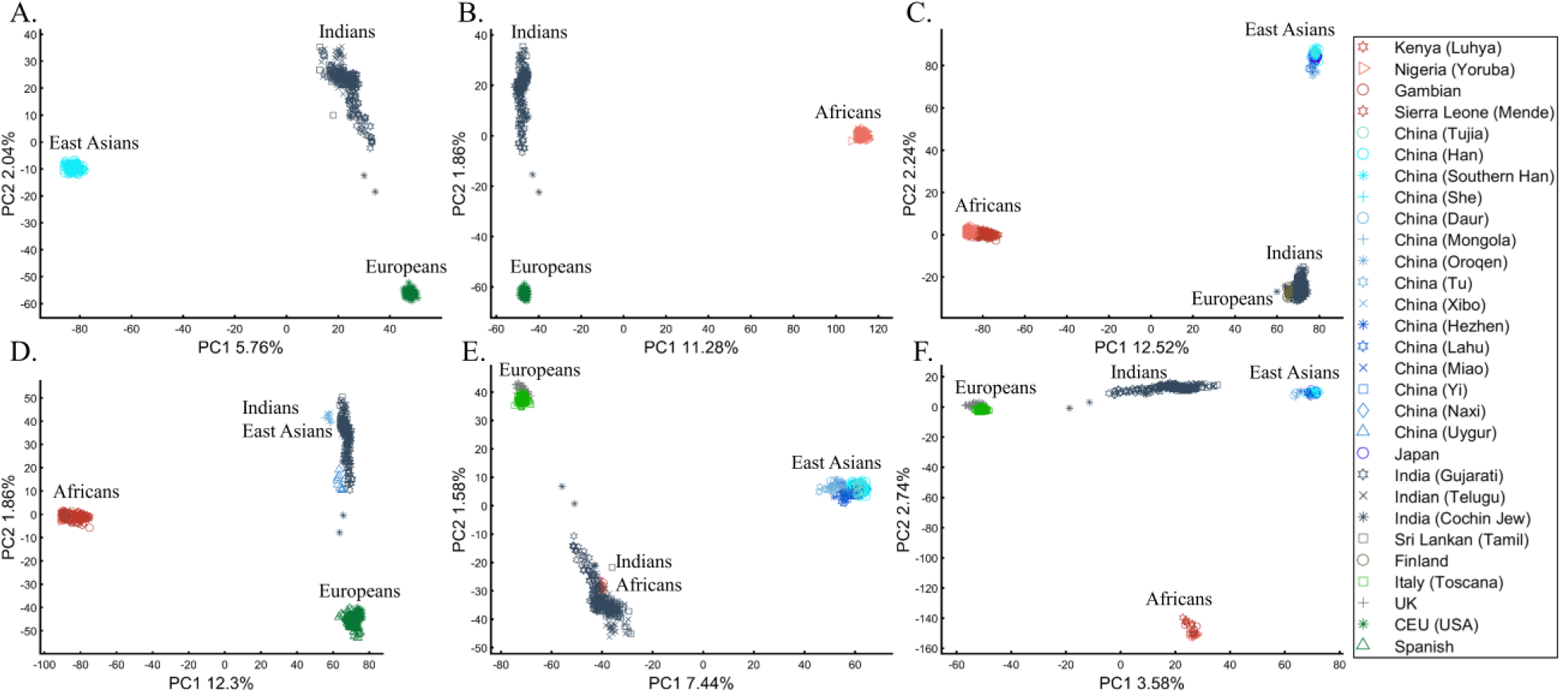
Studying the origin of Indians using PCA. A) Replicating Reich et al.’s (2009) results using *N_Eu_*=99; *N_As_*=146; *N_Ind_*=321. Generating alternative PCA scenarios using: B) *N_Af_*=178; *N_Eu_*=99; *N_Ind_*=321, C) *N_Af_*=400; *N_Eu_*=40; *N_As_*=100; *N_Ind_*=321, D) *N_Af_*=477; *N_Eu_*=253; *N_As_*=23; *N_Ind_*=321, E) *N_Af_*=25; *N_Eu_*=220; *N_As_*=490; *N_Ind_*=320, and F) *N_Af_*=30; *N_Eu_*=200; *N_As_*=50; *N_Ind_*=320.

In a separate effort, Need et al. (2009) applied PCA to 55 Ashkenazic Jews (AJs) and 507 non-Jewish Caucasians to study the origins of AJs. The PCA plot showed that Jews formed a distinct cluster from non-Jews. Based on these results, the authors suggested that PCA can be used to detect linkage to Jewishness. A follow-up PCA where Middle Eastern (Bedouin, Palestinians, and Druze) and Caucasus (Adygei) populations were included showed that AJs formed a distinct cluster that nested between the Adygei (and the European cluster) and Druze (and the Middle Eastern cluster). The authors then concluded that AJs might have mixed Middle Eastern and European ancestries. The proximity to the Adygei cluster was noted as interesting but dismissed based on the small sample size of the Adygei (*N*=17). The authors concluded that AJ genomes carry an “unambiguous signature of their Jewish heritage, and this seems more likely to be due to their specific Middle Eastern ancestry than to inbreeding.” A similar strategy was employed by Bray et al. (2010) to claim that PCA “confirmed that the AJ individuals cluster distinctly from Europeans, aligning closest to Southern European populations along with the first principal component, suggesting a more southern origin, and aligning with Central Europeans along the second, consistent with migration to this region” with other authors (Tian et al. 2008b; Tian et al. 2009) making similar claims.

It is easy to show why PCA cannot be used to reach such conclusions. We first replicated Need et al.’s (2009) primary results (Figure 7A) showing that AJs cluster separately from Europeans. However, such an outcome is typical to any non-European population like Turks (Figure 7B). It is not unique to AJs, nor does it prove that they are genetically detectable. A slightly modified design shows that most AJs overlap with Turks and allows us to promote the claim that AJs have a Turkic origin (Figure 7C). We can easily refute our conclusion by including continental populations and showing that most AJs cluster with Iberians rather than Turks (Figure 7D). This last design explains more of the variance than all the previous analyses together. This analysis questions PCA’s use as a discriminatory genetic utility.

**Figure 7.**
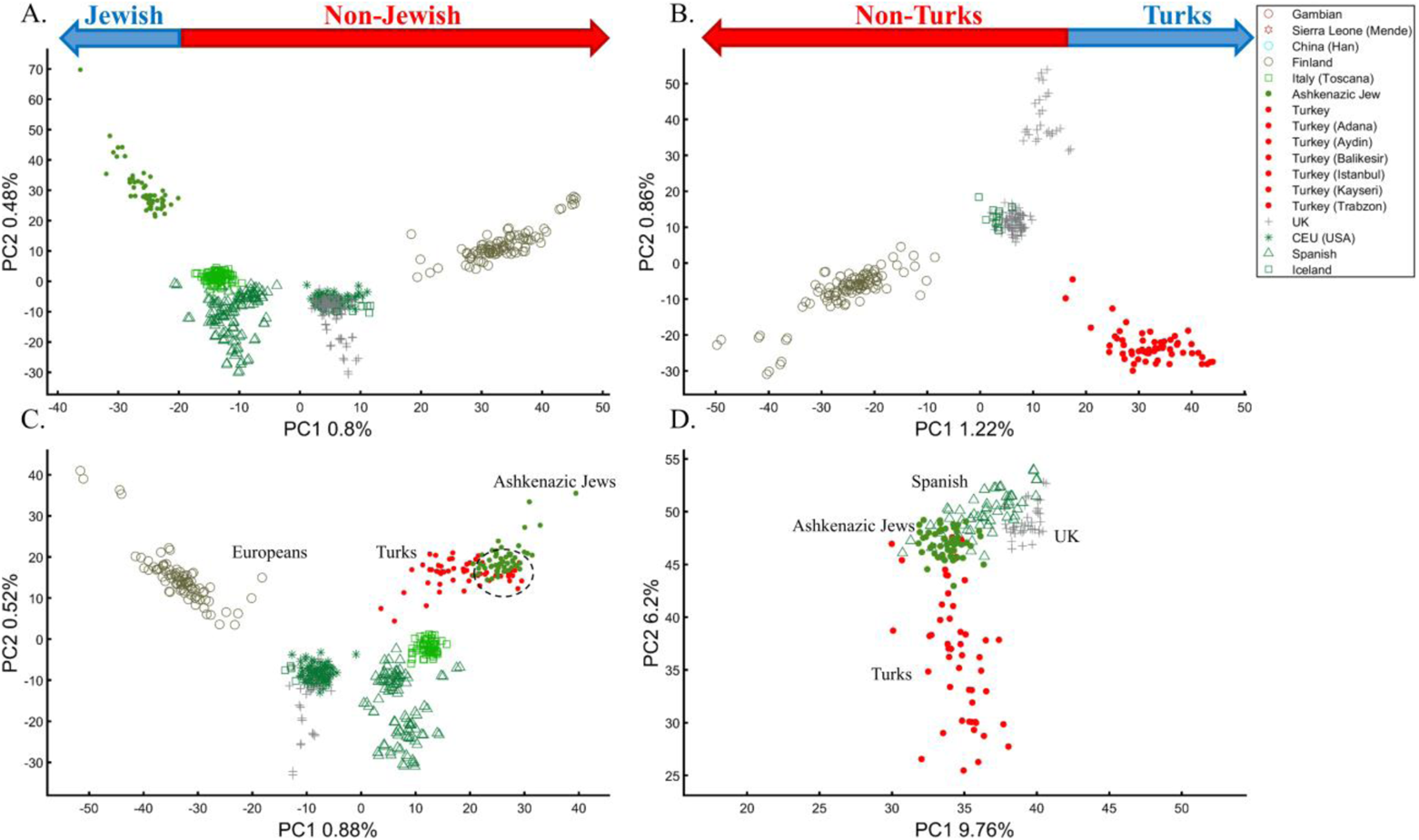
Studying the origin of 55 AJs using PCA. A) Replicating Need et al.’s results using *N_Eu_*=507; Generating alternative PCA scenarios using: B) *N_Eu_*=223; *N_Turks_*=56; C) *N_Eu_*=400; *N_Turks+Caucasus_*=56, and D) *N_Af_*=100, *N_As_*=100 (Africans and Asians are not shown), *N_Eu_*=100; and *N_Turks_*=50.

There are several more oddities with the report of Need et al. (2009). First, they did not report the variance explained by their sampling scheme (it is likely to be close to 1%, as in Figure 7A). Second, their interpretation of AJs as a mixed population is questionable, provided that most populations are nested between and within other populations, but the authors did not suggest that they are all admixed. Finally, the authors never justified their choice of populations. The conclusions of Need et al. (2009) were thereby obtained based on particular PCA schemes and preconceived ideas of AJs origins that are no more real than the Iberian origin of AJs (Figure 7D).

### 4. The case of two or three admixed population

#### Box 4

**Studying the origin of Black using the primary and secondary colors**

Though the previous analyses could not resolve the origin of Black (Box 3), there was a consensus that admixed cohorts can provide novel insights even though the issue of sampling bias remained unaddressed. An even-sample analysis (*N*=10) that included Cyan and Purple supported earlier suspicions that Black is a Green-Red admix (Figure 8A). Concerned about the low sample sizes, the Black-is-Green increased the Red, Blue, and Purple sample sizes (*N_Red_*, *N_Blue_*, *N_Purple_*=100), which both showed that Black is closer to Green and that Cyan is closer to Blue (Figure 8B). The Black-is-Blue group argued that admixed individuals should be sampled at lower numbers (*N_Cyan_*, *N_Purple_*=5) and since Blue and Green shared common origins (Figure 5D), they are interchangeable and should be sampled in an even amount to Red and Black (*N_Green_* + *N_Blue_*, *N_Red_*, *N_Black_*=50). The results showed perfect Black-Blue, Green-Cyan, and Red-Purple overlap (Figure 8C), at odds with the historical records of the origin of secondary colors. To test that, an independent group ventured to the field and collected Yellow. The results positioned Cyan and Purple close to their postulated ancestral cohorts and yielded a new genuine insight that Black is Yellow (Figure 8D). The Black-is-Red group did not dispute this claim but argued that Yellow is a recent Black admixture and that Yellow should be excluded from future analyses if we wish to understand the ancient origins of Black. The removal of Yellow from the analysis showed a complete Black-Red overlap and concrete evidence that Black has a Red origin (Figure 8E). Yet those results could not be accepted as follow-up analyses provided credence to a new novel finding proving that Black has originated from Cyan (Figure 8F).

The origin of Papuans and Bouganvilleans or Northern Melanesians (NM) has been extensively debated in the literature (e.g., McEvoy et al. 2010a; Isshiki et al. 2020). PCA yielded conflicting results not only about their closeness to other samples but to each other. When analyzed against East Asians and Oceanians, Papuans and Bouganvilleans either cluster together but separately from other samples, typically at the edge of the plot (Hudjashov et al. 2018) or separately from each other and other samples at the edges of the plot (Pugach et al. 2018). Perhaps consequently, Papuan ancestry is considered distinct from Asian ancestry, which, in turn, is considered distinct from non-Asian ancestries (Isshiki et al. 2020). However, this is not obvious from PCA studies, as Papuans were also shown to cluster with Amerindians and close to Central-Southern Asia (McEvoy et al. 2010b).

**Figure 8.**
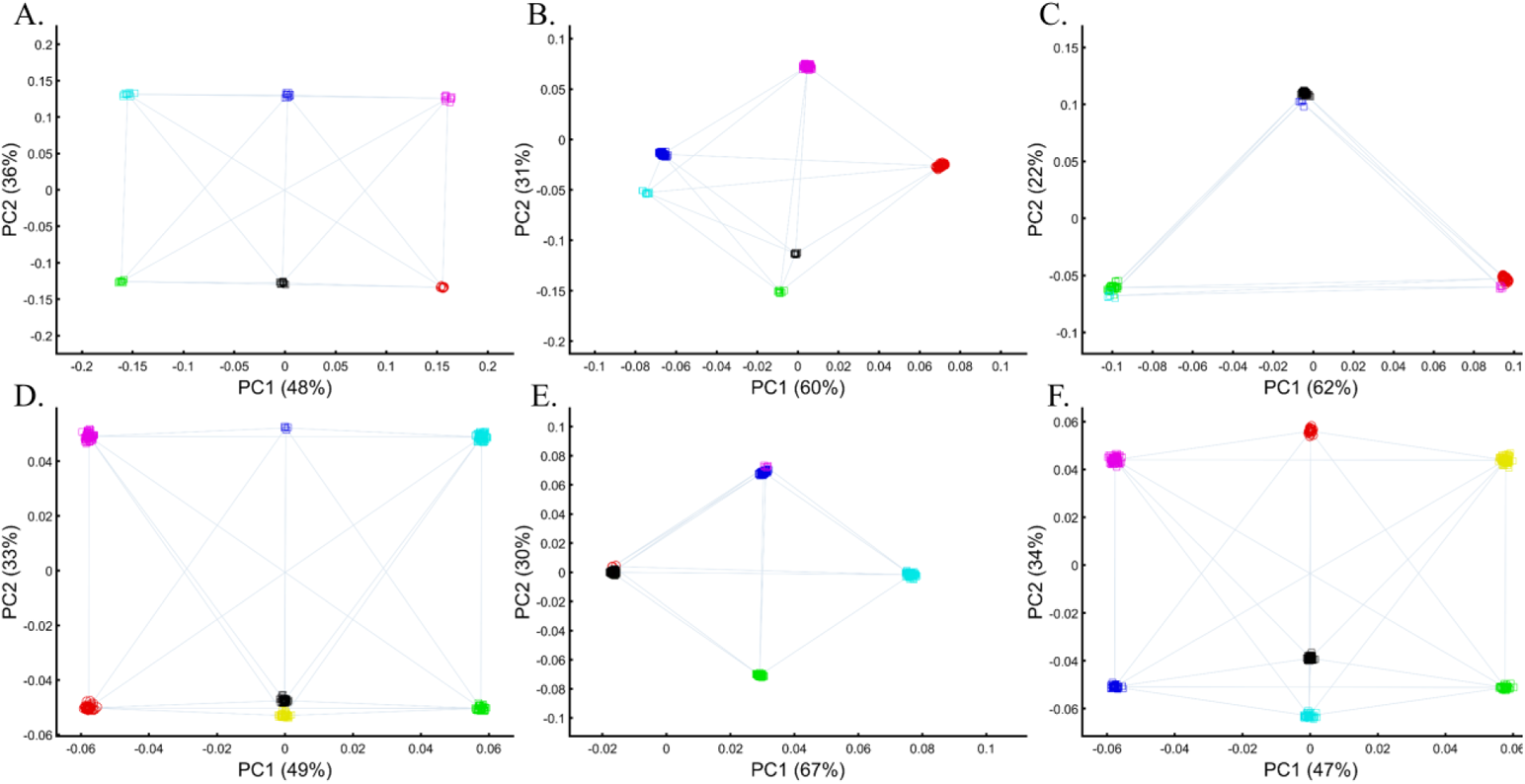
PCA with the primary and two or three mixed color populations. A) *N_all_*=100, B) *N_Red_*=*N_Blue_*=*N_Purple_*=100; *N_Green_*=*N_Black_*=*N_Cyan_*=10, C) *N_Red_*=*N_Black_*=50; *N_Blue_*=*N_Cyan_*=*N_Purple_*=5; *N_Green_*=45, D) *N_Red_*=*N_Green_*=*N_Black_*=*N_Yellow_*=50; *N_Blue_*=5; *N_Purple_*=*N_Cyan_*=100, E) *N_Black_*=800; *N_Red_*=*N_Purple_*50; *N_Blue_*=*N_Green_*=*N_Cyan_*=100. Scatter plots show the top two PCs.

We first show that when using even-sampling, the inclusion of NM creates a European-Asian cluster that was not observed before (Figure 4A) with NM clustering separately from one another at the extreme edge. Using uneven sampling, as done in all the studies, yields various contradictory results with NM clustering together and close to East Asians (Figure 4B), appearing as a three-way tri-continental admix group distinct from each other and close to Pakistani (Hazara) (Figure 4C), or remaining highly distinct and separate from other population but influencing the formation of an African-European cluster with Pakistani (Hazara) as an outgroup. Each of these results supports a different explanation for the origin of NM. Because all the results are equally mathematically valid, we can conclude that PCA produces meaningless genetic results.

**Figure 9.**
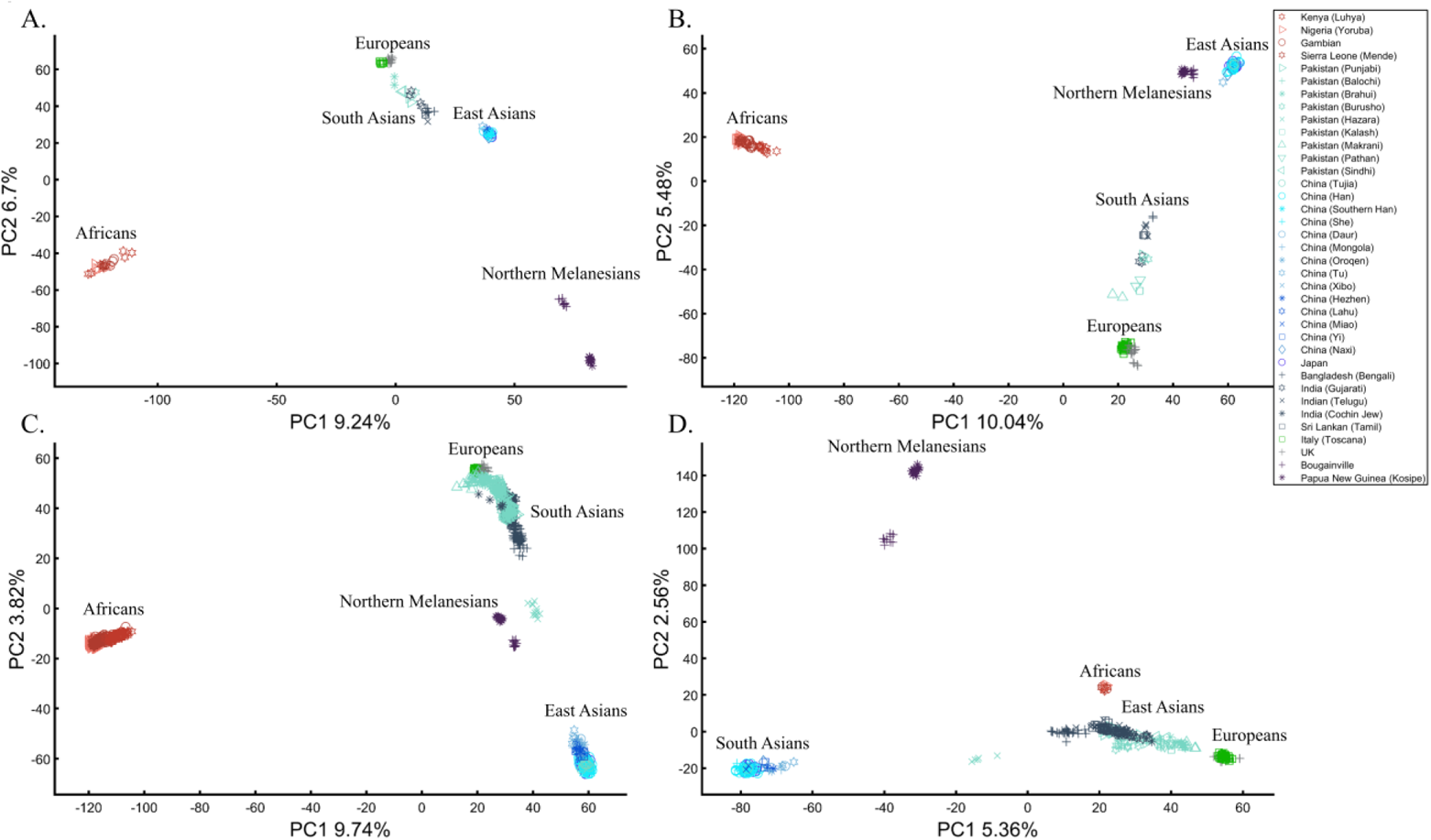
Studying the origin of Northern Melanesians using PCA. Five populations are analyzed: Africans (Af), Europeans (Eu), East Asians (EA), South Asians (SA), and Northern Melanesians (NM). Results vary based on the sample size of each population. A) *N_all_*=20, B) *N_Af_*=*N_Eu_*=*N_EA_*=50; *N_Sa_*=*N_NM_*=20, C) *N_Af_*=*N_EA_*=300; *N_Eu_*=50; *N_Sa_*=500; *N_NM_*=24, D) *N_Af_*=10; *N_Eu_*=60; *N_EA_*=100; *N_Sa_*=200; *N_NM_*=24.

### 5. The case of multiple admixed population

#### Box 5

**Studying the origin of Black using the primary and multiple mixed colors**

The value of using mixed color populations to study origins prompted new analyses using even (Figure 10A) and variable sample sizes (Figures 10B-D). Using this novel sampling scheme, the Black-is-Green school proved that Black is the closest to Green (Figure 10A), but a follow-up analysis using a different cohort yielded a novel finding that Black is closest to Pink (Figure 10B). Two final studies demonstrated that Black appears like a primary color, an ingredient of other colors (Figures 10C-D). Since Black clustered with Green and Blue, a consensus emerged that Black is the ancestor of Green and Blue, which partially explained many of the conflicting results reported thus far. That light Blue and Cyan clustered together (Figure 10D) supported previous observations (Figure 8F).

PCA has been used extensively to investigate the origins of AJs. In such analyses, it was assumed that the clustering of AJs alone is evidence of Levantine origin (e.g., Atzmon et al. 2010; Behar et al. 2010; Carmi et al. 2014) and that the “short” distance between AJs and Levantine populations indicates their close genetic relationships and thereby the Levantine origins of AJ (e.g., Behar et al. 2010). The “short” distance between AJs and Europeans was also interpreted as evidence of admixture (Atzmon et al. 2010). As a rule, the much shorter distances between AJs and the Caucasus or Turkish populations, observed by all recent studies, were ignored (Need et al. 2009; Atzmon et al. 2010; Behar et al. 2010; Bray et al. 2010; Carmi et al. 2014) in favor of Levantine of South European populations. Bray et al. (2010) concluded that not only AJs have a “more southern origin” but that their alignment with Central Europeans is “consistent with migration to this region.” In these studies, the proposition “between” received a multitude of interpretations. For example, Carmi et al.’s (2014) PCA plot that showed AJs in the extreme edge of the plot with Bedouins and French in the other edges was interpreted as AJs clustering “tightly between European and Middle Eastern populations.” The authors interpreted the lack of “outliers” (which were never defined) as evidence of common AJ ancestry.

**Figure 10.**
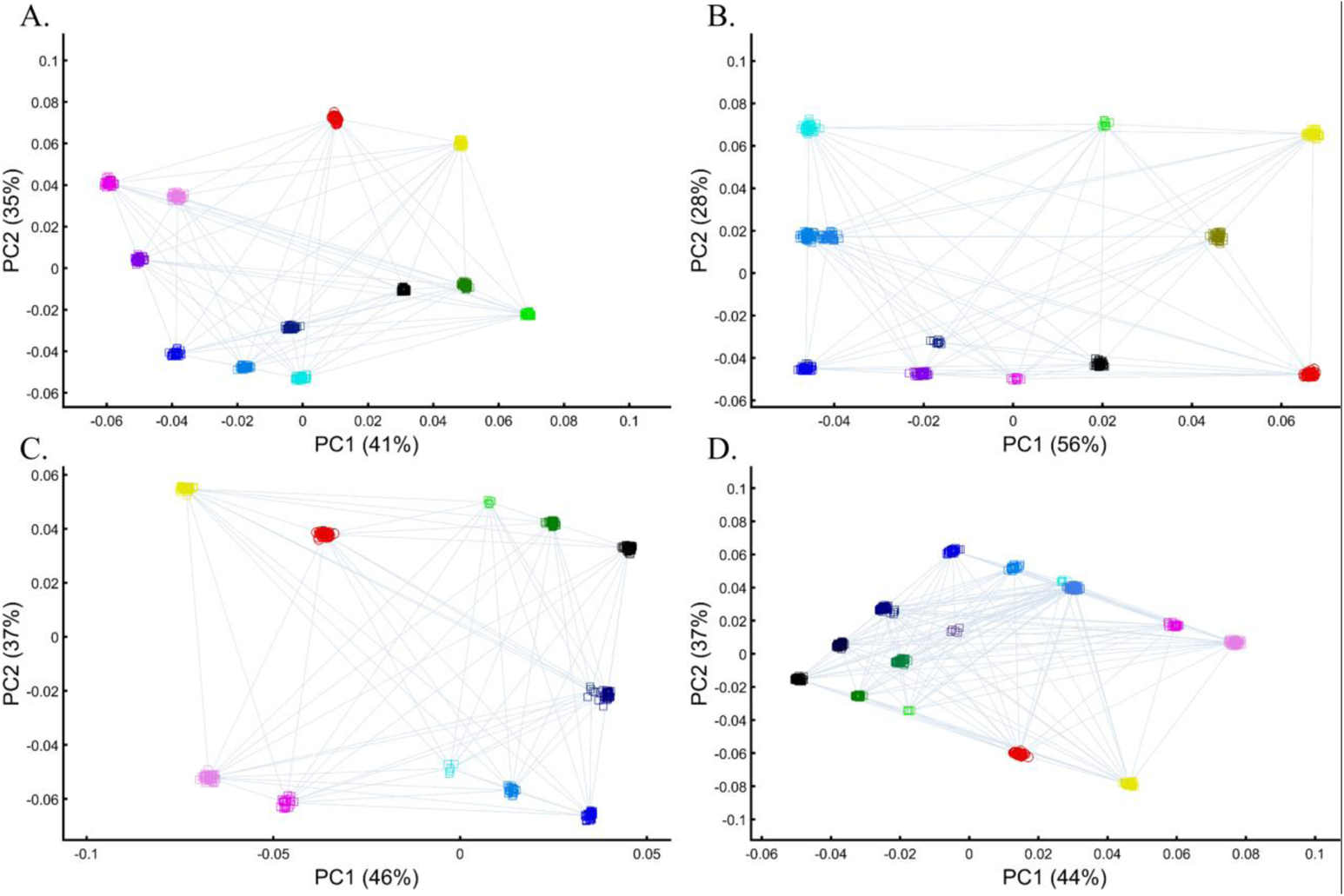
PCA with the primary and multiple mixed color populations. A) *N_all_*=50, B) *N_all_*=50 or 10, C and D) *N_All_*=[50, 5, 100, or 25]. Scatter plots show the top two PCs.

Following the rationale of these studies, it is easy to show how PCA can be orchestrated to yield a multitude of interpretations concerning the origin of AJs. We replicated the observation that AJs are “population isolate,” i.e., AJs form a distinct group, separated from all other populations (Figure 11A), and are thereby genetically distinguishable (Need et al. 2009). We also replicated the most common yet often-ignored observation in previous studies, that AJs tightly cluster with Caucasus populations (Figure 11B). We next utilized PCA to produce novel results where AJs cluster tightly with Amerindians, most likely due to the north Eurasian origins of both groups (Figure 11C). We can also show that AJs cluster much closer to South Europeans than Levantines (Figure 11D), at odds with our previous finding (Figure 11A), and overlap the Finnish samples entirely, in solid evidence of AJ’s ancient Finnish origin (Figure 11E). Last, we wish to refute our previous finding and show that only half of the AJs are of Finnish origin. The remaining have a Levantine origin (Figure 11F) – a discovery touted by all the previous reports though never actually shown. Excitingly enough, the primary PCs of this last Finnish-Levantine mixed origin depiction explained the highest amount of variance, but only a percent higher from an all-Finnish origin. We can interpret those results by the recent migration of the Finnish origin’s AJs to the Levant, where they experienced high admixture with the local Levantine populations that altered their genetic profile.

**Figure S11.**
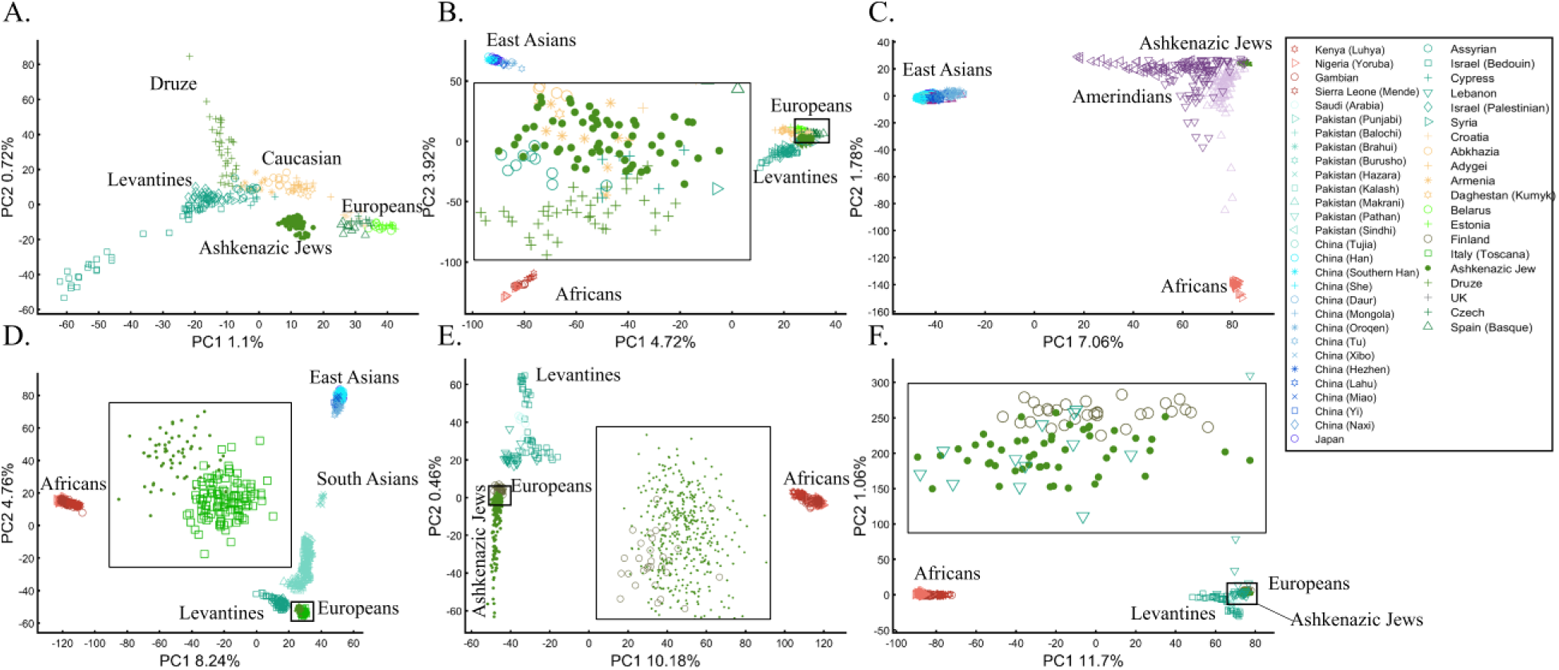
An in-depth study of the origin of AJs using PCA in relation to Africans (Af), Europeans (Eu), East Asians (Ea), Ameridians (Am), Levantines (Le), and South Asians (Sa). A) *N_Eu_*=159; *N_AJ_*=60; *N_Le_*=82, B) *N_Af_*=30; *N_Eu_*=159; *N_Ea_*=50; *N_AJ_*=60; *N_Le_*=60, C) *N_Af_*=30; *N_Ea_*=583; *N_AJ_*=60; *N_Am_*=255; D) *N_Af_*=200; *N_Eu_*=115; *N_Ea_*=200; *N_AJ_*=60; *N_Le_*=235; *N_Sa_*=88, E) *N_Af_*=200; *N_Eu_*=30; *N_AJ_*=400, *N_Le_*=80 F) *N_Af_*=200; *N_Eu_*=30; *N_AJ_*=50; *N_Le_*=160. Large square indicate insets.

### 6. The case of multiple admixed populations without “unmixed” populations

Unlike stochastic models that possess inherent randomness, PCA is a deterministic process where the model’s output is determined entirely by the parameter values and the initial conditions. This property of PCA adds to its perceived robustness. In reality, however, PCA behaves unexpectedly, where minor variations can lead to an ensemble of different outputs that appear stochastic. This effect is more substantial when populations that explain most of the variation are excluded from the analysis.

To demonstrate this effect, we show that the *same code* produces different results (Figures 12A-F) if the only variable is the standard randomization technique used to generate the individual samples of the color populations (to avoid clutter). Here, Black was the closest to Yellow (Figure 12A), Purple (Figure 12C), and Cyan (Figure 12D-E). When adding White, Black behaved as an outgroup as the distances between the secondary colors largely deviated from the expectation and produced false results (Figure 12D-F).

**Figure 12.**
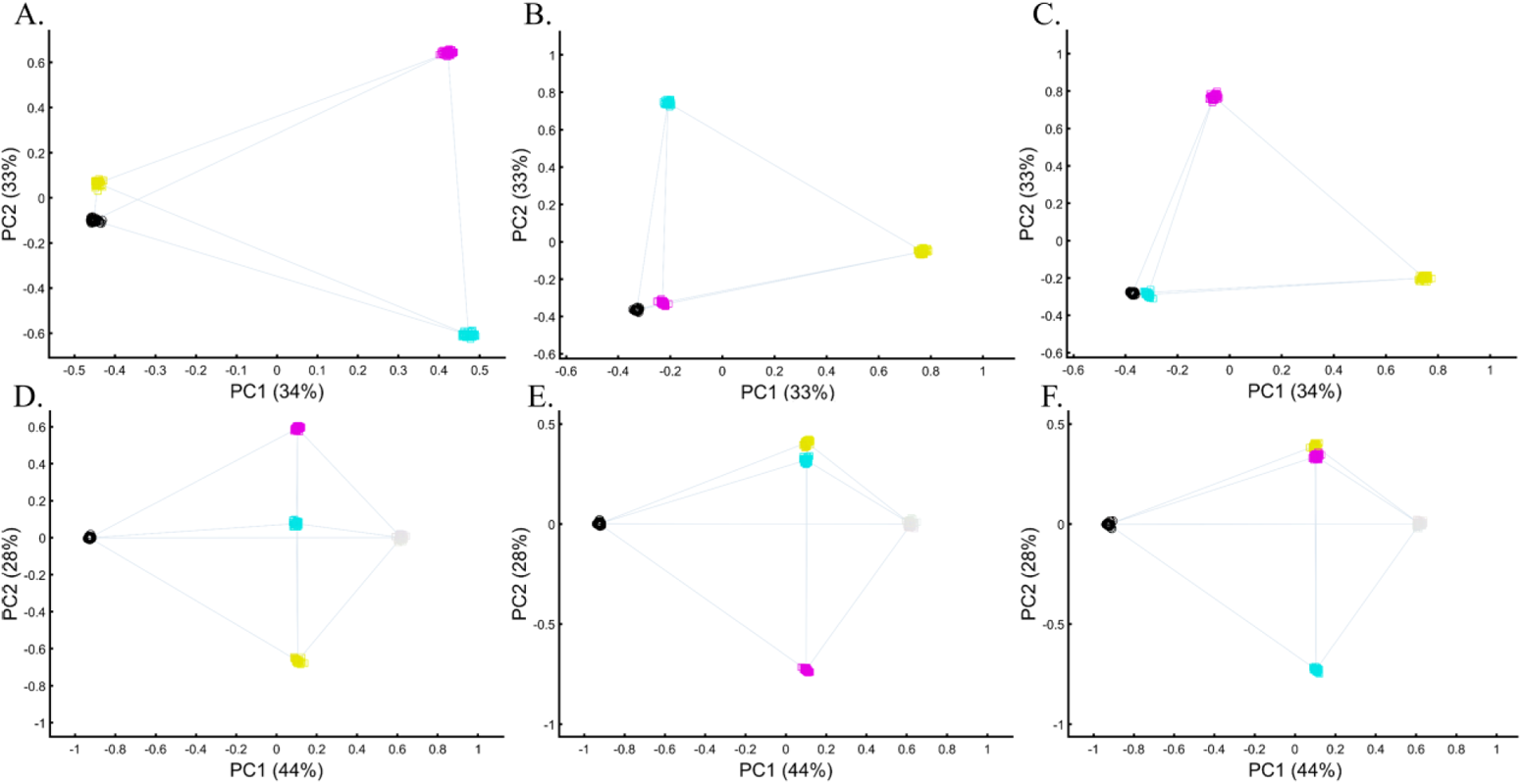
Studying the effects of minor sample variation on PCA results using color populations (*N_all_*=50). A–C) Analyzing secondary colors and Black. D–E) Analyzing secondary colors, White, and Black. Scatter plots show the top two PCs.

To demonstrate this effect on real populations, we curated a cohort of 16 populations. We carried out PCA on ten random individuals from 15 random populations. We show that these analyses result in spurious and conflicting results. Puerto Ricans, for instance, clustered close to Europeans (A), between Africans and Europeans (B), close to Adygei (C), and close to Europe and Adygei (D). Indians clustered with Mexicans (A, B, and D) or apart from them (C). Mexicans themselves cluster with (A and D) or without Africans (B and C). Papuans and Russians cluster close (B) or afar (C) from East Asian populations. More robust clustering was observed for East Asians, Caucasians, and Europeans, as well as Africans. However, these were not only indistinguishable from the less robust clustering but also failed to replicate over multiple runs (results not shown). Note that the proportion of explained variance was similar in all the analyses, which demonstrates that it is not an indication of accuracy nor robustness.

### 7. The case of pairwise comparisons

Several authors adopted a pairwise comparison scheme to assess the genetic similarity between two cohorts of interest (e.g., Need et al. 2009; Chiang et al. 2010; O’Connor et al. 2015; Rahman et al. 2018; Wang et al. 2018a). This setting is prevalent in case-control analyses that seek overlap between the compared groups (e.g., cases and controls) (e.g., Luca et al. 2008; Genovese et al. 2010; Willis et al. 2014; Mobuchon et al. 2017). However, this setting can also lead to erroneous conclusions. To demonstrate that the existence or absence of overlap in PCA in a pairwise study design is an artifact of the sampling scheme, we analyzed two non-overlapping color populations (Figure 14A) and show that in the presence of two other samples, they highly overlap (Figure 14B). Although in this analysis, the additional samples are distinct, in real-time, they may appear as part of the cohorts and alter the findings. Moreover, the latter analysis explains 99% of the variation compared to the former analysis (94%), which may appear more reliable. We next demonstrate the opposite effect by analyzing two cohorts of interest alongside other populations. Whether the two cohorts overlap or not depends on the choice of the other populations (Figure 14C-D). In other words, PCA outcomes as to whether populations overlap are independent of whether or not the populations are distinct from each other but instead are based on the sampling scheme. Misinterpreting PCA results can lead to a reduction in power and erroneous conclusions.

**Figure 14.**
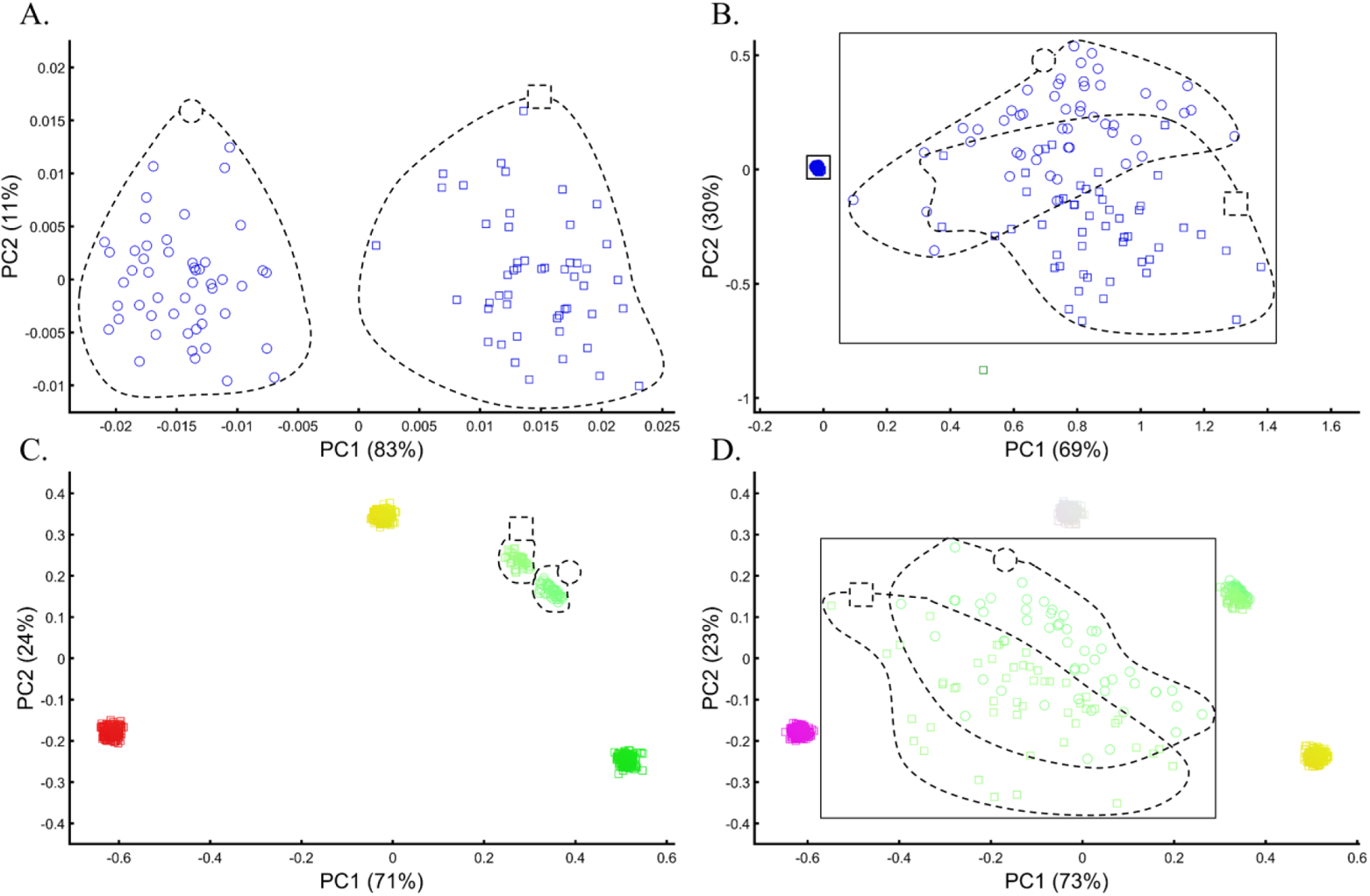
Using PCA in a pairwise setting to assess the similarity between two color-population cohorts. Even-size cohorts (*N*=50) of two distinct shades of Blue (circles and squares) do not overlap (A) and mostly overlap (B) when analyzed along with Green and White samples. Even-sized cohorts (*N*=50) of two distinct shades of Green (circles and squares) do not overlap when analyzed with three even-sized (*N*=250) populations (C) but overlap when analyzed with other even-sized (*N*=250) populations (D).

We further demonstrate that PCA produces conflicting results in real populations based on the choice of the reference populations or merely their inclusion or exclusion. In Figure 15A, we show that two Chinese populations, which PCA purports are a single homogeneous population, can be split if Japanese are included in the sampling scheme (Figure 15B). That is, PCA’s outcome for the genetic relationships between Dai and Southern Han Chinese are relative to the inclusion of another population. Likewise, PCA’s answer to whether Mexicans and Peruvians share common origins and can thereby be compared in a pairwise setting depends on the presence of Africans in the scheme (Figure 15D-D), which determines the genetic relationships between the two populations, as far as PCA is concerned. These examples show that PCA outcomes are unreliable when studying the relationships between populations in a pairwise scheme.

**Figure 15.**
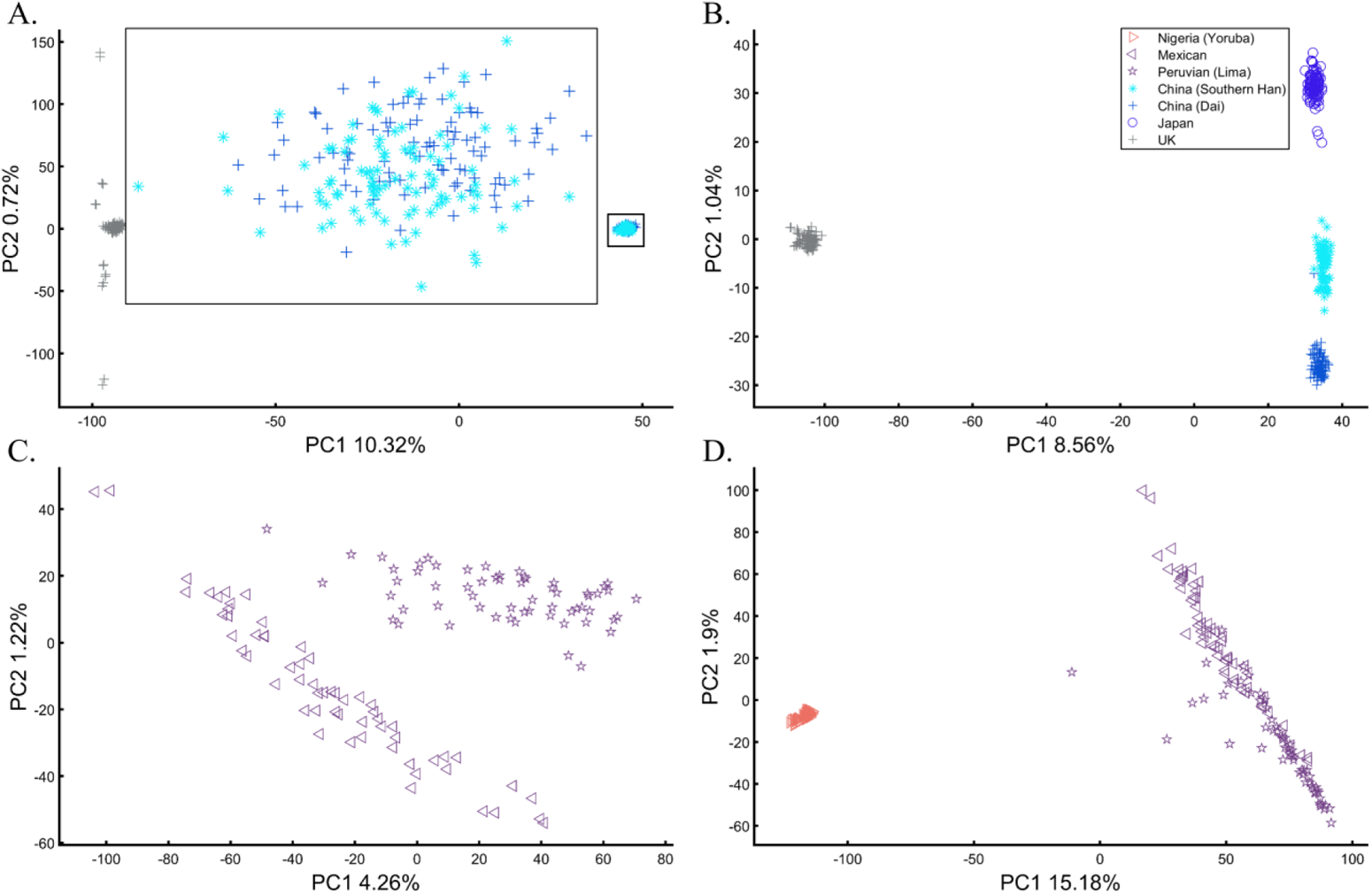
Evaluating the reliability of cohort clustering to assess their homogeneity. Southern Han and Dai Chinese appear overlapping when analyzed with UK samples (A) but completely distinct when analyzed alongside Japanese (B). PEL (Lima Peruvians) and Mexican Americans (MXL) cluster distinctively from each other (C) but completely overlap when analyzed with Africans (D). The large square indicates an inset.

### 8. The case of case-control matching and GWAS

Samples of unknown ancestry or with self-reported ancestry are typically identified by applying PCA to the cohort combined with reference populations of known origins (e.g., 1000 Genomes) (e.g., Chen et al. 2017; Connolly et al. 2019; Müller et al. 2019; e.g., Wright et al. 2019). To test whether using PCA to identify the origin of an unknown cohort with known samples is feasible, we simulated a large and heterogeneous Cyan population (Figure 16A, circles) of self-reported Blue origin. Following a typical GWAS scheme, we carried out PCA for these individuals and seven known and distinct color populations. PCA grouped the simulated individuals with Blue and even Black individuals (Figure 16B), although none of the simulated individuals are Blue nor Black (Figures 16A), as a different PCA scheme confirmed (Figure 16C). The clustering with Blue and Black is an artifact due to the choice of reference populations. A case-control assignment of this cohort to Blue or Black based on the PCA result (Figure 16B) would produce poor matches that will reduce the power of the analysis. To demonstrate that, we repeated the analysis with different reference populations (Figures 16D). Here, the simulated individuals exhibit minimal overlap with Blue and no overlap with Black and overlapped mostly with the Cyan reference population present this time.

**Figure 16.**
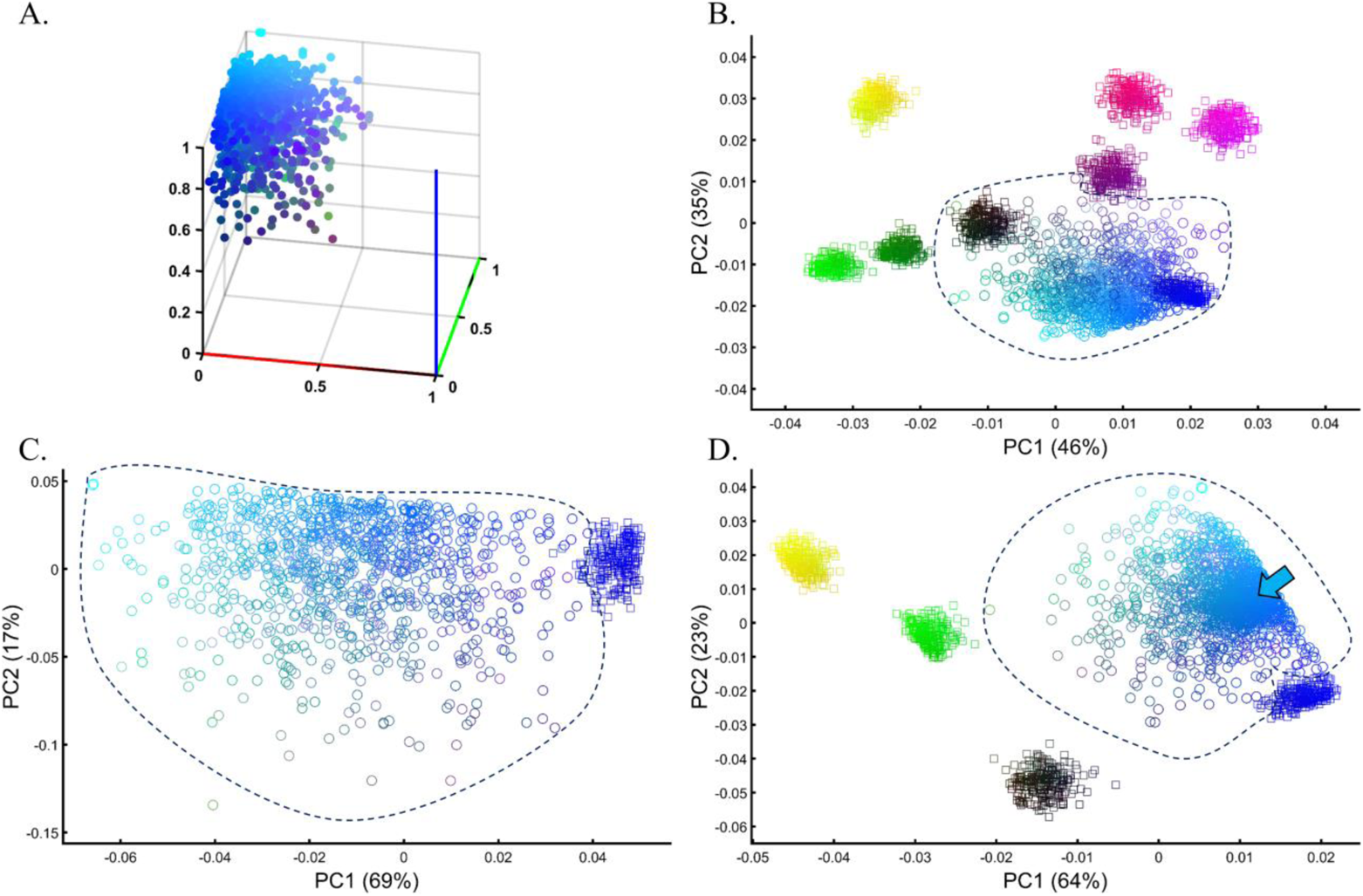
Evaluating the accuracy of PCA clustering of a heterogeneous test population in a simulation of a GWAS setting. A) The true distribution of a large heterogeneous Cyan population (*N*=1000). B) PCA of the test population with eight even-sized (*N*=250) reference populations. C) PCA of the test population with Blue from the previous analysis shows a minimal overlap between the cohorts. D) PCA of the test population with five even-sized (*N*=250) reference populations, including Cyan (marked by an arrow).

We next asked whether PCA results can be used to group Europeans into homogeneous clusters. These can be thought of as samples of both known and unknown ancestry to identify the latter. Cluster homogeneity was calculated by applying *k*-means clustering to PC1 and PC2 for *K* clusters, where *K* was the square root of the number of samples. If all the individuals in the cluster belonged to the same population, that cluster was considered homogeneous. The percent of homogeneous clusters is reported. Analyzing four European populations yielded 43% homogeneous clusters (Figure 17A). Adding Africans and Asians and then South Asian populations decreased the European cluster homogeneity to 14% and 10%, respectively (Figures 17B-C). Including the 1000 Genome populations, as customarily done, yielded 14% homogeneous clusters (Figure 17D). Although the Europeans remained the same, the addition of other continental populations resulted in a three to four times decrease in the homogeneity of their clusters.

**Figure 17.**
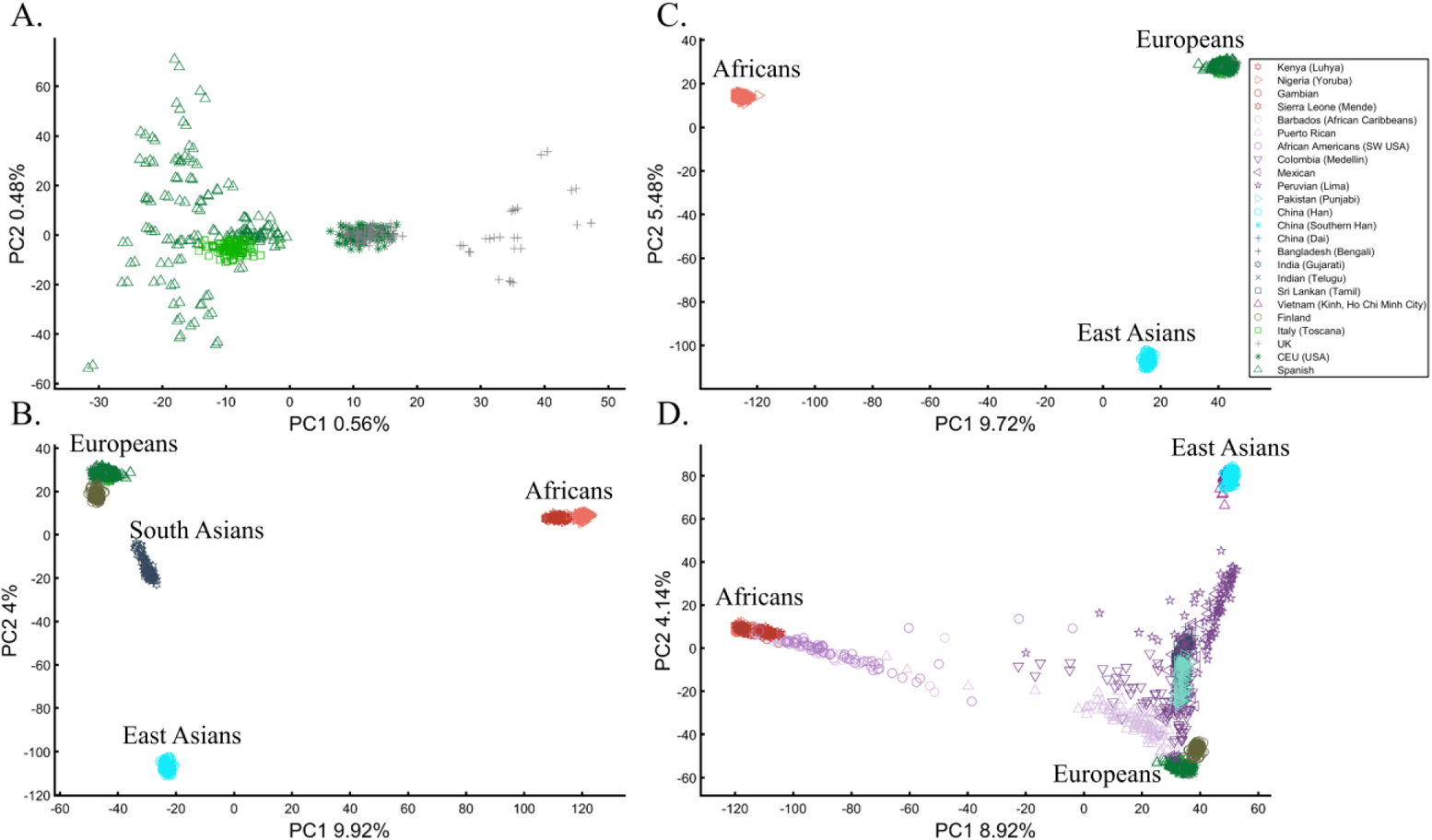
Evaluating the clustering homogeneity of European samples. PCA was applied to the four European populations (Tuscan Italians [TSI], Northern and Western Europeans from Utah [CEU], British [GBR], and Spanish [IBS]) alone (A), together with an African and Asian population (B), as well as South Asian population (C), and finally with all the 1000 Genomes Populations (D).

Since the first PCs explain a higher proportion of the variation, it is customary to assume that using more PCs can increase cluster homogeneity. This notion, however, may be supported up to a certain point. We demonstrate the effect of considering more PCs by calculating the homogeneity for different PCs for either 10 or 20 African, Asian, and European populations (Figure 17E). The maximum proportion of cluster homogeneity is reached between 10-15 PCs and fluctuates afterward. Adding more populations decreased the mean percentage of homogeneous clusters. As expected, East Asians had the highest proportion of homogeneous clusters and Africans the lowest. Note that the cluster homogeneity in this limited setting should not be confused with the amount of variance explained by additional PCs.

**Figure 17E.**
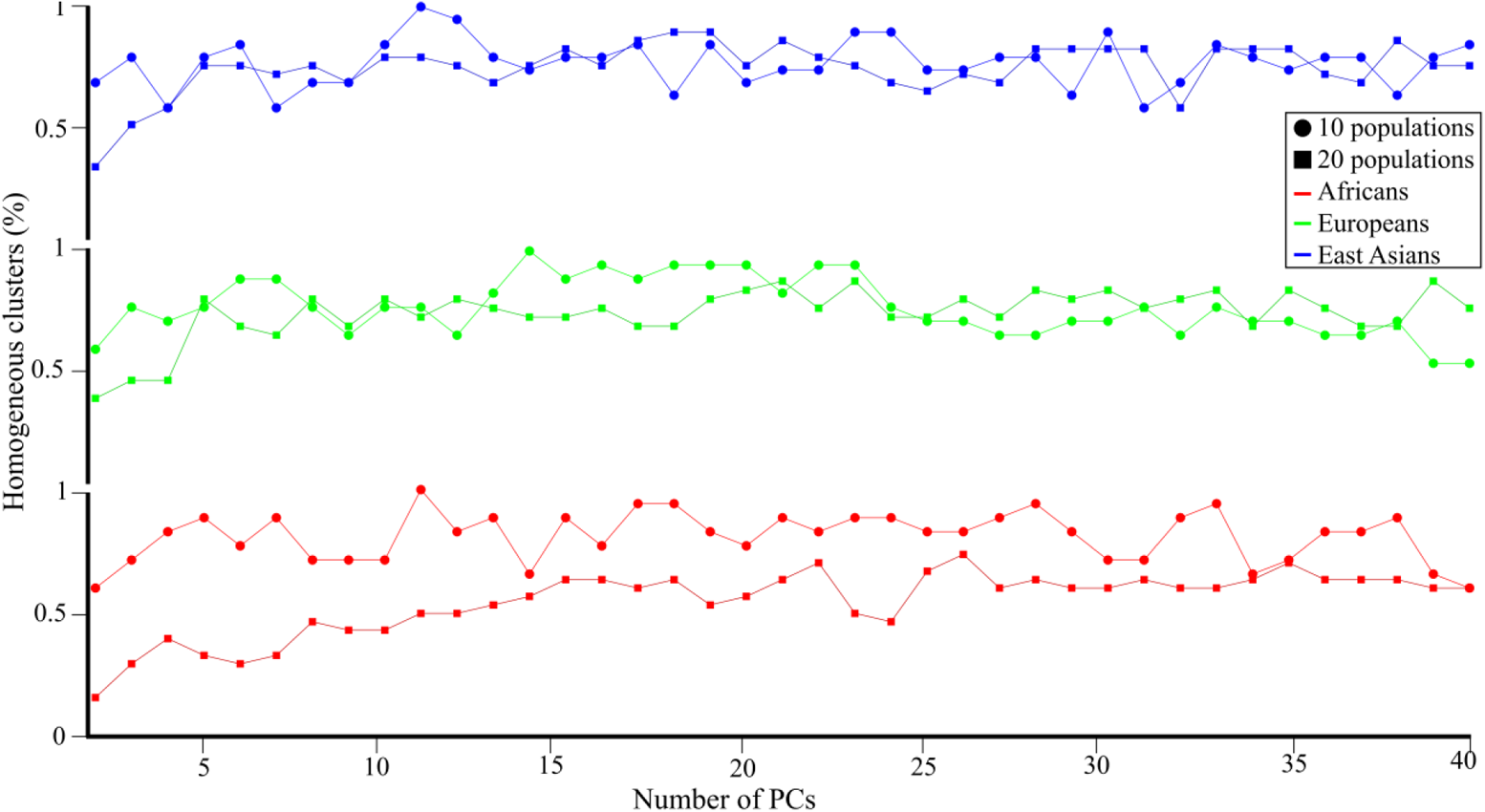
Cluster homogeneity for 10 or 20 random African, European, or Asian populations on the first 40 PCs. The proportion of homogeneous clusters in each analysis is plotted against the number of PCs.

To further demonstrate that PCA clustering does not equal shared ancestry nor biogeography, two of the most common applications of PCA, whether in GWA or population genetic studies (e.g., Chen et al. 2017), we applied PCA to 20 Puerto Ricans (Figure 18) and 300 Europeans. The Puerto Ricans clustered indistinguishably with the Europeans, whether in the first two or higher PCs (Figure 18). The Puerto Ricans represented over 6% of the cohort, sufficient to generate a stratification bias in an association study.

**Figure 18.**
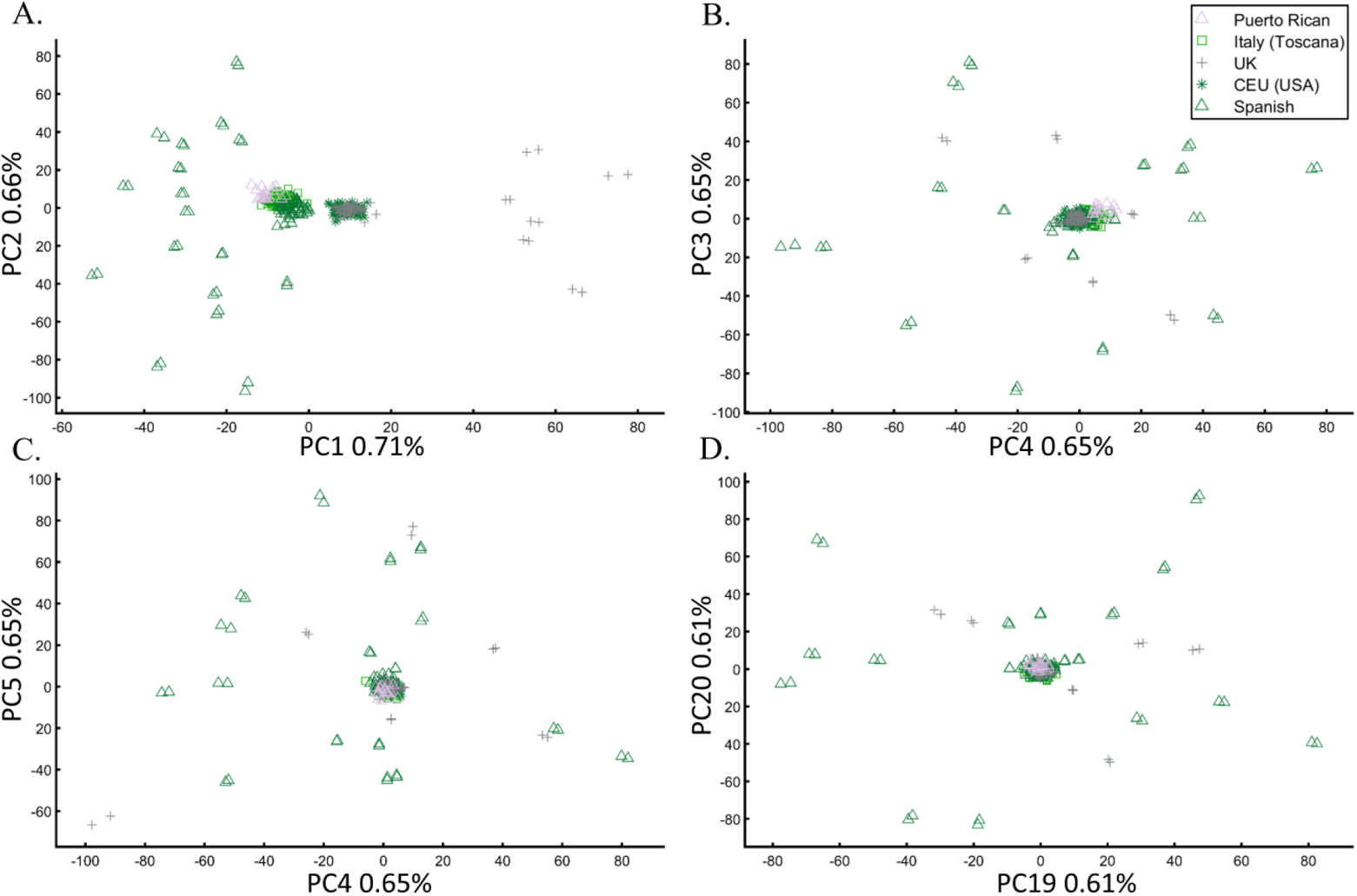
PCA applied to 20 Puerto Ricans and 300 random Europeans. The results are plotted on various PCs.

We next demonstrate that the distance between individuals and populations is meaningless and is an artifact of the cohort rather than an informative biological or demographic quantity. For that, we sought to study the relationships between Chinese and Japanese, as has been done in previous studies (e.g., Tian et al. 2008a; Wang et al. 2018b). We applied PCA to Chinese and Japanese, using Europeans as an outgroup three times. The only element that varied in those analyses was the number of Mexicans as the second outgroup (5, 25, and 50). Figures 19A-C demonstrate that the genetic distances between Chinese and Japanese were entirely dependent on the number of Mexicans in the cohort.

**Figure 19.**
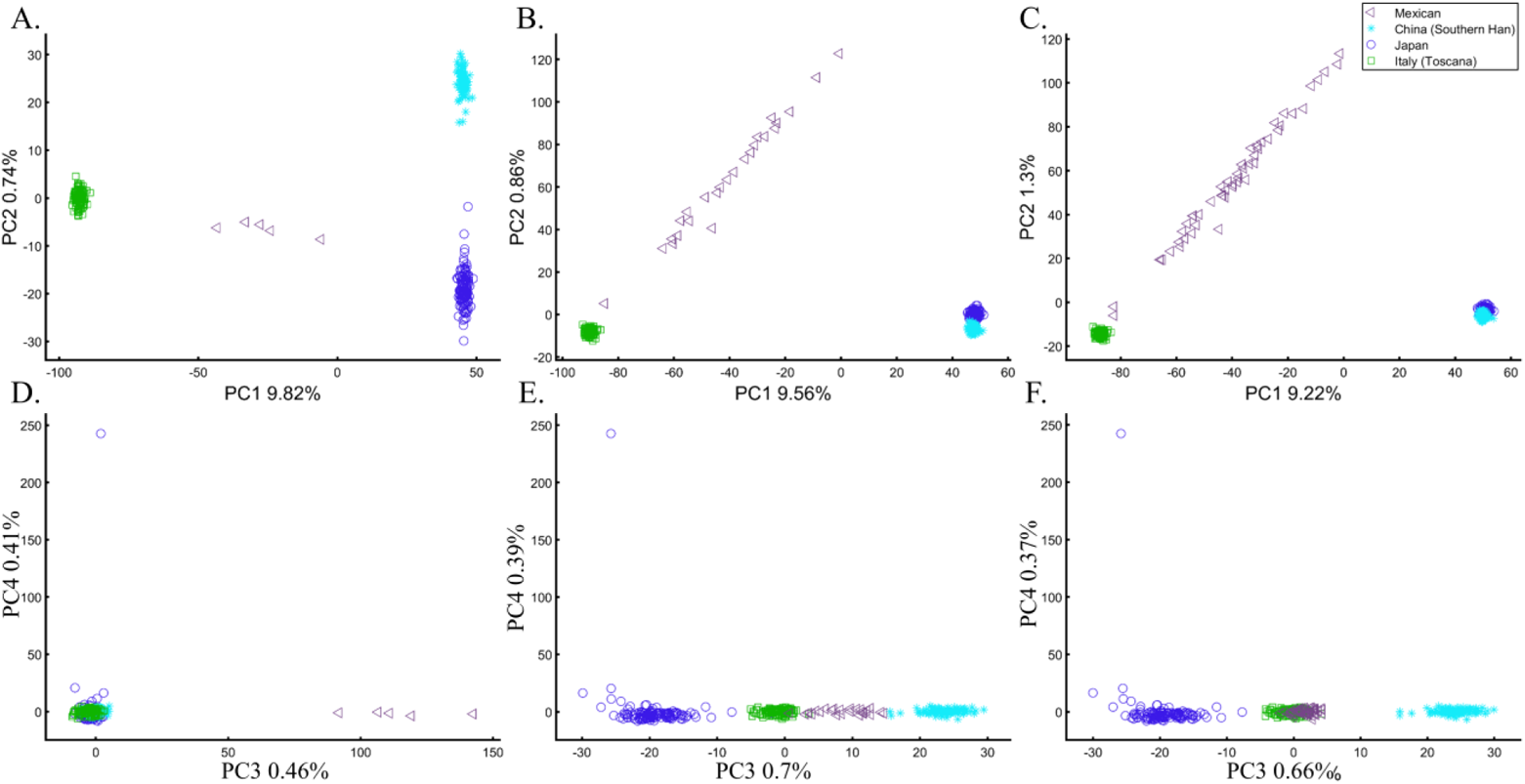
The effect of varying the number of Mexican on the inference of genetic distances between Chinese and Japanese. We analyzed a fixed number of 135 Han Chinese (CHB), 133 Japanese (JPT), 115 Italians (TSI), and a variable number of Mexicans (MXL), including 5 (A, D), 25 (B, E), and 50 (C, F) individuals over PCs 1 and 2 (A-C) and 3 and 4 (D-F). The overlap between Chinese and Japanese, typically used to infer genomic distances, was unexpectedly conditional on the number of Mexican in the cohort.

Some authors consider higher PCs to be informative and advise to consider these PCs alongside the first two. In our example, however, these PCs were not only susceptible to bias due to the addition of Mexicans but also exhibited the exact opposite pattern to that observed by the primary PCs (Figures 19D-F). These results demonstrate that the observable distances between populations in a PCA plot can be manipulated in a way that produces unpredictable results and that considering higher PCs may produce conflicting patterns.

We thereby demonstrated that PCA results (Figure 17–19), evaluated through clusters or distances between individuals, do not provide meaningful biological or historical information and cannot be used to support such claims.

### 9. The case of unions and projections

There are two common approaches to incorporate previous PCA results in a new analysis. The first is to combine PCs from past studies and is common in meta-GWAS or Mendelian Randomization studies (e.g., Hemani et al. 2018). In such studies, PCs stored in GWAS catalogs [e.g., GWAS Central as used by (de Kovel et al. 2016)] or reported for similar populations are being applied if the studied cohorts are considered similar by the experimenter (e.g., Katrinli et al. 2019). The second approach is to project the PCA results from the first dataset onto the second one (e.g., Moorjani et al. 2011).

We first combined PCs that were calculated separately for two datasets with mostly similar (Figures 20A) and dissimilar populations (Figures 20B). If merging PCs was a viable approach, we would expect similar populations to overlap and vice versa for different populations. Instead, we observed the opposite patterns, where different populations from the two datasets overlapped and the distances among the original populations were distorted, yielding incorrect results. This is because the PCs are determined by the composition of the cohort and sample sizes. PCA may also rotate the axes in some of the datasets. The more heterogeneous the two datasets are, the higher is the distortion. As previously demonstrated, this effect is more pronounced for unequal and admixed samples.

**Figure 20.**
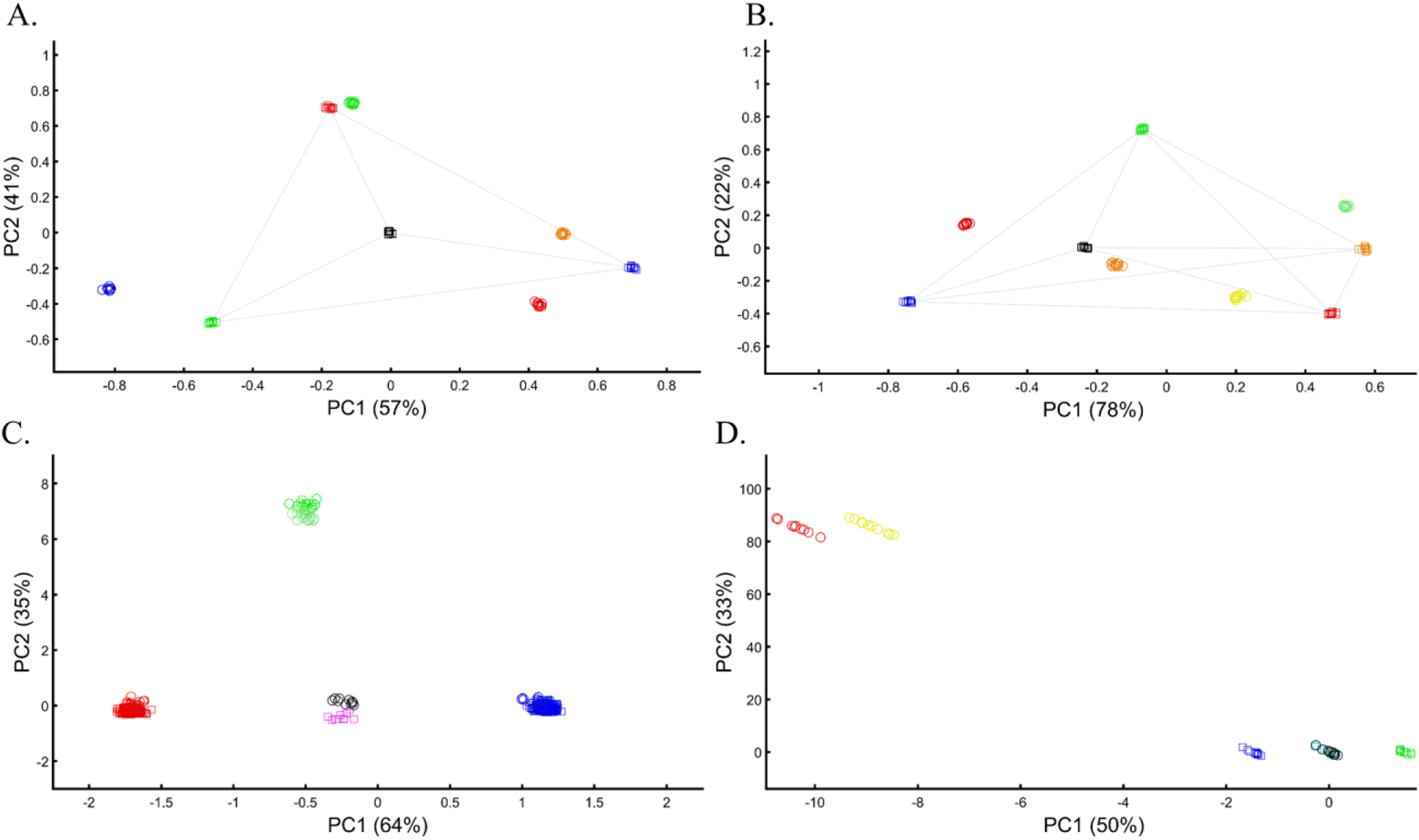
Combining two datasets in PCA. Datasets were merged either by plotting together their PCA results calculated separately for each dataset (squares and circles) (A and B) or by projecting the PCA results of one dataset (circles) onto another (squares) (C and D). An even sample size (*N*=10) was used in A, B, and D. In C, the sample size varied (10≤*N*≤300) for both datasets.

Next, we tested the projection approach, which was seemingly more successful. However, the results were dependent on the identity of the populations of the two cohorts. When the same populations were analyzed, they indeed overlapped (Figures 20C); but when unique populations were found in the two datasets, misleading matches occurred (Figures 20C-D). Overall, we found that PCA shows an overlap of different populations and distance distortion among all the populations using either approach.

The results for real populations closely followed our former results. When analyzing the same four continental populations from two datasets separately and plotting the results together, populations from the two datasets clustered incorrectly (e.g., Amerindians clustered with East Asians) (Figure 21A). When the populations in the second dataset differed from the first one, PCA distorted the results (Figure 21B). Here, Ashkenazic Jews completely overlapped with Spanish, Southern Han Chinese with Japanese, and Han Chinese with Vietnamese. UK individuals clustered closer to Africans than Europeans. When the heterogeneity of the second dataset increased, so did the distortion. When analyzing other datasets, the same populations clustered separately and overlapped with different populations (Figure 21C-D). In one case, East Asians were positioned on the trajectory between the British of the first and second cohorts (Figure 21D). We thereby demonstrated that merging PCs of different datasets does not produce a correct inference of ancestry or admixture.

**Figure 21.**
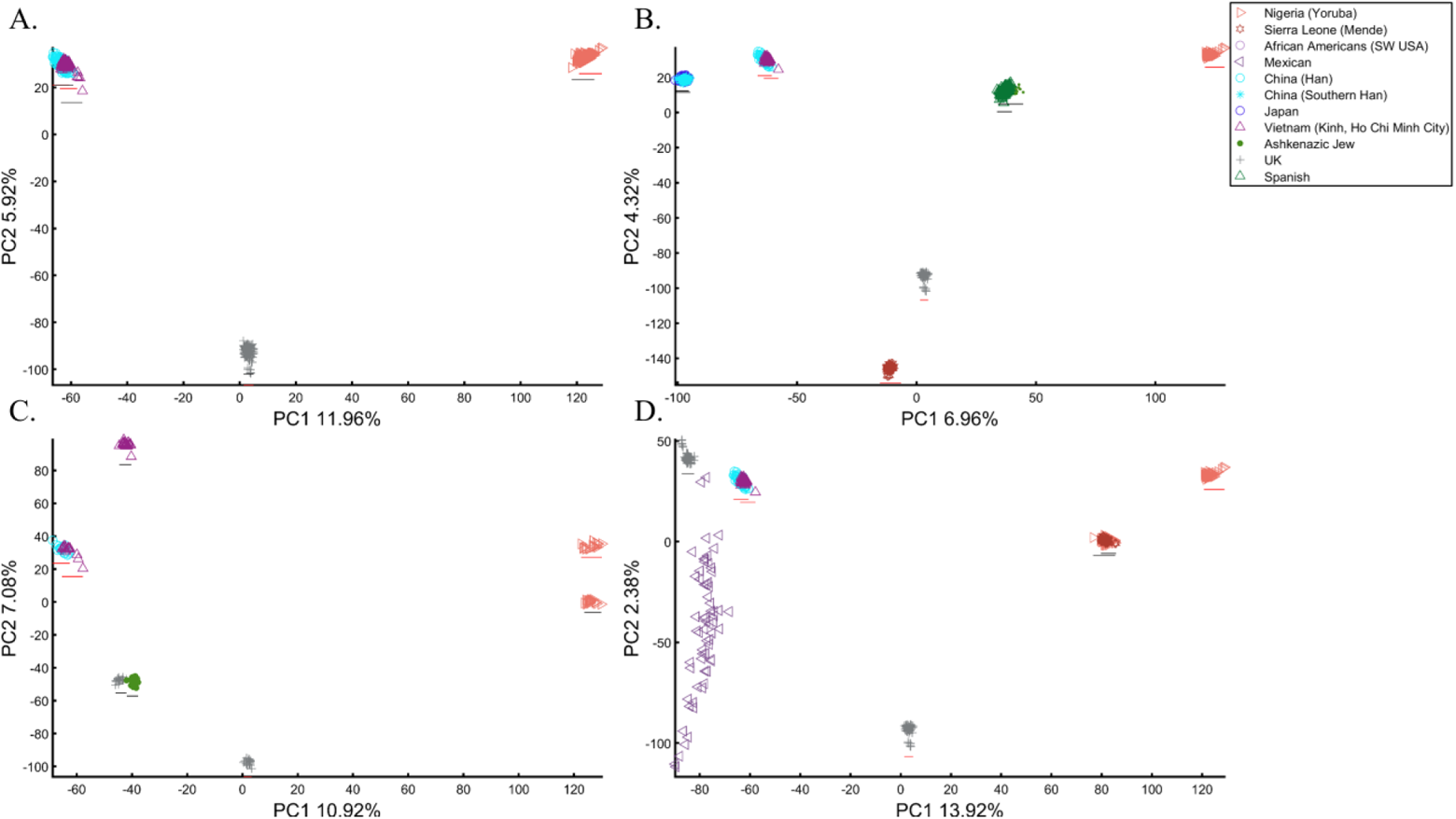
Combining population PCA results from separate datasets. PCA results were calculated separately for two datasets (red and black lines) when plotted together. The first dataset (red line) used even-sized populations in A, B, D (*N_all_*=50), and C (*N_all_*=20). In A, the second dataset had the same even-sized populations (*N*=93). The population size varied in B (*N_Nigerians_*=85, *N_Japanese_*=133, *N_AJ_*=478, *N_Spanish_*=154, *N_Chiese_*=105) and C (*N_Af_*=178, *N_Eu_*=105, *N_AJ_*=478, *N_Vietnamese_*=93). In D, it remained constant (*N_all_*=93).

There are more reasons to be concerned about using projections. They can be shown to correctly align with continental populations when the base and the projected populations are very similar (Figure 22A) and gain trust in their accuracy. However, even in the simplest scenario of using three continental populations, it is unclear how to interpret the overlap between the base and projected populations since the Spanish would not be considered genetically closer to Finns than they are closer to Italians, as suggested by PCA. In another simple scenario, where Europeans are projected onto other Europeans, distinct populations like AJs, Iberians, French, CEU, and British overlap entirely (Figure 22B), whereas Finns and Italians are separate. Not only do the results share no apparent resemblance to the geographical distribution, but they also misrepresent the genetic distances between these populations – two properties that PCA claims to represent. Adding more populations, even if only with the projected populations, contributes towards further distortion of the results with previously distinct populations (Figure 22B) now clustering together (Figure 22C). In a different dataset, projecting Japanese onto a base dataset of Africans and Europeans places them as an admixed African-European population. The projected Finns cluster with other Europeans (Figure 22D), at odds with the previous results (Figure 22B) that singled them out.

**Figure 22.**
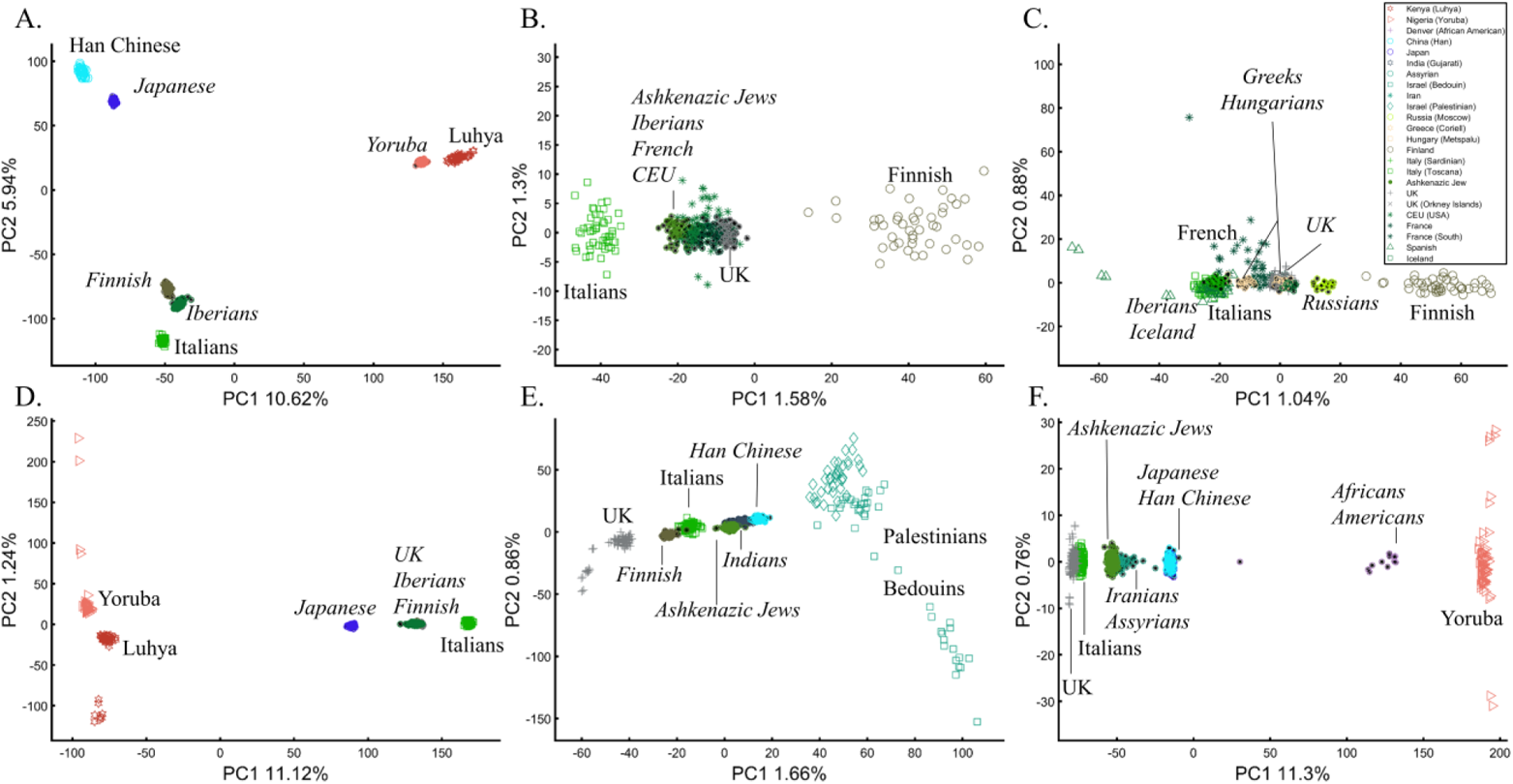
PCA projections of populations (*italic* & black star inside the shape) onto base even-sized populations (*N*=50, unless noted otherwise) (regular font). In A) *N_projected_*=100, B) *N_projected_*=50, C) *N_projected_*=20, D) *N_projected_*=100, E) *N_projected_*=80 and *N_projected_*=100, and F) 80≤*N_projected_*≤100 and 12≤*N_projected_*≤478.

To demonstrate that projecting populations that are different from the base populations yields highly misleading results, we projected Chinese, Finns, Indians, and AJs onto Levantine and two European populations (Figure 22E). The results imply that the Chinese are of an Indian origin originating from a European-Levantine mix. AJs are a sister population to the Chinese from the same Indian origin. Replacing Levantines with Africans does not add stability to the projected results (Figure 22F). Now the projected Chinese and Japanese overlap, and AJs cluster with Iranians.

### 10. The case of ancient DNA

PCA is the primary tool in paleogenomics, where ancient samples are identified based on their clustering with modern or other ancient samples. Here, a wide variety of strategies is employed. In some studies, ancient and modern samples are combined (Gamba et al. 2014). In other studies, PCA is performed separately for each ancient individual and “particular reference samples,” and the PC loadings are combined (Skoglund et al. 2012). Sometimes present-day human populations are projected onto the top two principal components defined by ancient hominins (and non-humans) (Reich et al. 2010) and sometimes, it is the other way around (Lazaridis et al. 2014).

We focused on projecting ancient populations onto modern ones, as it is the most common study design. Since ancient populations show more genetic diversity than modern ones (Lazaridis et al. 2016), we defined “ancient colors” (*a*) as brighter colors whose allele frequency is 0.95 with an SD of 0.05 and “modern colors” (*m*) as darker colors whose allele frequency is 0.6 with an SD of 0.02. Both samples had null alleles. Black was neutral. Two approaches were used in analyzing the two datasets: calculating PCA separately for the two datasets and presenting the results jointly (Figures 23A-B), and projecting the PCA results of the “ancient” populations onto the “modern” ones (Figures 23C-D). In both cases, meaningful results would show the ancient colors clustering close to their modern counterparts in distances corresponding to their true distances.

**Figure 23.**
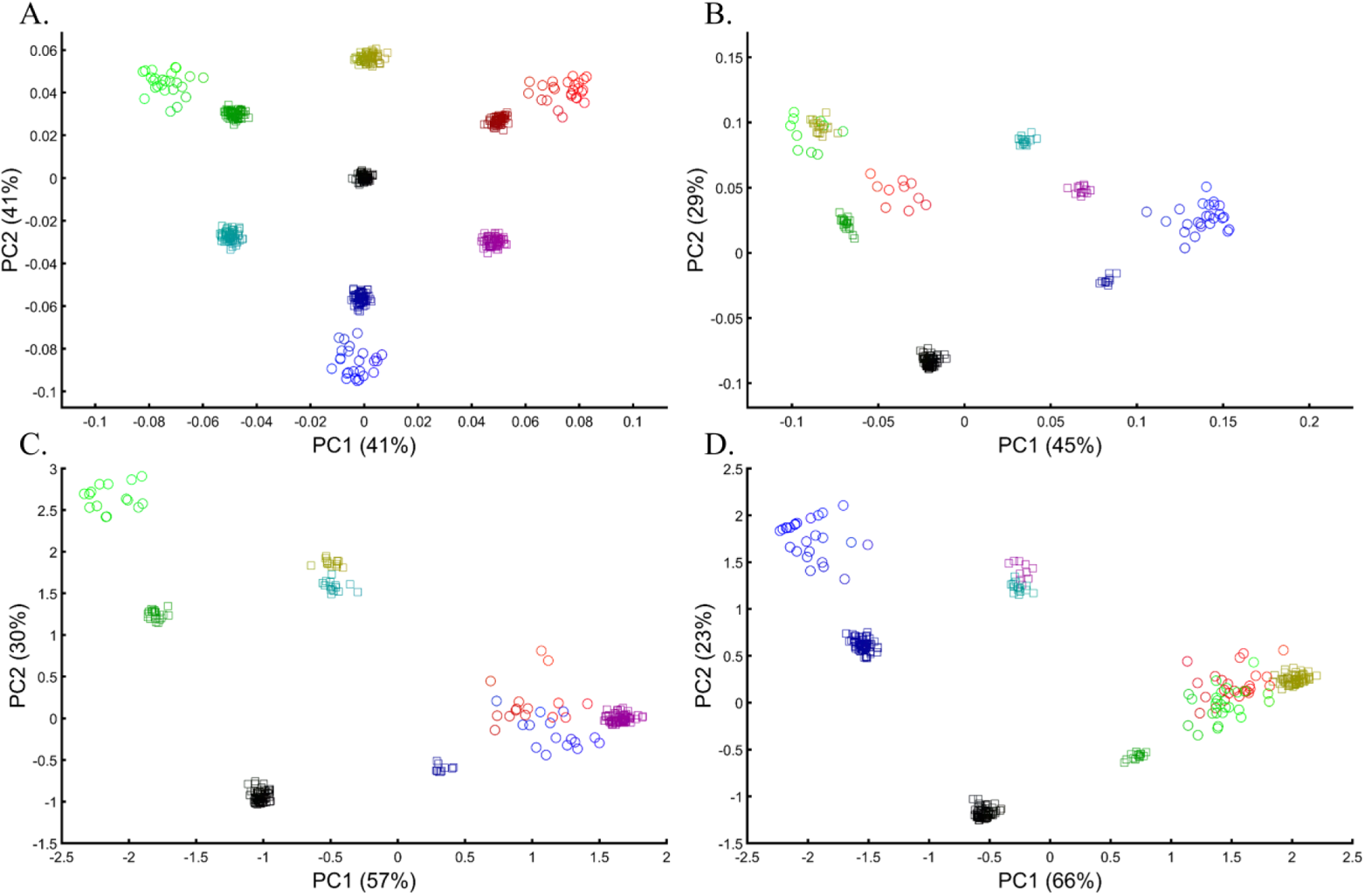
Merging PCA of “ancient” (circles) and “modern” (squares) color populations using two approaches. First, PCA is calculated separately on the two datasets, and the results are plotted together (A-B). Second, PCA results of “modern” populations are projected onto the PCs of the “ancient” ones (C-D). In A, even-sized “ancient” (*N*=25) and “modern” (*N*=75) color populations are used. In B, different-sized “ancient” (10≤*N*≤25) and “modern” (10≤*N*≤75) populations are used. In C and D, different-sized “ancient” (10≤*N*≤75) are used alongside even-sized “modern” populations: C) (*N*=15) and D) *N*=25.

These are indeed the results of PCA when even-sized “modern” and “ancient” color populations are analyzed as well as a balanced color pallet (Figure 23A). In the more realistic scenario where the color pallet is imbalanced and sample sizes differ, PCA produced incorrect results where ancient Green (aGreen) clustered with modern Yellow (mYellow) away from its closest mGreen that clustered close to aRed. mPurple appeared as 4-ways mixed of aRed, aBlue, mCyan, and mDark Blue. Black, instead of being at the center as in Figure 23A, now appeared as an outgroup, and its distances to the other colors were distorted (Figure 23B). Projecting “ancient” colors onto “modern” also highly misrepresented the relationships among the ancient samples as aRed overlapped with aBlue or aGreen, mYellow appeared closer to mCyan or aRed, and the outgroups continuously changed (Figure 23C-D). Note that the first PCs of the last results explained most of the variance (89%), not the primary PCs of the first analysis (82%).

Recently, Lazaridis et al. (2016) projected ancient Eurasians onto modern-day Eurasians and reported that ancient samples from Israel clustered at one end of the Near Eastern cline and ancient Iranians at the other, close to modern-day Jews. Insights from the positions of the ancient populations were then used in their admixture modeling that supposedly confirmed the PCA results. Using similar modern and ancient populations, we replicated the results of Lazaridis et al. (2016) (Figure 24A). However, this is neither the only possible nor most informative PCA outcome. By adding the modern-day populations that Lazaridis et al. (2016) omitted, we can show that the ancient Levantines cluster with Turks (Figure 24B), Caucasians (Figure 24C), Iranians (Figure 24D), Russians (Figure 24E), and Pakistani (Figure 24F) populations. The overlap between the ancient Levantines and other populations also varied widely, whereas they cluster with ancient Iranians and Anatolians, Caucasians, or alone, like a “population isolate.” Moreover, the remaining ancient populations exhibited conflicting results inconsistent with our understanding of their origins. Mesolithic and Neolithic Swedes, for instance, clustered with modern Eastern Europeans (Figure 24A-C) or remotely from them (Figure 24D-F). It is difficult to justify Lazaridis et al.’s (2016) preference for the first outcome where the first two components explained only 1.35% of the variation (in our replication analysis. Lazaridis et al. omitted the proportion of explained variation) (Figure 24A), compared to all the alternative outcomes that explained a much larger portion of the variation (1.92-6.06%).

**Figure 24.**
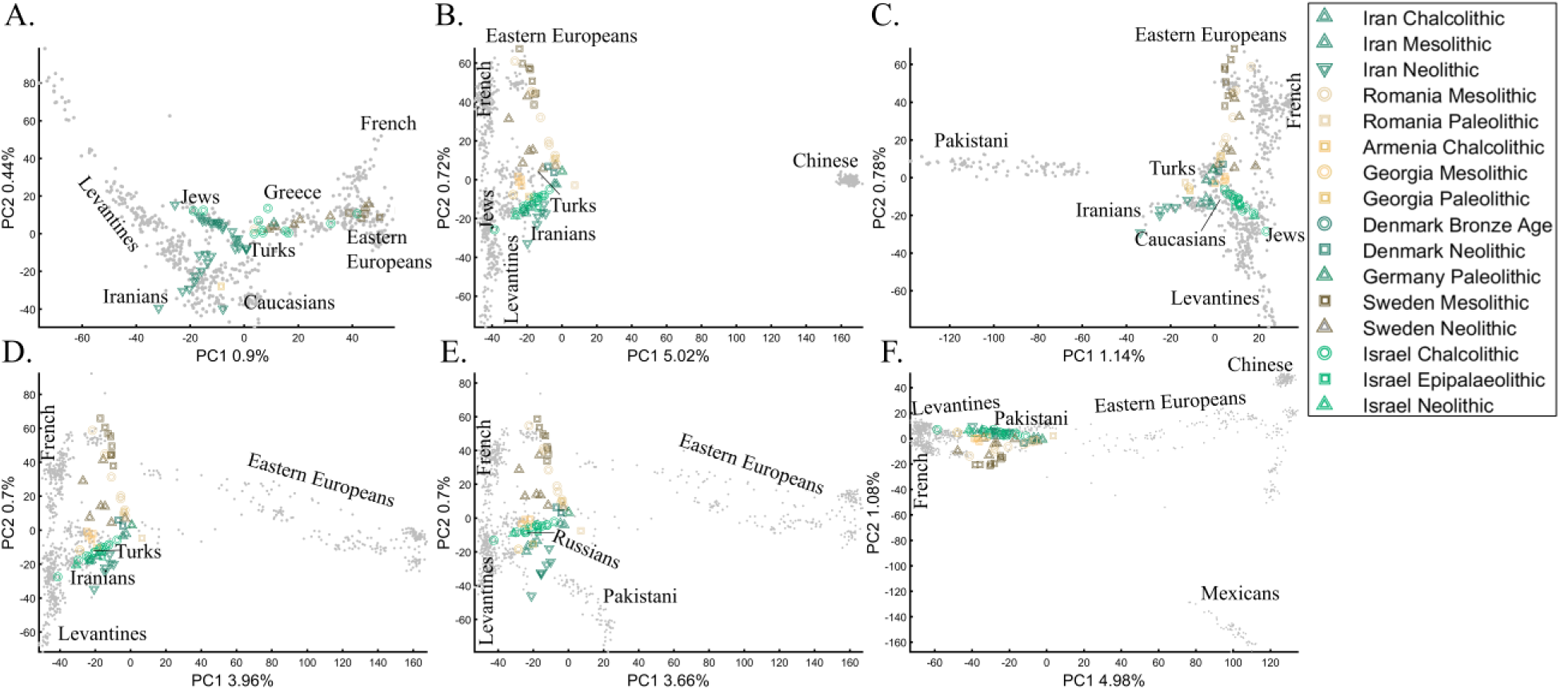
PCA of 65 ancient Palaeolithic, Mesolithic, Chalcolithic, and Neolithic from Iran (12), Israel (16), the Caucasus (7), Romania (10), Scandinavia (15), and Central Europe (5) projected onto modern-day populations of various population sizes. In addition to the modern-day populations used in A), the following subfigures also include B) Han Chinese, C) Pakistani (Punjabi), D) additional Russians, E) Pakistani (Punjabi) and additional Russians, and F) Pakistani (Punjabi), additional Russians, Han Chinese, and Mexicans. The ancient samples remained the same in all the analyses. In each of the plots (A-F), the ancient Levantines cluster with different modern-day populations. Present-day individuals are shown in grey dots alongside their broad population labels for coherency. The full population labels are shown in Figure S4.

### 11. The case of marker choice

The choice of markers is another primary concern in PCA that received little attention in the literature. Although PCA is routinely applied to different SNP sets, the PCs are typically deemed comparable. In forensic applications, where sample sizes usually range from 100 to 300 markers, this is a major problem. Thereby, for our last test case, we evaluated the effect of various markers on PCA outcomes. It is unfeasible to use our color model to study biologically different markers, yet it can be used to study the effects of missing data and noise, which are common in genomic datasets and reflect the biological properties of different marker types in capturing the population structure. Compared to the original figure (Figure 8A), the addition of 50% (Figure 25A) and even 90% missingness (Figure 25B) allowed recognizing the original population structure. The structure decayed when random noise was added to the latter dataset (Figure 25C). To further explore the effect of noise, we added random markers to the dataset. An addition of 10% of noisy markers increased the dataset’s disparity, but it still retained the original structure (Figure 25D). Interestingly, even adding 100% noisy markers allowed identifying the original structure’s key features (Figure 25E). Only when adding 1000% noisy markers did the original structure disappear (Figure 25F). Note that the introduction of noise has also sliced the percent of variation explained by the PCs. These results highlight the importance of using ancestry informative markers (AIMs) to uncover the true structure of the dataset and accounting for disruptive markers.

**Figure 25.**
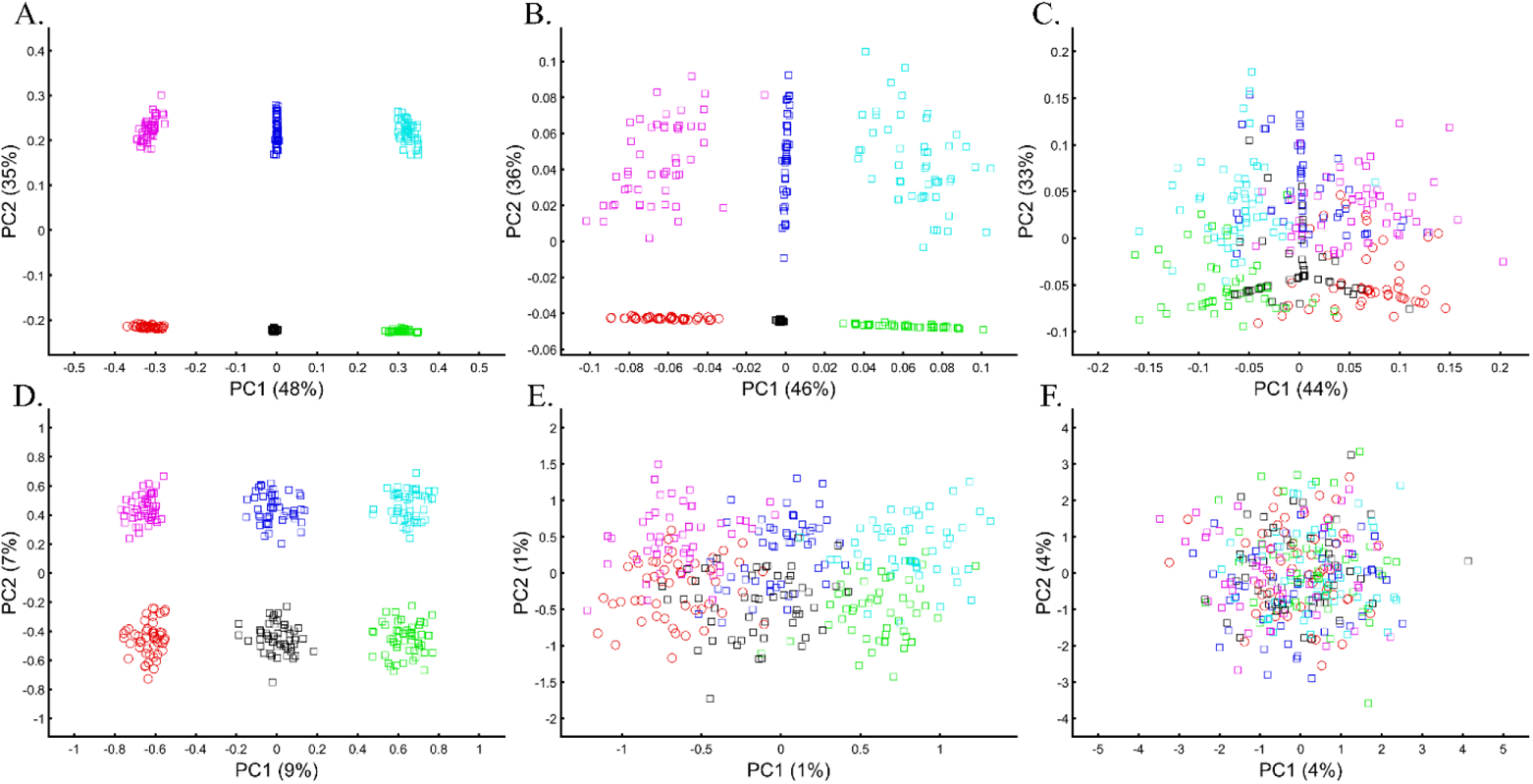
Testing the effects of missingness and noise in a PCA of six fixed-size (*n*=50) color populations. The top plots show the effect of missingness alone or combined with noise: A) 50% missingness, B) 90% missingness, and C) 90% missingness and low-level random noise in all the markers. The bottom plots test the effect of noise when added to the original markers in the above plots using: D) 30 random markers, E) 300 random markers, and F) 3,000 random markers.

Different marker types represent the population structure differently, with coding markers carrying the least and non-coding the most information. To illustrate the extent to which marker types represent the population structure, we studied the relationships between UK British and other Europeans (Italians and Iberians) using different types of 30,000 SNPs, a number consistent with the number of SNPs typically analyzed (e.g., Hyde et al. 2016; Watkins et al. 2016; Wright et al. 2019). According to the full SNP set, the British do not overlap with Europeans (Figure 26A). However, coding SNPs show considerable overlap (Figure 26B) compared with intronic SNPs (Figure 26C). Protein coding SNPs, RNA molecules, and upstream or downstream SNPs (Figure 26D-F, respectively) also show small overlap. The identification of “outliers,” already a subjective measure, may also differ based on the proportions of each marker type. These results not only illustrate how the choice of markers and populations profoundly affect PCA results but also the difficulties in recovering the population structure in exome datasets.

**Figure 26.**
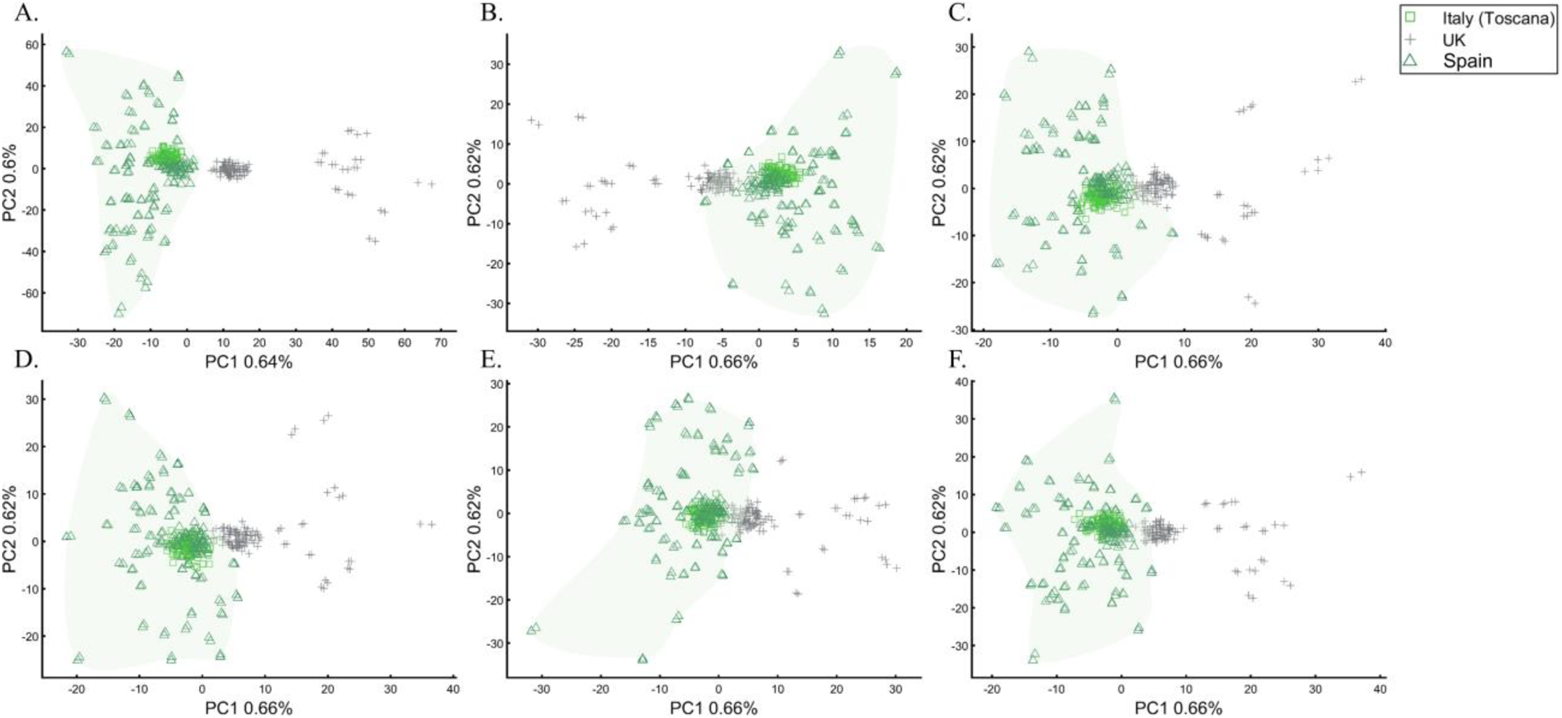
PCA of Tuscany Italians (*N*=115), British (*N*=105), and Iberians (*N*=150) across all markers (*N*∼129,000) (A) and different marker types (*N*∼30,000): B) coding SNPs, C) intronic SNPs, D) protein-coding SNPs, E) RNA molecules, and F) upstream and downstream SNPs. The Italian-Spanish cluster is shadowed.

## Evaluating the core properties of PCA

Several limitations of PCA are worth highlighting since they may not have been evident in the test cases reviewed here. First, PCA typically explains a tiny part of the variation (Figure S5), which does not only grow smaller as more samples are added (Figure S5) but also grows in inaccuracy (Figure 10). This leads to a paradox, whereas increasing the sample size, which intuitively should be expected to increase the accuracy of analyses, decreases the proportion of explained variance. Second, analyzing only the two primary components does not solve the rapid decline in the proportion of explained variation (Figure S6). Interestingly, the average variance explained by the two primary PCs over hundreds and thousands of individuals is on average 0.12 (Figure S6, inset), the same amount of genetic variation distributed between continental populations calculated by hierarchical *F_ST_* (Elhaik 2012). Third, PCs higher than three not only explain a minuscule amount of variation, they also cannot distinguish the true data structure from noise (Figure S7). In other words, PC plots where two primary components explain around 1% of the variance, likely as in Lazaridis et al. (2016), capture as much of the population structure as they would from a randomized dataset. Recall that all the datasets analyzed here consist of ancestry informative markers (AIMs) and are thereby optimized for discovering population structure. The fourth concerning the characteristic of PCA is the “big-p, little-n,” where *p* stands for predictors and *n* for samples, otherwise known as the *p*>>*n* problem or simply the curse of dimensionality (Bellman 1961; Rohlf 2021). Briefly, it refers to the phenomenon that arises when analyzing data in high-dimensional spaces unobserved in lower-dimensional space. As a dimensionality reduction technique, PCA aims to address this problem; however, PCA introduces biases of its own. We observed that PCA misrepresents the distances even in the simplest and near-perfect analysis of four colors (Figure 1C). We have also seen PCA repeatedly misrepresent the true distances for color populations. In high-dimensional space, the distances between the data points increase compared to low-dimensional space (Figure S8). As such, samples that are closer to one another appear more distanced and no longer cluster. In other words, cases and controls cannot be reliably identified in high-dimensional data. PCA violations of the true distances between samples limit the ability to reduce high-dimensionality genetic variation data correctly.

## Discussion

The reproducibility crisis in science called for a rigorous evaluation of scientific tools and methods. Due to PCA’s centrality in population genetics, and since it was never proven to yield correct results, we sought to assess its reliability, robustness, and reproducibility. Here, we evaluated PCA performances for eleven use-cases using a simple color-based model where the true structure was known and real populations and showed that PCA failed in all three measures.

We found that PCA is an unreliable tool because, excepting the simplest model case (Figure 1C), it has failed to produce correct results across all the design schemes, whether even-sampling was used or not, and whether for least or more admixed populations. Whereas the clustering of populations between other populations in the scatter plot has been regarded as “decisive proof” or “very strong evidence” of their admixture (Patterson et al. 2010), we demonstrated here that such patterns are an artifact of the sampling scheme and meaningless for any bio historical purposes. The clustering of the results, a subject that received much attention in the literature (e.g., Patterson, Price, and Reich 2006), is another artifact of the sampling scheme and likewise biologically meaningless (Figures 13–18). Overall, the notion that PCA can yield biologically or historically meaningful results is a misconception supported by *post hoc* reasoning. Like a broken watch, PCA *can* be engineered through careful data manipulation (e.g., Figure 24) to yield presumed correct results. However, it is not a guaranteed outcome, and those “correct” results are indistinguishable from alternative and contradictory results constructed using other schemes. That all the results presented here for either the color or real populations are mathematically true but realistically and biologically false proves that bio historical inferences drawn only from PCA are fictitious. PCA is highly subjected to minor alterations in the allele frequencies (Figure 12), study design (e.g., Figure 11), or choice of markers (Figure 26) (see also Wang et al. 2015; Elhaik and Ryan 2019). Finally, PCA results cannot be reproduced (e.g., Figure 13) unless an identical dataset is used, which defeats the usefulness of this tool. Note that variations in the implementations of PCA (e.g., PCA, singular value decomposition [SVD], and recursive PCA), as well as using the various flags, as implemented in EIGENSOFT, yield major differences in the results, none more biologically correct than the other.

**Figure 13.**
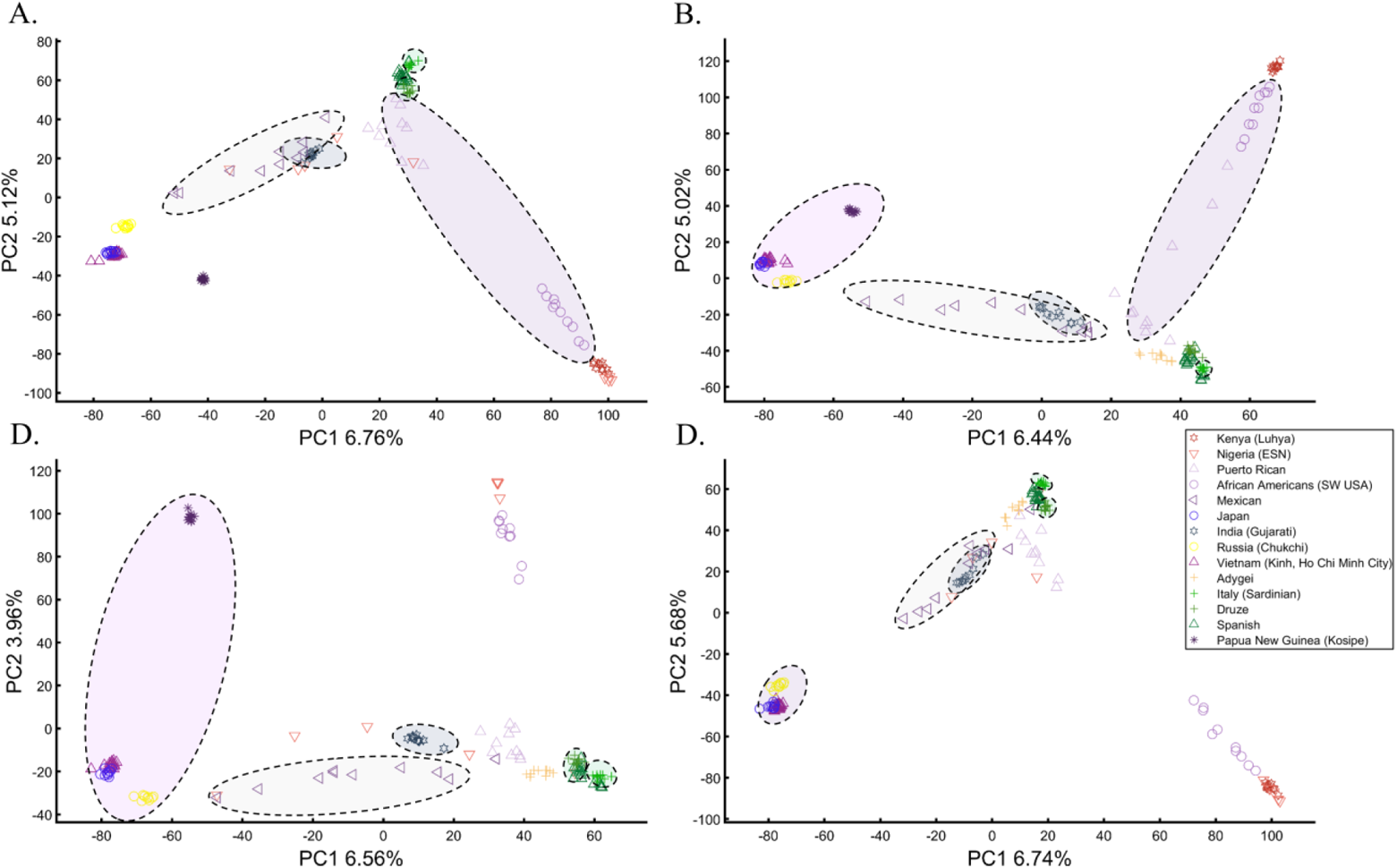
Studying the effect of sampling on PCA results. A cohort of 16 worldwide populations (see legend) was selected. In each analysis, a random population was excluded. Populations were represented by ten random samples. The clusters highlight the most notable differences.

Several aspects of this study are important to emphasize. First, this study does not ask whether PCA is *correct* or *wrong* but whether it gives *true* or *false* results in genetic studies. PCA is a computational procedure that computes the principal components and uses them to change the basis of the data. PCA has multiple implementations and broad applications in various fields. Thereby, if properly implemented, PCA is considered *correct*. This study is concerned with whether PCA results are genetically and historically *true* and informative in understanding the patterns in the dataset better than other procedures. Second, this study focuses on genetic variation data, particularly human data, that have a particular behavior. For other data types or datasets not tested here, PC analyses may be more successful (e.g., Elhaik, Graur, and Josić 2010). We note, however, that PCA criticism is neither rare nor unique to genetics (e.g., criticism of PCA for geology (Trochimczyk and Chayes 1977)). To better understand how PCA reached prominence, we shall review the historical debate on whether PCA represents the genetic data correctly.

### A brief history of PCA and its application to population genetics

It is well-recognized that Pearson (1901) introduced PCA and Hotelling (1933) the terminology. Hotelling’s motivation was to address the problem of evaluating independent mental traits in psychology. Thurstone presented another principal axes solution to the problem of factor analysis (Hotelling 1933). However, he later reconciled, as he could not see how they describe a meaningful psychological model (Thurstone 1935). The argument about the truthfulness and reliability of PCs continued to this day (Hubert 2016).

In population genetics, PCA is primarily used to reduce the dimensionality of multivariate datasets by linearly transforming the genotypes into a set of mutually uncorrelated principal components (PCs) ranked according to their variances. As most of the original variability is contained in the primary two PCs, they are typically visualized on a colorful scatter plot. The early work of Cavalli-Sforza suggested that PCA can detect ancient migrations and population spreads (Menozzi, Piazza, and Cavalli-Sforza 1978) and uncover “hidden similarities” (Piazza, Menozzi, and Cavalli-Sforza 1981) in the genomic data. The authors proposed that PCA will “give us new insight into the evolutionary history of the populations represented in the map” and that “using a different color for each one and making use of the capacity of the human eye to synthesize three colors.” Cavalli-Sforza’s arguments were not very convincing.

During the 20^th^ century, PCA was sparsely employed in genomic analyses alongside other multidimensional scaling tools. The next-generation sequencing revolution in the early 21^st^ century resulted in the emergence of large genomic datasets that required new and powerful computational tools with appealing graphical interfaces, like STRUCTURE (Pritchard, Stephens, and Donnelly 2000). PCA was not used in the publications of the first two HapMaps nor the HGDP dataset (The International HapMap Consortium 2005; Conrad et al. 2006; The International HapMap Consortium 2007).

In 2006, Price et al. (2006) introduced the SmartPCA tool (EIGENSOFT package) and claimed that the method has “a solid statistical footing” that can “discover structure in genetic data” even in admixed populations. Those claims were made based on a simulated dataset and an application of PCA to a dataset of European Americans, which revealed an incoherent pattern claimed to reflect genetic variation between northwest and southeast Europe. Simultaneously, Patterson et al. (2006) applied PCA to three African and three Asian populations and claimed that the dispersion patterns of the primary two PCs reflect the true population structure.

The next milestone in the rise of PCA to prominence was the work of Novembre and colleagues (2008) that showed a correlation between PCA and geography among Europeans. The authors applied PCA to the dataset and positioned the PCs on Europe’s map and rotated their axis to increase the correlation with Europe’s map. After fitting a model of longitude and latitude that included PC1, PC2, and their interactions, samples were positioned on Europe’s map. The authors claimed that “The resulting figure bears a notable resemblance to a geographic map of Europe” and reported that, on average, 50% of samples from populations with greater than six samples were predicted within less than 400km of their country. Most of those populations, however, were from the extreme ends of the map (Italy, UK, and Spain) and were predicted most accurately because PCA maximizes the variance along the two axes. By contrast, samples from mid and north-Europe were predicted most poorly. Overall, the authors’ approach classified about 50% of the samples in the final dataset to within 400km of their countries. Looking at samples from all the European countries (their Table 3), only 24% were predicted to their correct country, 50% of the populations were predicted within 574km (about the distance from Berlin to Warsaw), and 90% of the populations were predicted within 809km (about the distance from Berlin to Zurich). Overall, it is fair to say that in practice, this method does not perform as implied because it strongly depends on the specific cohort. Therefore, it does not have any practical applications. A more proper title for the paper would have been “populations can be selected to mirror geography in a quarter of Europe.” Later, Yang et al. (2012) claimed to have expanded the method to global samples. Elhaik et al. (2014) showed that the new method has less than 2% accuracy, with some samples being predicted outside our planet. Thus far, no PCA or PCA-like application has ever reached an accuracy higher than 2% worldwide (Mason-Buck et al. 2020). By contrast, an admixture-based approach achieved 83% accuracy in classifying individuals to countries and even islands and villages (Elhaik et al. 2014).

Ignoring these methodological problems and further promoting their PCA tool, Reich and colleagues (2008) wrote that “PCA has a population genetics interpretation and can be used to identify differences in ancestry among populations and samples, regardless of the historical patterns underlying the structure” and that “PCA is also useful as a method to address the problem of population stratification—allele frequency differences between cases and controls due to ancestry differences—that can cause spurious associations in disease association studies” and finally that “PCA methods can provide evidence of important migration events” – none of which were supported by the work of Novembre et al..

After its applications to the HGDP (Biswas, Scheinfeldt, and Akey 2009) and HapMap 3 (Altshuler et al. 2010) datasets, PCA became the foremost utility in population genetic investigations, reaching ”fixation,” the point where it is used almost in every paper in the field, by 2013 (Figure 27).

**Figure 27.**
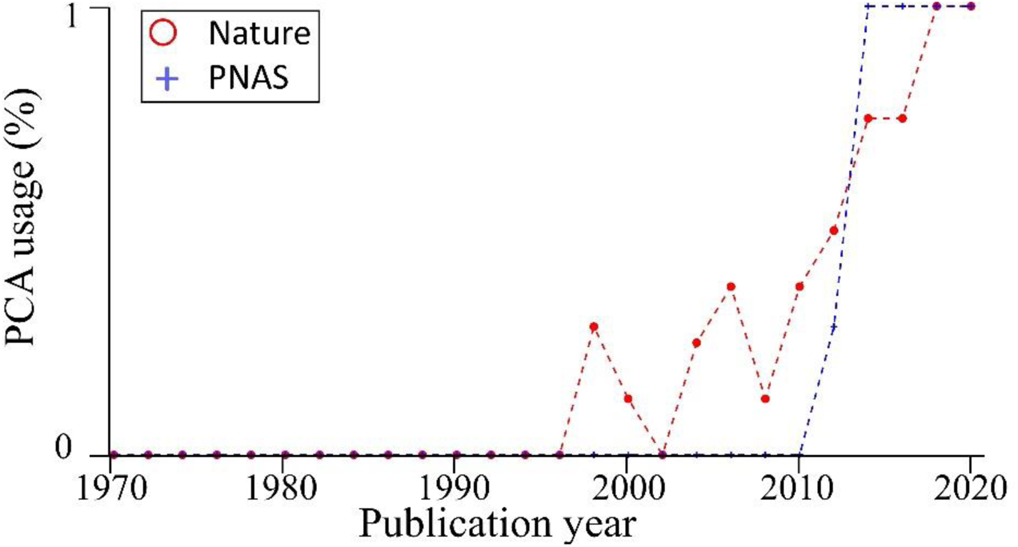
Evaluating the existence of a PCA in population genetic publications by sampling four random population genetic papers per year from Nature and PNAS. The percent of publications that used at least one PCA is shown.

### Misuses of PCA in the literature

To understand why a tool with so many limitations became the foremost tool in population genetics, we will briefly review how authors handled limitations.

We already demonstrated that the authors had misinterpreted PCA findings and did not disclose the amount of variation explained by PCA. Fasciently, in 2008 Reich and colleagues found it necessary to assess “whether the proportion of the variance explained by the first PC is sufficiently large,” most likely before they realized just how small this variation really is. To the best of our knowledge, their numerous publications that employed PCA have all omitted that information (e.g., Reich et al. 2009; Reich et al. 2010; Pickrell et al. 2012; Moorjani et al. 2013; Lazaridis et al. 2016; Mathieson and Reich 2017; Olalde et al. 2019).

Remarkably, Novembre and Stephens (2008) warned that “PCA results depend on the details of a particular dataset, they are affected by factors in addition to population structure, including distribution of sampling locations and amounts of data. Both these features limit the utility of PCA for drawing inferences about underlying processes” but nonetheless found PCA to be “undoubtedly an extremely useful tool for investigating and summarizing population structure,” and correctly anticipated that it will play “a prominent role in analyses of ongoing studies of population genetic variation.”

Although many of the aforementioned authors were aware that PCA results depended on the sample cohort, they continue using it, presenting only the results that fit their hypotheses. For example, Tian et al. (2008b) were aware that PCA “is sensitive to differences in the inclusion or exclusion of specific population groups” and that it “can be dramatically affected by differences in relatively small genomic regions that may not reflect true population substructure.” Likewise, Tian et al. (2009) noted that Ashkenazic Jews (AJs) “have a unique genotypic pattern that may not reflect geographic origins” and that “the inclusion or exclusion of particular ethnic groups… shifted the relationships in PCA.” They acknowledged that their findings “show that PCA results are highly dependent on which population groups are included in the analysis.” Still, both groups drew conclusions based on PCA. It should come as no surprise that PCA (Tian et al. 2008b; Tian et al. 2009; Atzmon et al. 2010; Behar et al. 2010; Behar et al. 2013) and, in one case, Multi-dimensional scaling (MDS) (Kopelman et al. 2020) spearheaded claims of Levantine origin for AJs. Here, we showed that PCA could be easily engineered to foster American, Iberian, West, Central, and South European, Britain, Scandinavian, Indian, Iranian, Turkish, Caucasian, and even Levantine origins for AJs.

PCA has been criticized by several groups. McVean (2009) cautioned that “Sub-sampling from populations to achieve equal representation, as in Novembre et al. (2008), is the only way to avoid this problem [=the distortion of the projection space]” and that “the influence of uneven sample size can be to bias the projection of samples on the first few PCs in unexpected ways.” However, these statements are incorrect. First, Novembre et al.’s population sizes ranged from 1 to 219. Second, McVean’s simulation was limited to the case of symmetric populations arranged in a lattice formation, as in Figure 1C or Figure 23A. This led McVean to believe that PCA accuracy can be achieved when sample sizes are even and thereby have some merit (“The result provides a framework for interpreting PCA projections in terms of underlying processes, including migration, geographical isolation, and admixture”). Had McVean explored the slightly more realistic case of even-sized populations with uneven contributions to the covariance matrix (e.g., Figures 5A and 8A), he would have realized that PCA’s accuracy is extremely limited to well-controlled simulations of even-sized populations that are isotropic (symmetrically distributed across all the dimensions). In reality, “populations” are unknown, are of uneven sizes, and are anisotropic. These limitations invalidate PCA as a tool for population genomic studies.

Elhaik and Ryan (2019) showed that PCA could not model admixed samples. Elhaik et al. (2014) showed that PCA-like tools could not be used for biogeography. François et al. (2010) observed that the gradients observed in the first PC are often contrary to most often formulated expectations and offered a biological explanation to the phenomenon. They concluded that PCA should be considered as a data exploration tool and that interpreting the results in terms of past routes of migration “remains a complicated exercise.” Björklund (2019) raised concerns about sampling problems that render PCA biologically meaningless and provided several recommendations, like evaluating the distinctness of the PC’s and present the percent of explained variance. Our findings, albeit in population genetics, demonstrate that with the minor exceptions discussed above, all PCA results are wrong and are independent of the level of “cautiousness” exhibited by the reader. The practice of ignoring sample dates in paleogenomic analyses has also been criticized (Francois and Jay 2020).

### PCA as a *Dataism* exercise in population genetics

*Dataism* describes an ideology formed by the emergence of Big Data, where measuring the data is the ultimate achievement (Brooks 2013). Dataism proponents believe that with sufficient data and computing power, the world’s mysteries would reveal themselves. Dataism enthusiasts rarely ask themselves *if PCA results are correct* but rather *how to interpret the results correctly*. As such, clustering is interpreted as *common ancestry* and its absence as *genetic drift*. Populations nested between other populations are *admixed* or *isolates,* and those at the extreme edges of the PC scatter plot are *unmixed, pure,* or *races*.

Although a newly coined term, the roots of the dataism philosophy are traceable to the Hotelling-Thurstone debate and specifically to the Cavalli-Sforza-Sokal conundrum. Cavalli-Sforza et al. (1994: 338) explained the first six components in ancient human cross-continental expansions, but they never explained to what extent those historical inferences were distinguished from the null hypothesis since they did not have any. Sokal and colleagues were set to test that point and showed that the PCA maps are subject to substantial errors and that apparent geographic trends may be detected in spatially random data (the null). Sokal et al. did not express doubt in human history but only that it reveals itself in the PC maps, as do we. Cavalli-Sforza’s group responded that Sokal et al.’s sampling scheme was extremely irregular (Rendine et al. 1999) and questioned Sokal et al.’s disbelief in a wrong method that yields a conclusion that they were willing to accept otherwise. Sokal et al. (1999a) were concerned with the lack of response to their original inquiries, the PC’s interpolation (to overcome gaps in the data) and smoothing technique that introduced more noise, the specific sampling scheme of Cavalli-Sforza and colleagues that appeared incidental rather than genuinely comprehensive, and the continued absence of a null model. In further criticism of Cavalli-Sforza et al. (1994), they claimed that whereas some of the results are biologically sound, others are not, yet both are discussed equally seriously. Cavalli-Sforza (Manni 2010) stuck by PCA and the historical inferences (The Neolithic spread to Europe made “between 8,000 and 5,000 years ago”) that can be allegedly derived from it. In other words, whereas Cavalli-Sforza and colleagues believed that once sufficient data are available, the value of PCA for bio-history would reveal itself, Sokal and colleagues questioned the robustness and reliability of the approach to generate valid historical and ethnobiological results and cautioned that data that “have been interpolated or smoothed, invite ethnohistorical interpretation by the unwary” (Sokal, Oden, and Thomson 1999b). The issues at the heart of the debate were not as much about biostatistics as about dataism.

At first, Sokal and colleagues had the upper hand in the debate. PCA was not used in the first Big Data analyses of 2003-2005 until resurrected by Price et al. (2006). Price et al. ignored Sokal’s reasoning. They produced no null model nor proved that the method yields biologically correct results. The appeal of their tool was mainly its applicability to the large datasets that emerged and visual appeal. Interestingly, Novembre and Stephens (2008) showed that the PCA structured patterns that Cavalli-Sforza and others have interpreted as migration events are no more than mathematical artifacts that arise when PCA is applied to standard spatial data in which the similarity between locations decays with geographic distance. Nonetheless, their warning was largely ignored, perhaps because the parallel study of Novembre et al. (2008) left a stronger impact, and Cavalli-Sforza’s dataism was vindicated.

Evidently, PCA produces patterns no more historical than Alice in Wonderland and bear no more similarity to geographical maps than to Alice’s rabbit (Figure S9). Remarkably, even a PCA-drawn rabbit explains as much of the genomic variation as Lazaridis et al.’s PCA (2016, Figure 1b).

Overall, the positioning of a method that lacks any measurable power, a test of significance, or a null model, which any diligent scientist should seek at the forefront of population genetic analyses, is problematic at the very least. It would not be an exaggeration to consider PCA the Rorschach of population genetics, a test that is entirely open to manipulations and interpretations, where some may see “geographical maps” and others bunnies (Figure S9) or mermaids (Figure S10). When using PCA, almost all the answers are equally acceptable, and the truth is in the eyes of the beholder.

As an alternative to PCA, we note the advantages of a supervised machine-like model. In this mode, gene pools are simulated from a collection of geographically localized populations. Next, the ancestry of all tested individuals is estimated in relation to these gene pools. In this model, all individuals are represented as the proportion of the gene pools. Their results do not change when samples are added or removed in the second part of the analysis. Population groups are bounded within the gene pools, and inclusion in these groups can be evaluated. This model was shown to be reliable, replicable, and accurate for many of the applications discussed here, including biogeography (Elhaik et al. 2014), population structure modeling (Das et al. 2016), ancestry inference (Baughn et al. 2018), paleogenomic modeling (Esposito et al. 2018), forensics (Mason-Buck et al. 2020), and cohort matching (Elhaik and Ryan 2019).

## Conclusions

PCA is a mathematical transformation that reduces the dimensionality of the data to a smaller set of uncorrelated dimensions called principal components (PCs), which has numerous applications in science. In population genetics alone, PCA usage is ubiquitous, with nearly a dozen of standard applications. PCA is typically the first and primary analysis, and its outcomes determine the study design. That PCA is completely non-parametric is the source of its strength. Any genotype dataset can be processed, typically rapidly, with no concerns about parameters or the validity of the data. It is also a weakness because the answer is unique and depends on the particular dataset, which is when reliability, robustness, and reproducibility become a concern.

The implicit expectation employed by PCA users is that the variance explained along the first two PCs provides a reasonable representation of the complete dataset. When this variance is minuscule (as often with human populations), it poorly represents the data. Rather than consider using alternative analyses, authors choose not to report the variation explained by PCA.

Here, we carried out extensive analyses on eleven use-cases of PCA, using model- and real-populations to evaluate the reliability, robustness, and reproducibility of PCA. We demonstrated that PCA failed in all criteria and showed how easily it could generate erroneous and contradictory results. The dominance of PCA to population genetics is difficult to justify as it lacks any measurable significance or accuracy, excepting the quantity of explained variance, which is not only minuscule to the extent that authors avoid reporting it but, as shown here, is not a proxy to the reliability of the results. As such, PCA can generate desirable outcomes that fit preconceived hypotheses. As a “black box” basking in bioinformatic glory free from any enforceable proper usage rules, PCA misappropriations, demonstrated here for the first time, are nearly impossible to spot. Our findings raise concerns about the validity of results reported in the literature of population genetics and adjacent fields like animal and plant or medical genetics (PCA tools were cited over 15,000 times) that relied on PCA outcomes. Researchers from those fields may be even less aware of the inherent biases in PCA and the veracity of results that it can generate. We consider PCA scatterplots analogous to Rorschach plots and show using “PCA-art” how artistic images can be drawn by applying PCA to genetic data. We find PCA unsuitable for population genetic investigations and recommend that the findings of genetic studies that employed PCA should be reevaluated.

## Methods

### Sample collection

Alongside the simulated color datasets, we assembled three datasets in our evaluation of PCA performances for human populations: 1) 2068 global samples (Lazaridis et al. 2016), 2) the first dataset, 2,504 1000 genome samples (MacArthur et al. 2012), and 471 Ashkenazic Jews (Bray et al. 2010), and 3) the second dataset and 514 ancient DNA samples from (Lazaridis et al. 2016). (Table S1). We used Lazaridis et al.’s (2016) dataset to LD prune the datasets (2068 samples and 621,799 SNPs). After LD pruning using PLINK command (50 10 0.8) and removing SNPs with missingness, allowing no more than five missing SNPs per sample, the datasets included: 230,569, 128,568, and 128,568 autosomal SNPs, respectively.

### Data analyses

All PCA analyses were carried out using Matlab’s (R2020a, Statistics and Machine Learning Toolbox Version 11.7) PCA function, which uses singular value decomposition (SVD), like SmartPCA, and yields nearly identical results to the basic SmartPCA tool (Patterson, Price, and Reich 2006) (Version 7.2.1 without removing outliers, normalization, or projection) (Figure S1-S2).

### Projection of ancient samples

A major challenge in projecting ancient samples onto modern-day samples is handling the high data absences. Lazaridis et al. (2016) addressed this problem using the least-squares projection (lsqproject) implemented in EIGENSOFT. Wang et al. (2015) cautioned that this method does not address the shrinkage problem (where all the projected samples cluster together) and that the results might be misleading. To avoid this problem and the difficulties associated with missing data, in the tenth test case, we analyzed 65 out of 102 of the ancient samples of interest with over 10,000 SNPs in our dataset (with a median of 48,249 SNPs). We then projected one ancient sample at a time, based on the modern-day samples, using only the genotyped SNPs of the former.

### Estimating the citation number of PCA tools

Our estimate that PCA tools were cited over 15,000 is based on the Google Scholar’s citation count for the most commonly used tools using the following searches: “EIGENSTRAT OR EIGENSOFT OR smartPCA” (>7600 times), “SNPRelate” (>1,200 times), “PLINK AND PCA -

EIGENSOFT SNPRelate” (>6700 times), “PCA in R AND genetics” (>400 times), and “FlashPCA OR FlashPCA2” (>200 times).

### Data and scripts availability

All our data and scripts that can replicate our results and figures are available via GitHub: https://github.com/eelhaik/PCA_critique

## Acknowledgment

EE was partially supported by the Crafoord Foundation, the Swedish Research Council (2020-03485), and Erik Philip-Sörensen Foundation (G2020-011). The computations were enabled by resources provided by the Swedish National Infrastructure for Computing (SNIC) at Lund, partially funded by the Swedish Research Council through grant agreement no. 2018-05973.

