## Supplementary materials for "Why most Principal Component Analyses (PCA) in population genetic studies are wrong" (from another version of this preprint: version 3, 2021-05-19)

#### Figure S1

The results of Matlab’s PCA function (pca.m) (A) and SmartPCA (B) are nearly identical when applied to the Lazaridis et al. (2016) ($N_{ind}$=2068) dataset. The axes of (A) were divided by 2500 to match the axes of (B). Populations ($N_{pop}$=189) are colored differently. Selected populations are labeled.

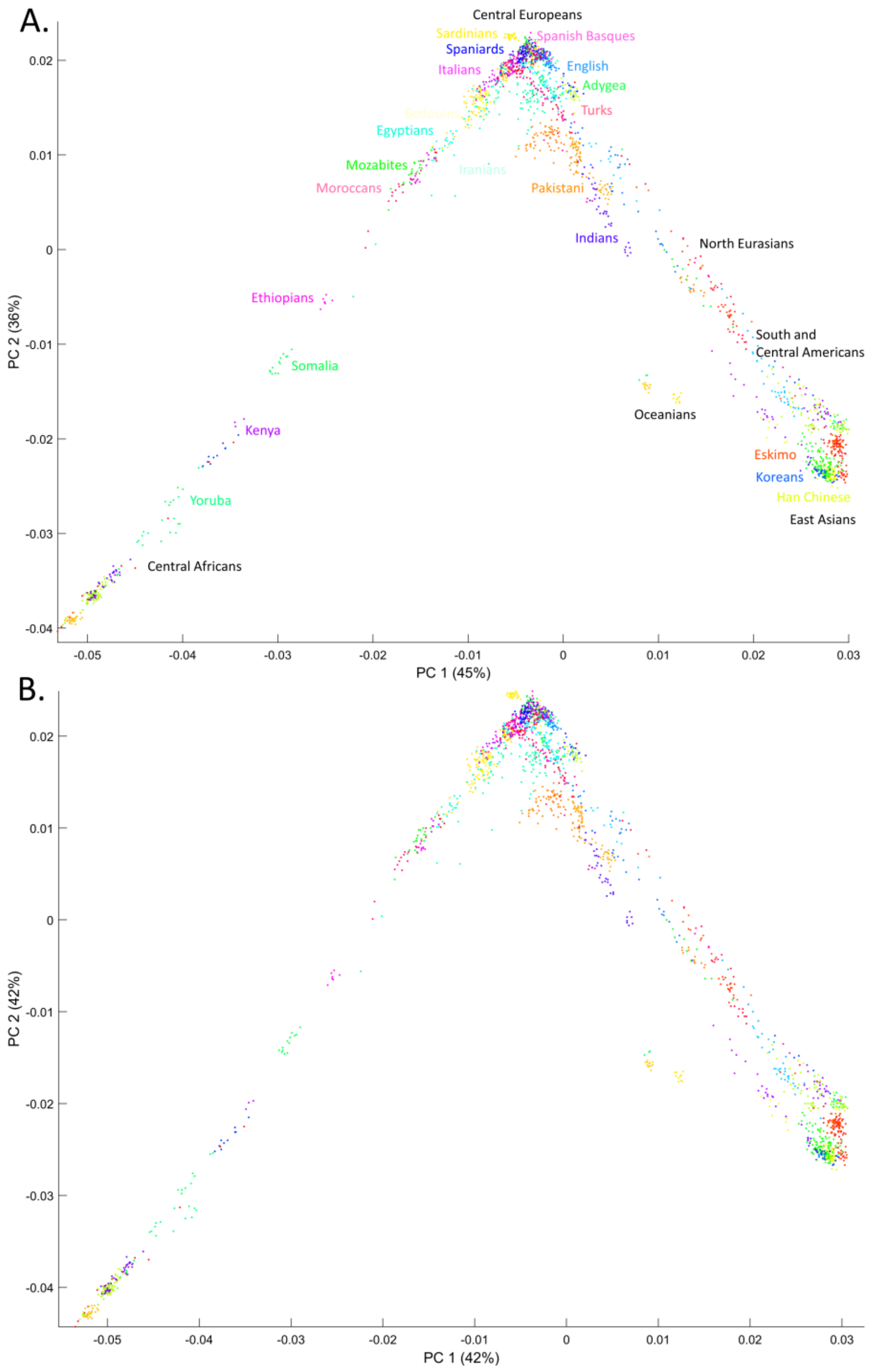

### Figure S2

The first nine principal components (PCs) calculated by Matlab's pca.m (*x-axes*) and SmartPCA (*Y-axes*) are highly correlated when applied to the Lazaridis et al. (2016) ($n_{ind}$=2068) dataset (A-I). Pearson's correlation coefficients range from 0.99 to 1, and the *p*-values are 0 (two-tailed test).

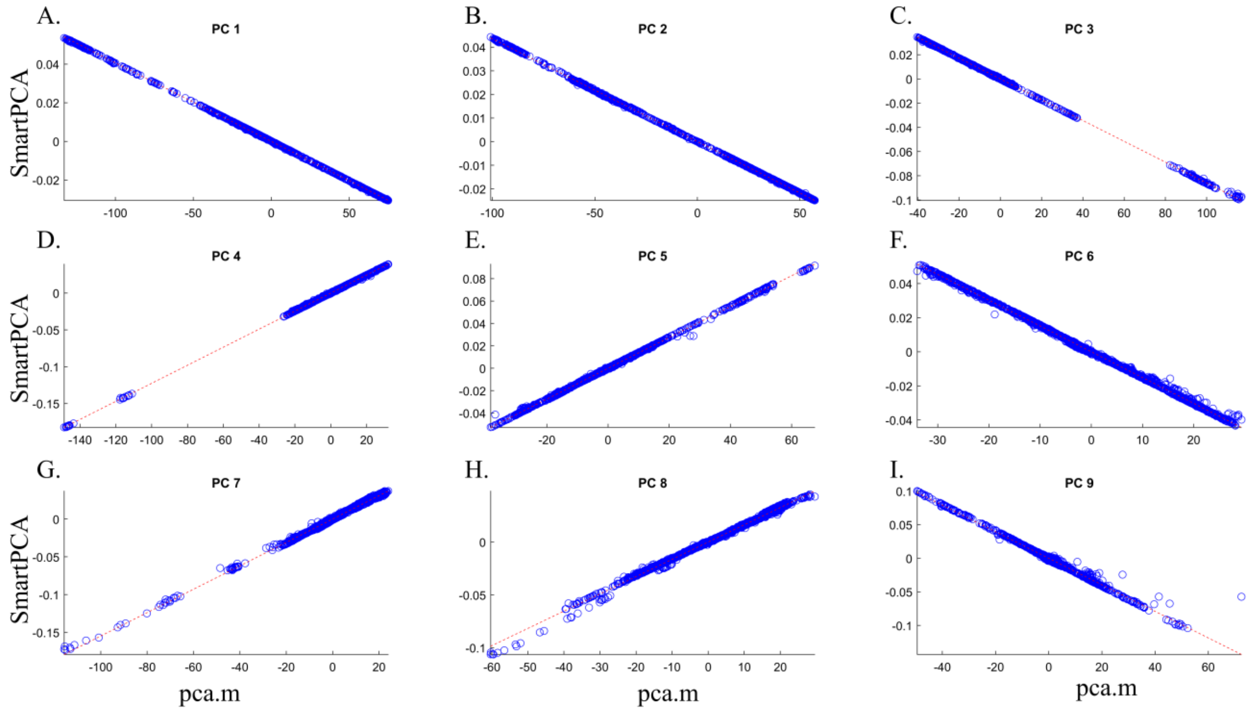

### Figure S3

PCA for Black, the three primary colors, and an admixed Blue-Green color (Cyan). Varying the sample sizes yields different results, such as: A. Black is closer to Blue and Green and Cyan cluster together ($N_{Red}$, $N_{Green}$, $N_{Black}$=100; $N_{Blue}$, $N_{Cyan}$=5), B. Black is closer to Green and Black and Cyan cluster together ($N_{Cyan}$, $N_{Green}$=20; $N_{Red}$=1600; $N_{Blue}$, $N_{Black}$=400), C. Black clusters between Red and Blue, as expected from an admixed color ($N_{Green}$=10; $N_{Black}$=20; $N_{Blue}$=50; $N_{Red}$, $N_{Cyan}$=1000), and D. Black clusters between Red and Green, as expected from an admixed color ($N_{Green}$=50; $N_{Black}$, $N_{Blue}$=10; $N_{Red}$, $N_{Cyan}$=1000);

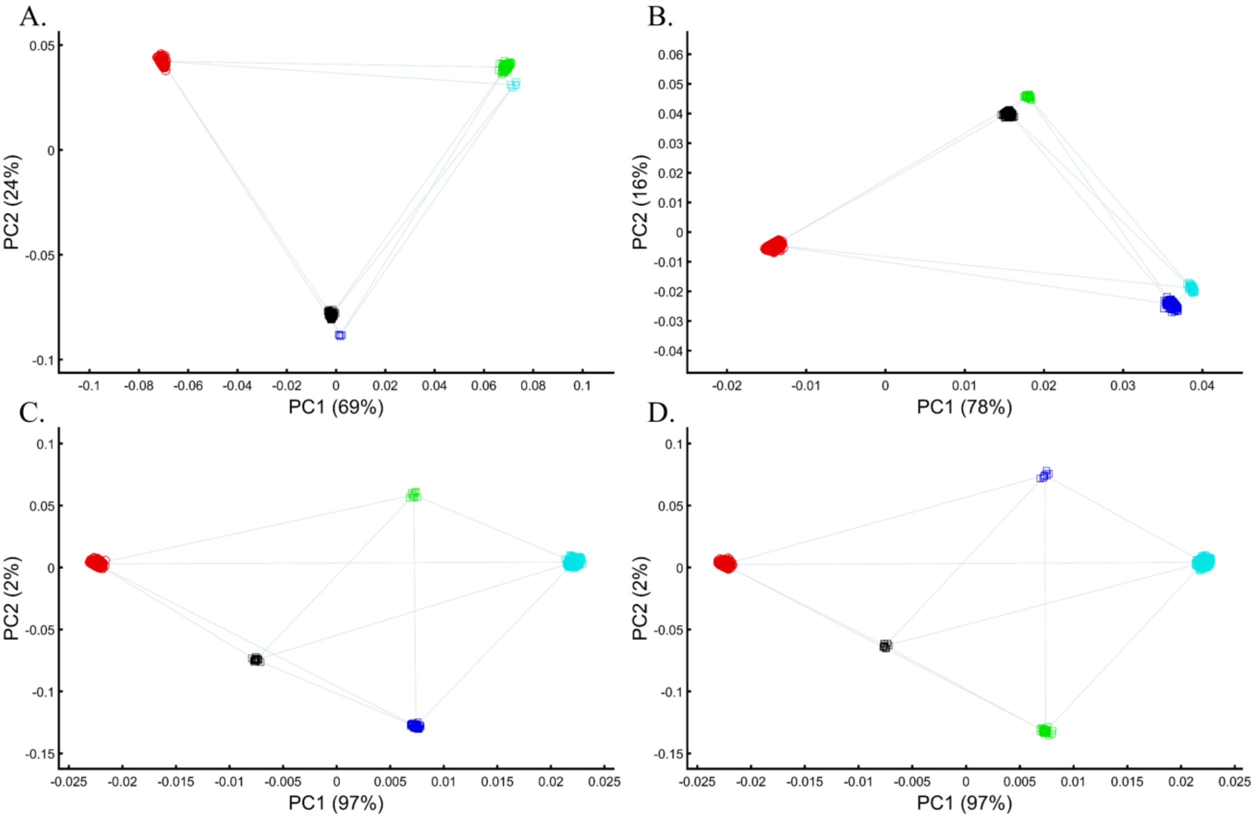

### Figure S4

PCA for present-day populations. The PCA is performed on the same set of individuals as are reported in Figure 24, but without the ancient populations. Whereas in Figure 24, the modern-day populations used to project the ancient samples are greyed (to highlight the ancient samples), here they are colored.

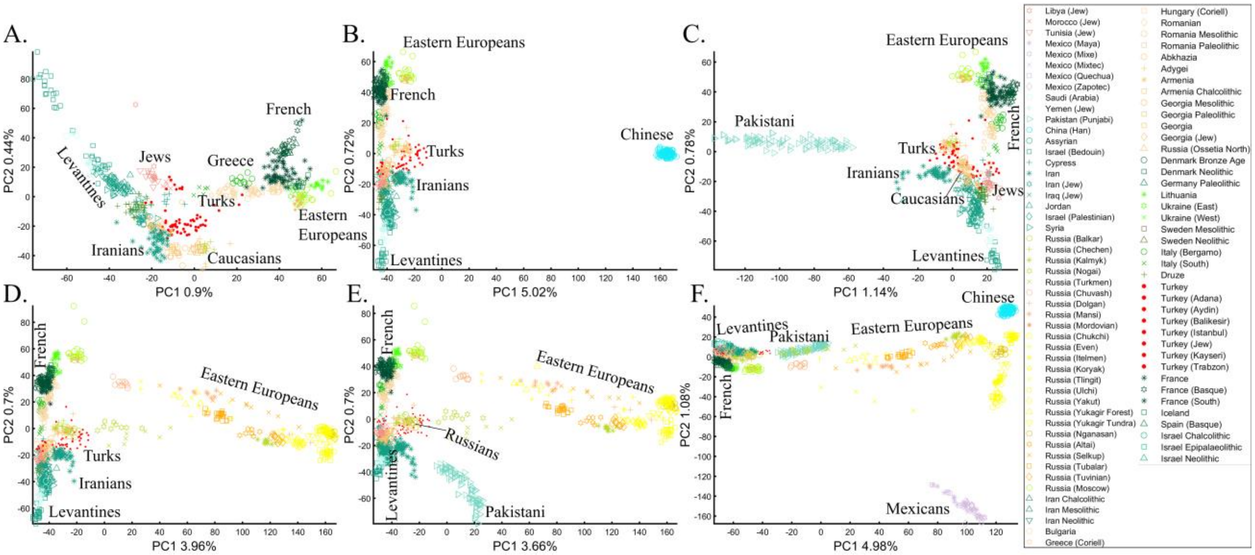

Figure S5

The proportion of explained variance by the first ten principal components (PCs) for a different number of samples (legend) selected at random from the entire dataset.

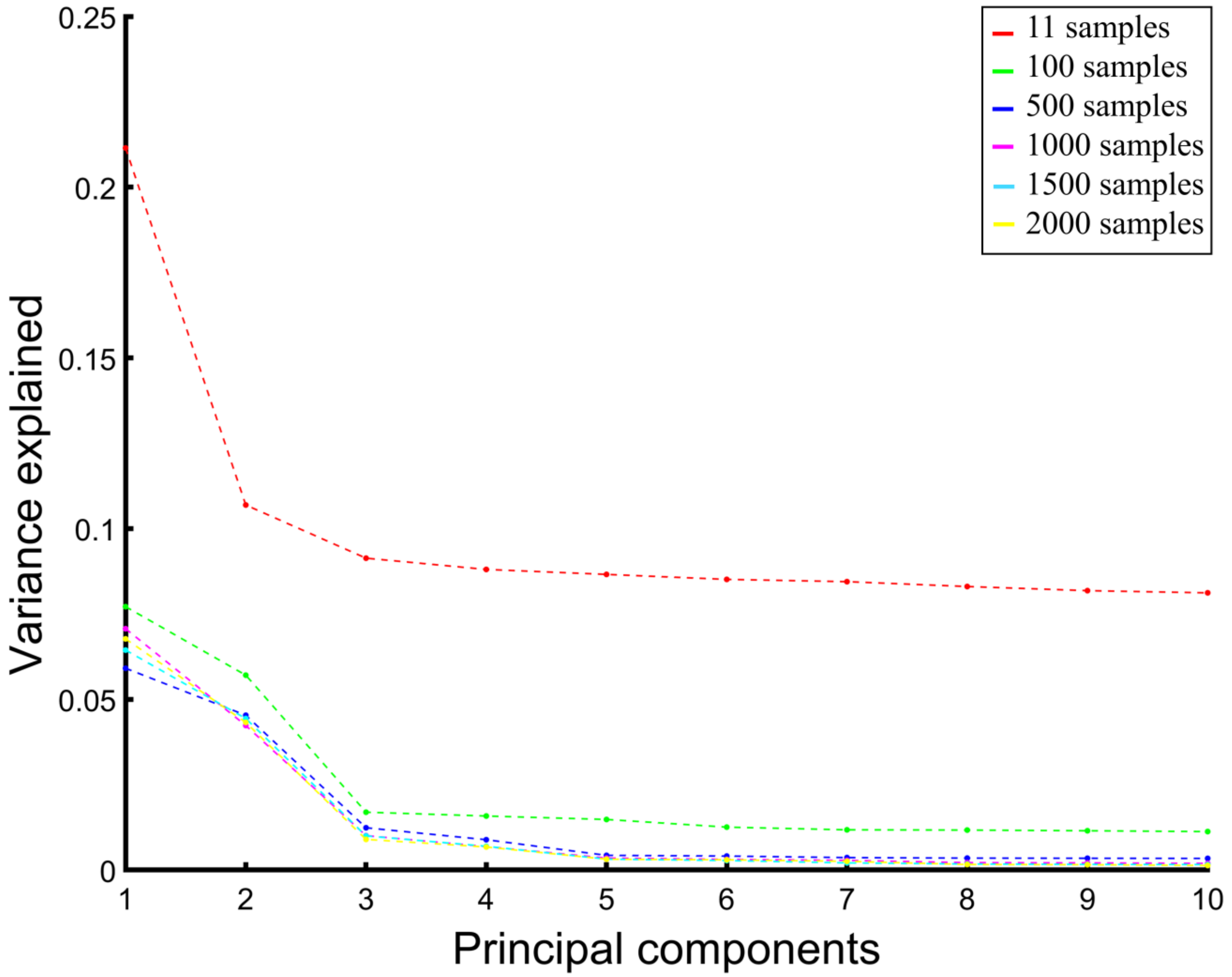

Figure S6

The proportion of variance explained by the first two principal components (PCs) for different numbers of individuals either from the same populations or selected at random from the entire dataset (inset).

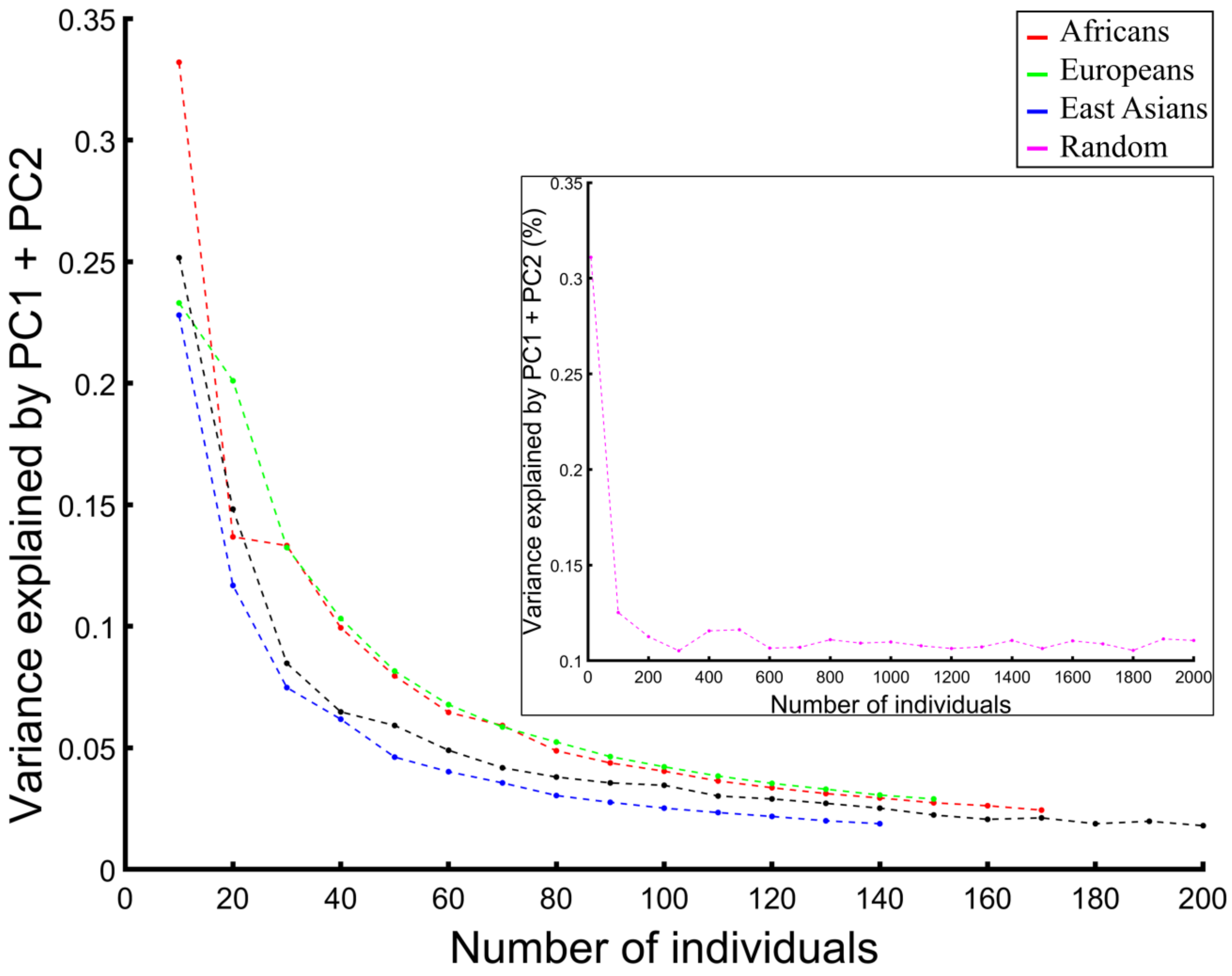

### Figure S7

The proportion of explained variance due to the true structure of the data versus noise. Five 1000 Genomes Populations were selected from four continents: Europeans-Americans (CEU), Telugu Indians (ITU), Vietnamese (KHV), African-Americans (ASW), and Lima Peruvians (PEL). PCA was applied to 1200 random individuals and the reshuffled genetic dataset. We repeated the analyses 50 times. The proportion of variance explained by the first twenty principal components (PCs) (x-axis) is shown in boxplots against the median proportion of variance calculated for the reshuffled matrix.

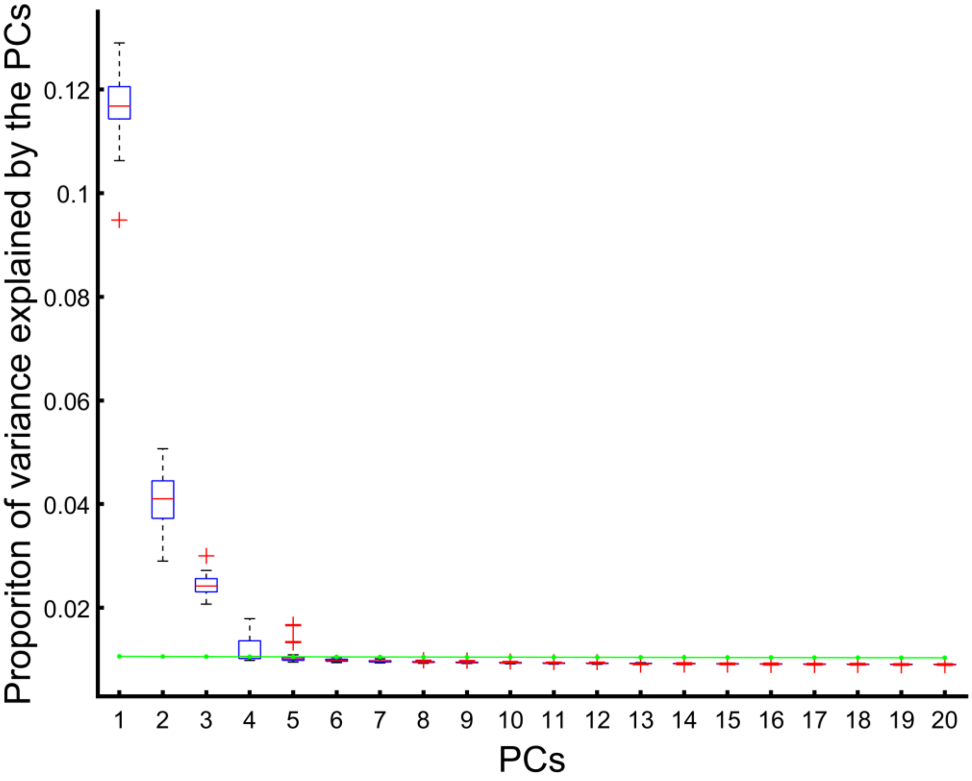

### Figure S8

The effect of dimensionality on sample distances in PCA. Selecting 100 random Yoruba, we calculated the distances between the first two PCs for all individual pairs when applying PCA to 10, 100, 1000, 10,000, or 100,000 SNPs (A). We next calculated the cumulative shortest distance between the first two PCs for every sample (B) and the cumulative median distance for every sample (C) along the same number of SNPs: 10 (red), 100 (green), 1000 (blue), 10,000 (magenta), and 100,000 (cyan). Notice that some of the nearest neighbors analyzed in the highest dimension of 100,000 SNPs (B) have the same distance as the more distanced samples in C).

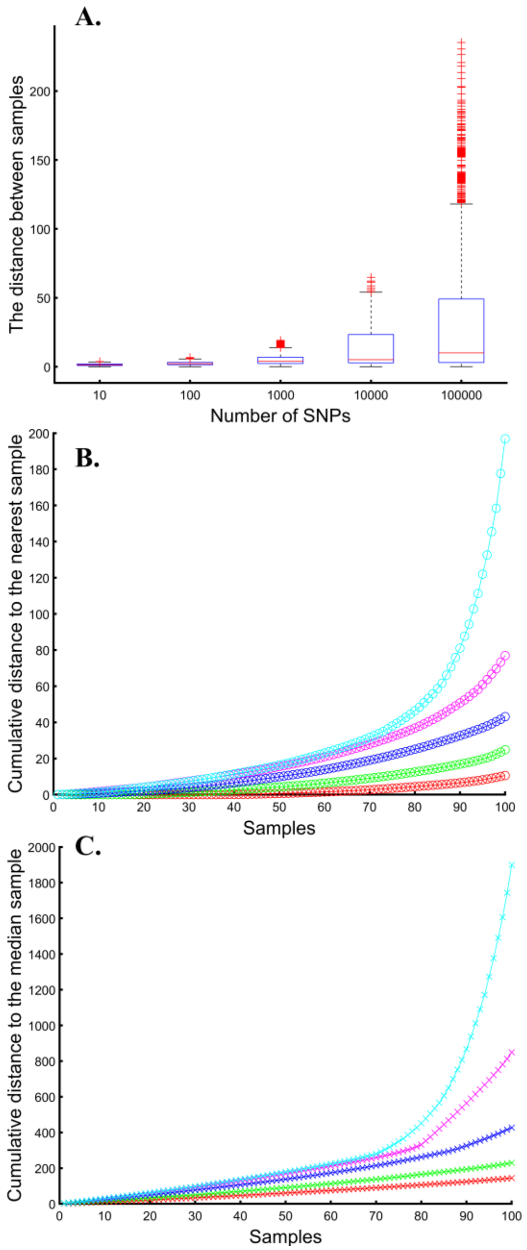

### Figure S9

PCA art: The Eu Bunny. To illustrate the flexibility of PCA, we applied PCA to European populations to form a bunny. European populations (legend) were selected to outline a bunny. Sample sizes were adjusted to avoid cluttering and ensure even distribution of samples. Samples were selected at random from each population. Lines and circles were added manually around population clusters. The code can be found in the GitHub repository of this paper.

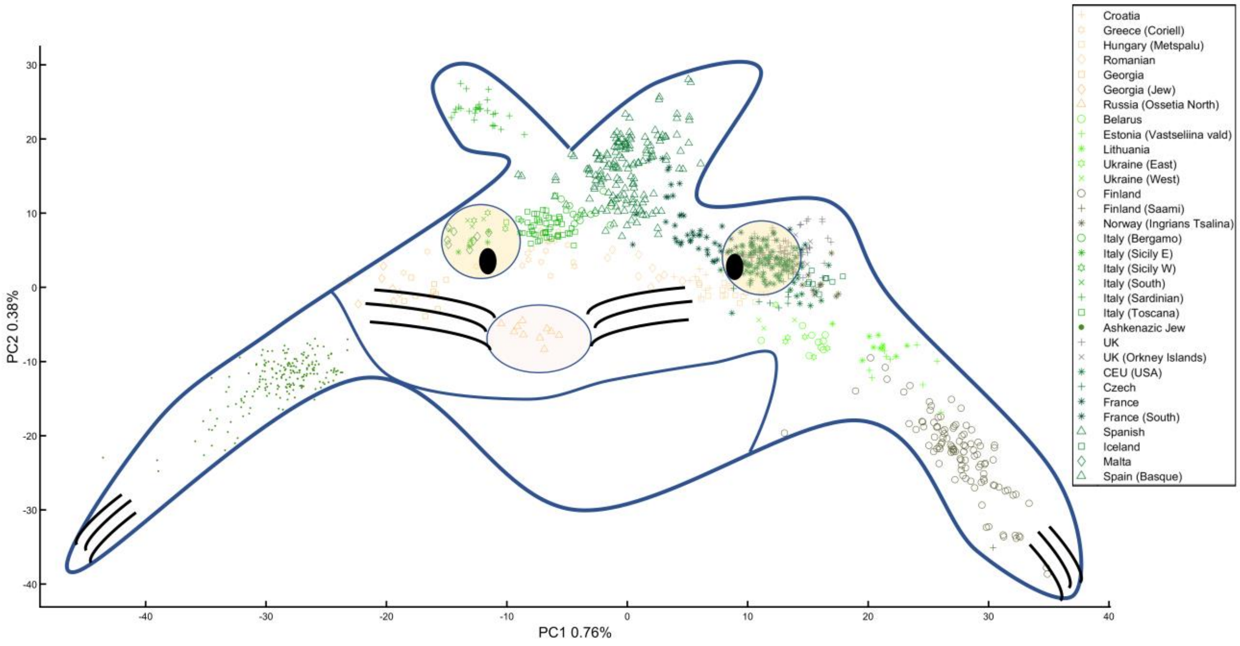

Figure S10

PCA art: An Eu mermaid. To illustrate that PCA plots bear no specific resemblance to geographical regions, we downloaded the PCA map produced by 23andme (https://www.23andme.com/en-int/ancestry-composition-guide/) termed "Principal components plot of 23andMe reference European populations." We added lines around population clusters and regional annotation according to the legend. Contrary to previous claims that PCA plots resemble geographical maps, we found that this map bears no resemblance to Europe's map but may fit the shape of a mermaid. This example illustrates that the alleged resemblance of PCA maps to geographical regions is an artifact of the sampling scheme.

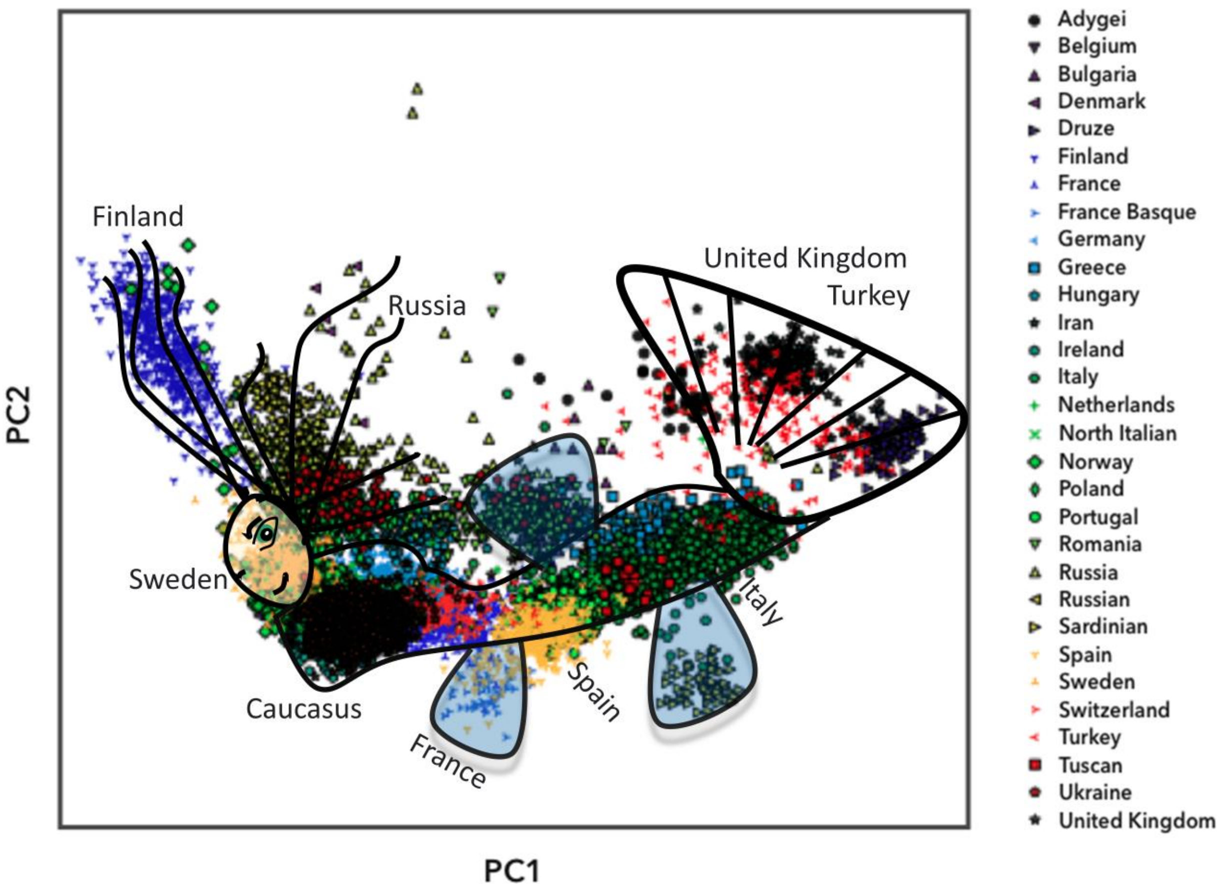

### Table S1

Source of ancient DNA samples used in this study.

| Sample Studied | Country | Publication |
|---|---|---|
| RISE61 | Denmark | (Allentoft, et al. 2015) |
| RISE71 | Denmark | (Allentoft, et al. 2015) |
| RISE94 | Sweden | (Allentoft, et al. 2015) |
| RISE97 | Sweden | (Allentoft, et al. 2015) |
| RISE98 | Sweden | (Allentoft, et al. 2015) |
| WC1 | Iran | (Broushaki, et al. 2016) |
| AH2 | Iran | (Broushaki, et al. 2016) |
| AH1 | Iran | (Broushaki, et al. 2016) |
| AH4 | Iran | (Broushaki, et al. 2016) |
| oase1 | Romania | (Fu, et al. 2015) |
| HohleFels49 | Germany | (Fu, et al. 2016) |
| Falkenstein | Germany | (Fu, et al. 2016) |
| Falkenstein | Germany | (Fu, et al. 2016) |
| SC2 | Romania | (González-Fortes, et al. 2017) |
| SC1 | Romania | (González-Fortes, et al. 2017) |
| I5408 | Romania | (González-Fortes, et al. 2017) |
| OC1 | Romania | (González-Fortes, et al. 2017) |
| sf12 | Sweden | (Günther, et al. 2018) |
| I1179 | Israel | (Harney, et al. 2018) |
| I1152 | Israel | (Harney, et al. 2018) |
| I1166 | Israel | (Harney, et al. 2018) |
| I1178 | Israel | (Harney, et al. 2018) |
| I1165 | Israel | (Harney, et al. 2018) |
| I1160 | Israel | (Harney, et al. 2018) |
| I1184 | Israel | (Harney, et al. 2018) |
| I1170 | Israel | (Harney, et al. 2018) |
| I1164 | Israel | (Harney, et al. 2018) |
| I0644 | Israel | (Harney, et al. 2018) |
| I1182 | Israel | (Harney, et al. 2018) |
| I1169 | Israel | (Harney, et al. 2018) |
| I1171 | Israel | (Harney, et al. 2018) |
| I1168 | Israel | (Harney, et al. 2018) |
| SATP | Georgia | (Jones, et al. 2015) |
| SATP | Georgia | (Jones, et al. 2015) |
| KK1 | Georgia | (Jones, et al. 2015) |
| I0016 | Sweden | (Lazaridis, et al. 2014) |
| I0017 | Sweden | (Lazaridis, et al. 2014) |
| I1632 | Armenia | (Lazaridis, et al. 2016) |
| I1631 | Armenia | (Lazaridis, et al. 2016) |
| I1634 | Armenia | (Lazaridis, et al. 2016) |
| I1407 | Armenia | (Lazaridis, et al. 2016) |
| I1665 | Iran | (Lazaridis, et al. 2016) |
| I1293 | Iran | (Lazaridis, et al. 2016) |

| | | |
|---|---|---|
| I1670 | Iran | (Lazaridis, et al. 2016) |
| I1290 | Iran | (Lazaridis, et al. 2016) |
| I1661 | Iran | (Lazaridis, et al. 2016) |
| I1662 | Iran | (Lazaridis, et al. 2016) |
| I1674 | Iran | (Lazaridis, et al. 2016) |
| I1671 | Iran | (Lazaridis, et al. 2016) |
| I0867 | Israel | (Lazaridis, et al. 2016) |
| I1072 | Israel | (Lazaridis, et al. 2016) |
| I0013 | Sweden | (Mathieson, et al. 2015) |
| I0012 | Sweden | (Mathieson, et al. 2015) |
| I0011 | Sweden | (Mathieson, et al. 2015) |
| I0015 | Sweden | (Mathieson, et al. 2015) |
| I0014 | Sweden | (Mathieson, et al. 2015) |
| I4081 | Romania | (Mathieson, et al. 2018) |
| I4582 | Romania | (Mathieson, et al. 2018) |
| I4607 | Romania | (Mathieson, et al. 2018) |
| I5411 | Romania | (Mathieson, et al. 2018) |
| I5436 | Romania | (Mathieson, et al. 2018) |
| Saxtorp5164 | Sweden | (Mittnik, et al. 2018) |
| Ajvide58 | Sweden | (Skoglund, et al. 2014) |
| Ajvide70 | Sweden | (Skoglund, et al. 2014) |
| Gokhem2 | Sweden | (Skoglund, et al. 2014) |
